# Shifting Resilience: Trends and Predictors of Mesic Resource Productivity in Western U.S. Rangelands

**DOI:** 10.64898/2026.03.27.714799

**Authors:** Kristopher R. Mueller, Scott L. Morford, John S. Kimball, Joseph T. Smith, J. Patrick Donnelly, David E. Naugle

## Abstract

Mesic resources, the late-season herbaceous vegetation found in riparian areas and wet meadows, provide disproportionately important forage and habitat across western U.S. rangelands, yet their response to climatic variability and anthropogenic influences remains poorly understood. Using a 40-year Landsat time series (1984–2024), we quantified trends in late-season productivity (NDVI) across 4.5 million hectares of the sagebrush biome and applied random forest models to distinguish between temporal and spatial predictors of mesic resource productivity. We identified a fundamental shift in how mesic resources respond to drought: from 1984 to 2004, mesic productivity was strongly correlated with drought severity (Palmer Drought Severity Index, R² = 0.92), but this relationship weakened substantially in the next two decades (2005-2024; R² = 0.28), during which time productivity increased despite persistent aridity. Temporal modeling identified rising atmospheric CO_2_ concentrations as the strongest predictor of this shift, consistent with enhanced plant water-use efficiency under CO_2_ fertilization. Spatially, large agricultural valley floodplains act as anthropogenic refugia, sustaining productive mesic resources through flood irrigation and subsequent groundwater recharge into late summer. These findings suggest that human water management and physiological shifts in vegetation are currently buffering mesic systems against meteorological drought throughout U.S. rangelands. However, this apparent buffering is spatially heterogeneous and may mask vulnerability to groundwater depletion, shifts in precipitation regimes, and woody encroachment. Sustaining these vital ecosystems will require conservation approaches that go beyond climate monitoring to include balanced management considering both agricultural and ecological water needs and constraints.

## 1 Introduction

Mesic ecosystems are vital to maintaining biodiversity and ecosystem functions in the arid and semi-arid rangelands worldwide (Grenfell et al., 2022; W. D. Williams, 1999). In these dry environments, riparian zones and wetlands - called mesic hereafter - support a disproportionately large share of wildlife, livestock, and vegetation, given their small geographic area. Within the western United States (U.S.), these mesic environments cover only 2% of rangelands, yet they provide essential resources to 80% of terrestrial wildlife during late summer and fall, when water becomes scarce (Maestas et al., 2023; Naiman et al., 1993; Thomas et al., 1979). Herbaceous vegetation in these ecosystems, termed mesic resources, is especially valuable. It offers highly productive forage that remains lush as adjacent upland soils dry out. Because these resources are typically phreatophytic, relying on access to persistent groundwater, they are highly sensitive to fluctuating water tables and serve as critical indicators of water availability in landscapes increasingly impacted by drought (Kolarik et al., 2023; Manning et al., 2020; Noy-Meir, 1973; Shrestha et al., 2024). As hydrological patterns change, understanding the factors that impact mesic resource productivity is essential for effective land management, though the nature of these climate-driven shifts remains uncertain.

In the western U.S., climate change threatens to alter water availability, causing complex shifts in the function of mesic ecosystems through more frequent and severe droughts and altered precipitation regimes (Dettinger et al., 2015; Polley et al., 2013). Regional forecasts reveal diverging patterns: the southwestern U.S. is moving toward warmer, drier conditions and earlier water shortages, whereas projections for the Northern Great Plains indicate potential increases in herbaceous forage due to more precipitation, higher temperatures, and higher atmospheric carbon dioxide (CO_2_) levels (Lilley et al., 2001; Morgan et al., 2011; Polley et al., 2013; Read et al., 1997). Compounding these climatic shifts, land management practices, particularly irrigation, strongly influence the availability of these water resources in large valley floodplains (Donnelly, Jensco, et al., 2024). Irrigation in the 17 Western states accounts for 74% of the country’s freshwater withdrawals (Dieter et al., 2018). Certain practices, notably flood irrigation, support mesic resources by increasing groundwater recharge and mimicking historical floodplain hydrology, ultimately sustaining more than half of all wetlands in the West (Donnelly, Jensco, et al., 2024; Essaid & Caldwell, 2017; Gordon et al., 2020). These predominantly privately owned working lands rely heavily on flood irrigation for livestock forage and hold 75% of available mesic resources (Donnelly et al., 2016). As water restrictions tighten across the arid West, the continued reliance on flood irrigation raises a critical question: Does human intervention play a greater role in maintaining late-season mesic resources than climate-driven drought in limiting them?

The continuous 40-year archive of Landsat satellite imagery enables scientists and land managers to evaluate these complex landscape processes across unprecedented spatial and temporal scales (Wulder et al., 2022). Annual vegetation mapping across the U.S. has already helped identify intact sagebrush ecosystems and enhance conservation efforts for the greater sage-grouse (*Centrocercus urophasianus*) and other sagebrush-dependent wildlife (Doherty et al., 2022; Kumar et al., 2024; Naugle et al., 2024). Although researchers have documented mesic resource responses to overgrazing, historical land cover changes, and localized drought (C. M. Albano et al., 2020; Kauffman et al., 2022; Macfarlane et al., 2017), the response of mesic productivity to environmental and human water use remains poorly understood at larger scales (Donnelly et al., 2018; Lundblad et al., 2022; Silverman et al., 2019). To address this gap, this study leverages the Landsat archive (1984–2024) to identify late-season mesic resources and track their productivity trends in relation to meteorological drought. By applying random forest modeling to evaluate a suite of abiotic environmental variables—including climate, surface water availability, and geomorphology—we distinguish the primary temporal and spatial predictors of mesic productivity. Deepening our understanding of how mesic resources respond to shifting climate and land-use drivers will provide land managers with targeted insights to better prioritize conservation and restoration, ultimately securing late-season mesic resources for wildlife and livestock when they are needed most.

## 2 Methods

### 2.1 Study Area

The study area covers the semi-arid sagebrush biome of the western U.S. We used EPA Level III Ecoregions (US EPA, 2015) to subdivide the study area based on dominant ecological processes and precipitation patterns (Fig. 1). Mountain and basin regions receive most precipitation in winter, leading to spring and early summer runoff. In contrast, the Northern Great Plains experience short, intense summer storms and fluctuating soil moisture during the growing season (Barker & Whitman, 1988; Lauenroth et al., 2014). Differences in dominant vegetation types (e.g., shrublands, forests, and grasslands) distinguish these regions and reflect ecosystem variation.

**Figure 1.**
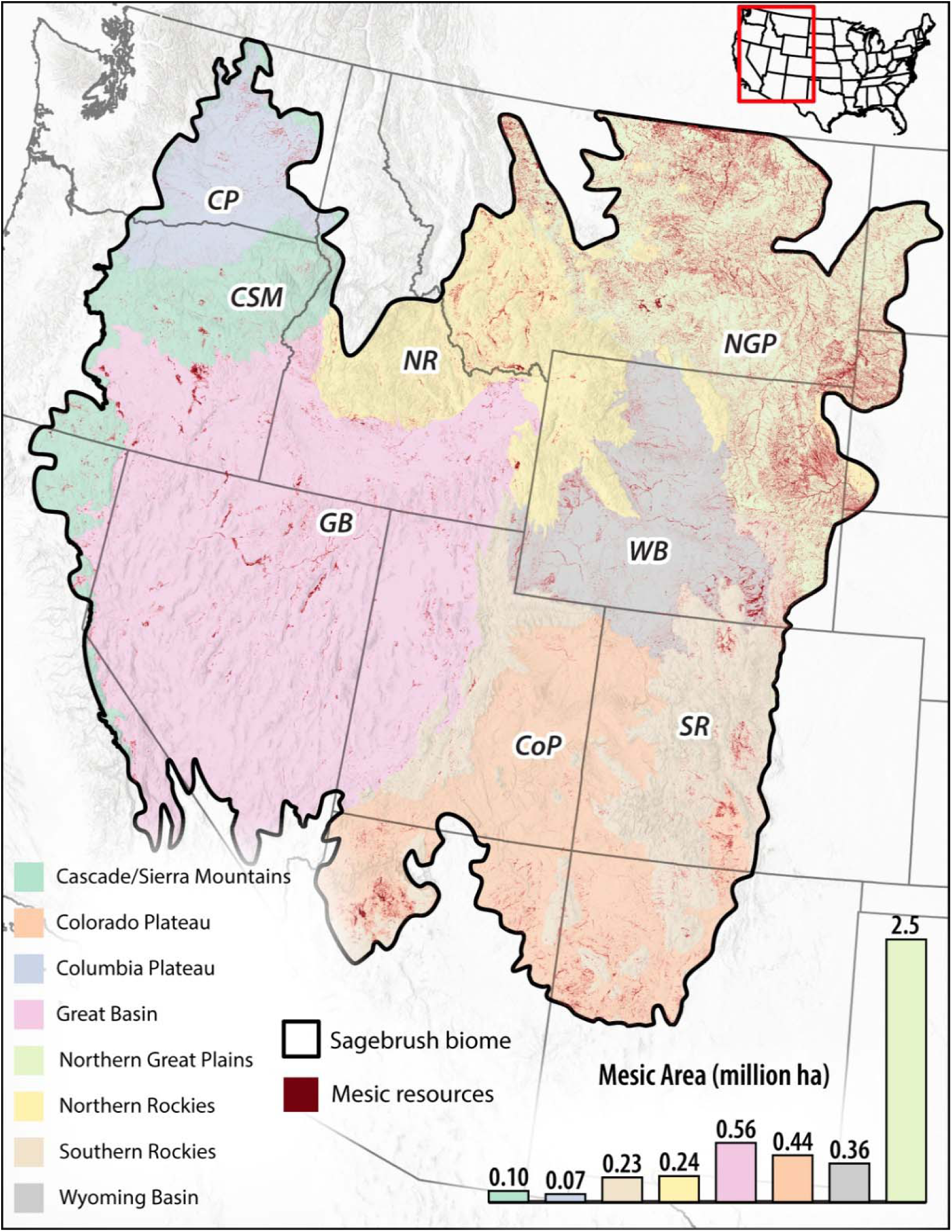
The study area (sagebrush biome) displays the ecoregions used in the analysis. The general climate and landscape vegetation include cold deserts (blue, pink, grey, and orange), forested mountains (teal, yellow, tan), and grassland plains (green). Red shading shows the final mesic mask used for sampling in the analysis (see Section 2.2). The area of identified mesic resources per ecoregion is measured in millions of hectares.

### 2.2 Identifying Mesic Resources

We used Google Earth Engine (Gorelick et al., 2017) to analyze Landsat 5, 7, 8, and 9 atmospherically corrected surface reflectance (Collection 2, Tier 1) imagery (1984–2024), focusing on the late growing season (July 15–September 30), when moisture stress limits herbaceous growth and mesic areas persist due to shallow groundwater (Vicente-Serrano et al., 2013). The QA_PIXEL band removed clouds, shadows, snow, and water. For each year, we calculated mean NDVI (Normalized Difference Vegetation Index) at 30-meter resolution as a proxy for late-season productivity and an indicator of water availability (Kolarik et al., 2023; Yengoh et al., 2015) (Fig. 2A). We classified pixels as potential mesic resources if they had at least one year with a mean NDVI ≥ 0.3, a standard threshold for productive mesic resources in western North America (Donnelly et al., 2016, 2018; Lundblad et al., 2022; Weier & Herring, 2000). We further masked out cultivated lands, urban areas, high-elevation rocky terrain, forests/woodlands, and pixels that had significant tree or shrub cover change (Fig. 2B; Appendix A, Tables A.1-A.2). To identify only wetlands and riparian floodplains we used the USDA Cropland Data Layer (2023) and LANDFIRE Biophysical Setting (2022), and the Valley-Bottom Extraction Tool (VBET) (Gilbert et al., 2016). The final mask (Appendix A, Fig. A.1) was used to sample pixels and to calculate the mean late-season NDVI for subsequent analyses.

**Figure 2.**
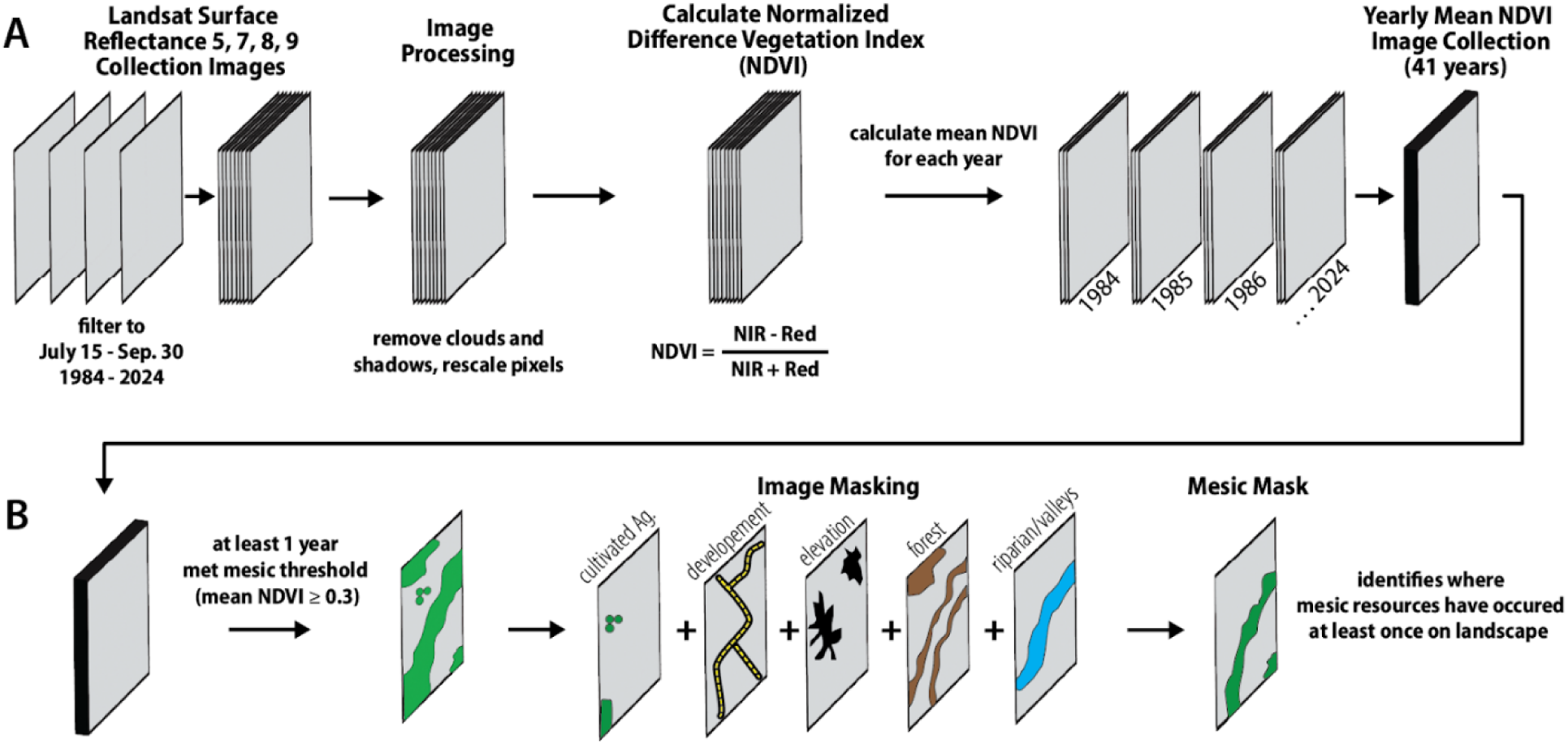
Image processing workflow within Google Earth Engine. Panel A illustrates the process for calculating yearly mean productivity during the late growing season (July 15 - September 30). Panel B demonstrates the method for masking out pixels not associated with mesic resources within the sagebrush biome.

### 2.3 Measuring Trends

We divided the 41-year time series into P1 (1984-2004) and P2 (2005-2024) to compare mesic resource dynamics and identify temporal changes in productivity. This split aligns with the onset of intensifying megadrought across the American West (A. P. Williams et al., 2020), enabling assessment of vegetation responses before and during heightened aridification. We excluded 2012 data due to limited Landsat coverage. To evaluate whether mesic productivity related to drought varies across periods, we calculated correlation coefficients (R²) between NDVI and the mean Palmer Drought Severity Index (PDSI) for each period. We aggregated mesic pixels to determine the mean annual late-season NDVI and PDSI using a sub-monthly drought index (Rhee & Carbone, 2007), and weighted these means by ecoregion size. We further analyzed ecoregional trends in mesic resource productivity using Mann-Kendall (MK) tests, which detect monotonic trends over time (Cliff & Charlin, 1991; *IBM Surveillance Insight for Financial Services 2.0.2*, 2021), yielding the Tau-b correlation coefficient (*τ_b_*). *τ_b_* values, ranging from −1 (negative trend) to 1 (positive trend), were computed for each mesic pixel using Google Earth Engine’s *ee.Reducer.kendallsCorrelation* (Knight, 1966), with values near 0 indicating minimal trends (Beal, 2016) (Appendix A, Fig. A.2). τ_b_ values were calculated for each period by randomly sampling 200,000 points from the mesic resource mask in each ecoregion. Box plots visualized trend distributions across regions and periods. Due to the large sample sizes involved in this spatial analysis, we examined the distribution of MK *τ_b_* effect sizes to identify trends, since standard pixel-level significance tests can be too sensitive in extensive remote sensing datasets.

### 2.4 Random Forest Modeling

We used random forests (RFs) to identify key variables for predicting late-season mesic resource productivity across the biome and within ecoregions. RFs are robust, nonparametric algorithms that are resistant to overfitting (Breiman, 2001), making them ideal for analyzing complex land-use and land-cover datasets (Lundblad et al., 2022; Menuz & Kettenring, 2013; Phan et al., 2020; Wang et al., 2022). We ranked variables by importance (VIMP) and used partial dependence plots (PDPs) to show the direction, shape, and magnitude of each variable’s effect on productivity, aiding interpretation of these “black-box” models (Friedman, 2001). Biome-wide models used a stratified random sample of 12,000 pixels, weighted by ecoregion area (Table 1). Each ecoregion model used 10,000 sampled pixels. The years 1984, 1985, 2012, and 2024 were excluded to align with the covariates’ temporal resolutions. To isolate temporal versus spatial influences on mesic productivity, we applied two data transformations before modeling:

1. Drivers of temporal change: Divide data into P1 and P2 and transform each annual observation into a late-growing-season anomaly by subtracting its location- and period-specific mean (Enders & Tofighi, 2007).
2. Drivers of spatial variation: Average over the full 1986-2023 time series at each sample location to identify spatial patterns.

**Table 1.** Sample sizes per ecoregion and area weights for the biome models.

| **Ecoregion** | **Biome Model**<br>**Total Samples** | **Sample**<br>**Weights** |
| --- | --- | --- |
| Cascade/Sierra Mountains | 816 | 0.068 |
| Colorado Plateau | 1,630 | 0.136 |
| Columbia Plateau | 511 | 0.043 |
| Great Basin | 3,615 | 0.301 |
| Northern Great Plains | 1,998 | 0.166 |
| Northern Rockies | 1,215 | 0.101 |
| Southern Rockies | 1,310 | 0.109 |
| Wyoming Basin | 905 | 0.075 |
| **Total samples per year** | 12,000 |  |
| **Years in Model (P1)** | 19 |  |
| **Years in Model (P2)** | 18 |  |

RF models were constructed using the *randomforestSRC* package (version 3.3.1) in R with default parameters (ntree = 1000 and importance = “permute”). Predictors included 17 abiotic environmental variables (Table 2). We excluded vegetation functional cover types (e.g., perennial grass, shrub cover, etc.) to avoid circularity, since these are derived from the same spectral imagery as our response variable (NDVI) (Jones et al., 2018). Static variables were excluded from temporal driver models because they do not vary over time (Table 2). VIMP scores ranked the most influential predictors of each model (Breiman, 2001). PDPs illustrated the marginal effect of each predictor on NDVI, aiding interpretation (Friedman, 2001). VIMP scores were calculated from the final models using subsampling (n=100) with the Breiman-Cutler permutation method and 95% confidence intervals via a delete-*d* jackknife estimator (Ishwaran & Lu, 2019; Politis & Romano, 1994; Shao & Wu, 1989). Out-of-bag (OOB) predictions and errors were used to assess model accuracy by measuring explained variance (R^2^), similar to cross-validation (Hastie et al., 2001).

**Table 2.** Covariates used in random forest modeling. Unless otherwise specified, all variables represent average values during the late growing season (July 15 - September 30). Density metrics indicate total hectares within 500 meters of each sample point. The table includes the data source, data spatial resolution, and ecological rationale for each covariate.

| Covariate | Dataset/Resolution | Reasoning |
| --- | --- | --- |
| **Irrigated agriculture density** | (Ketchum et al., 2020)<br>30-meter | Flood irrigation can recharge groundwater and raise the water table |
| **Annual watershed snow water equivalent**<br>**Max. temperature**<br>**Precipitation**<br>**Winter max. Temperature (Dec. - Feb.)**<br>**Spring precipitation** | DAYMET<br>(Thornton et al., 2022)<br>1000-meter | Climatic variables account for how much water is input into landscapes and for how long |
| **Temporary wetlands (Static; 1984 - 2023)**<br>**Seasonal wetlands (Static; 1984 - 2023)**<br>**Semi-permanent wetlands (Static; 1984 - 2023)** | (Donnelly, Collins, et al., 2024)<br>30-meter | The amount of water on the landscape and its duration dictate water table depth and availability for mesic resources |
| **Frost-free days** | GRIDMET<br>(Abatzoglou, 2013)<br>Meteorological Data<br>4600-meter | Equivalent to the potential growing season length |
| **Elevation (Static)**<br>**Slope (Static)** | U.S. Geological Survey,<br>3D Elevation Program 1-Meter Resolution Digital Elevation Model.<br>1-meter | Geomorphological effects on plant distribution and elevation affect the productivity of mesic resources |
| **Annual global CO<sub>2</sub> (ppm)** | Global Average CO<sub>2</sub><br>(Lan et al., 2026) | CO<sub>2</sub> fertilization accounts for up to 44% of the global GPP increase since the early 2000s |
| **Annual watershed runoff**<br>**Watershed runoff** | USGS WaterWatch<br>(*USGS Water Data for the Nation*) | Amount of water flow calculated at stream gauges at the outputs of watersheds |
| **Palmer Drought Severity Index (PDSI)**<br>**Palmer Z index** | GRIDMET<br>(Abatzoglou, 2013)<br>Drought Indices<br>4600-meter | Drought severity is a key indicator of water availability in soil; these two metrics use different time intervals to account for long-term and short-term cycles |

## 3 Results

### 3.1 Mesic Resource Productivity Trends

Mesic resources within the sagebrush biome covered 4.5 million hectares (2.5% of the study area; Fig. 1). Most identified mesic resources occurred in the Northern Great Plains (2.5 million ha), while other ecoregions had between 74,000 ha (Columbia Plateau) and 556,000 ha (Great Basin). This distribution reflects ecoregion size and climate, with drier basins and plateaus supporting fewer mesic resources. Over the past two decades, late-season productivity (NDVI) and climate aridity (PDSI) have decoupled. From 1984–2004 (P1), late-season mesic productivity closely tracked climate aridity, with mean PDSI accounting for 92% of NDVI variation (p < 0.001) and both variables declining over time (Fig. 3A–C). From 2005–2024 (P2), mean NDVI increased while PDSI remained steady, and the correlation weakened; PDSI explained only 28% of NDVI variation (p < 0.05). Across all eight ecoregions, productivity declined or remained stable during P1 and rose in P2 (Appendix A, Fig. A.3). In P2, productivity increased in all ecoregions, with more than 75% of sampled pixels showing upward trends, especially in the Northern Rockies and Wyoming Basin. NDVI and PDSI relationship figures for all ecoregions are in Appendix A; Fig. A.7-A.14).

**Figure 3.**
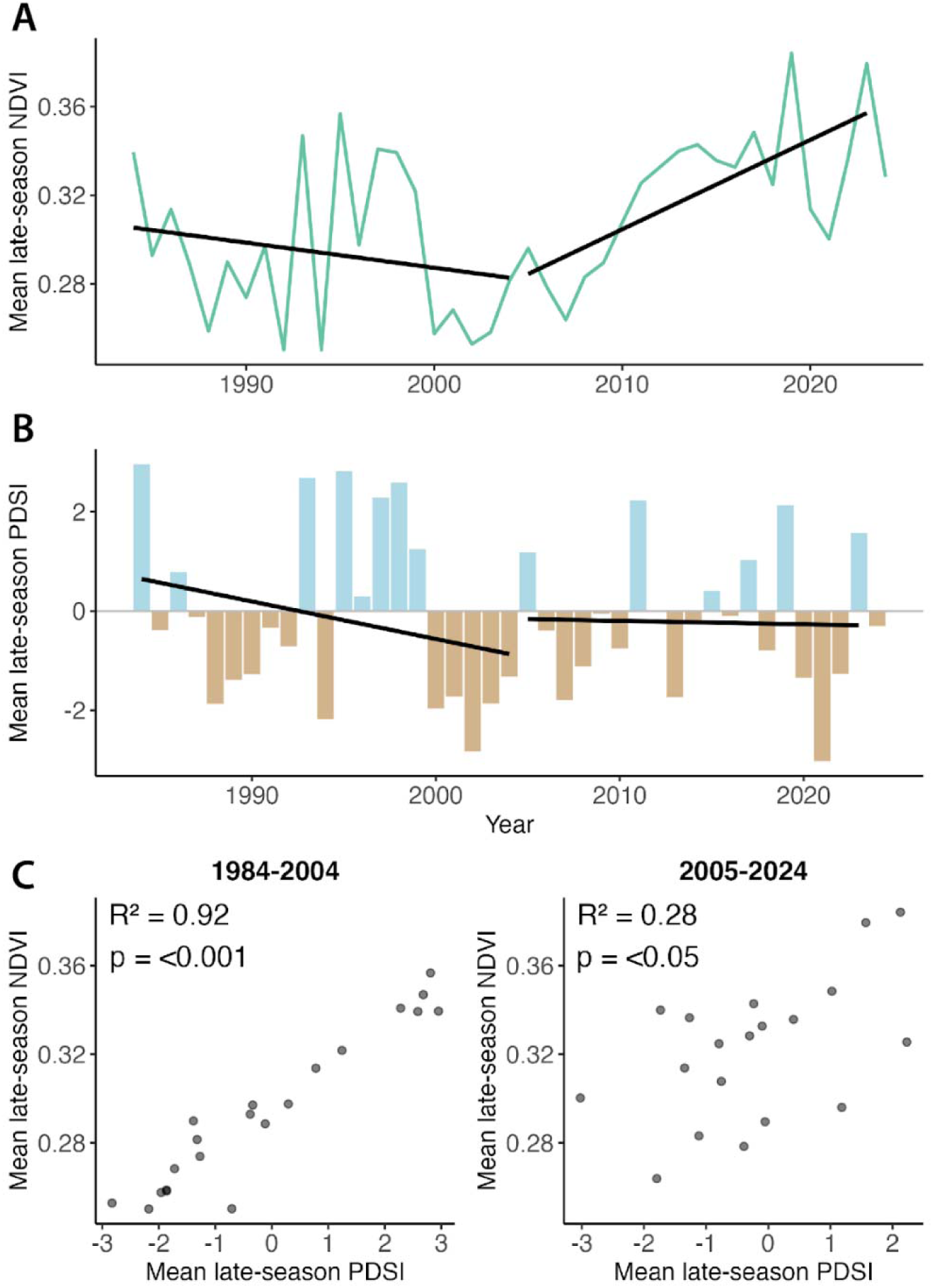
The relationship between mean late-season (July 15 - September 30) mesic resource productivity and drought conditions across the Sagebrush biome, 1984-2024. **A**: Time series of Normalized Difference Vegetation Index (NDVI; **B**) and Palmer Drought Severity Index (PDSI; B) show diverging trends between period 1 (1984-2004) and period 2 (2005-2024). Black lines indicate linear regression trends for each period. **C**: The correlation between NDVI and PDSI reveals a strong relationship during 1984-2004 (R² = 0.92, p < 0.001) that weakens substantially during 2005-2024 (R² = 0.28, p < 0.05).

### 3.2 Predictors of Mesic Resource Productivity

Random Forest models helped explain late-season mesic productivity variation across time and space (Appendix A, Table A.3). Biome-wide, temporal models accounted for 54% (P1) and 57% (P2) of the variation in mesic productivity (OOB error: 0.002). The biome-wide spatial model explained 65% of NDVI variation (OOB error: 0.005). Model performance varied by ecoregion; temporal models performed best during P2 (notably in the Northern Great Plains), while spatial models generally outperformed temporal models, except in smaller ecoregions such as the Columbia Plateau.

Biome-wide temporal analysis revealed shifts in drivers of interannual variation between periods. During P1 (1986–2004), late-season PDSI was the dominant predictor of late-season productivity across the biome (Fig. 4). In P2 (2005–2023), CO_2_ became the most important predictor. Other variables, including late-season Palmer Z-index, precipitation, and late-season temperatures, were still important in model predictions, though consistently less influential than during P1. Partial dependence plots confirm strong effects of PDSI and CO_2_ on mesic productivity in P2 (Fig. 5); drought (low PDSI value) was associated with lower productivity in both periods. CO_2_ had a minimal effect in P1 but a major influence in P2, similar in magnitude to PDSI. Irrigation density, while not a key temporal predictor, was strongly associated with higher productivity in the partial dependence plots for both periods.

**Figure 4.**
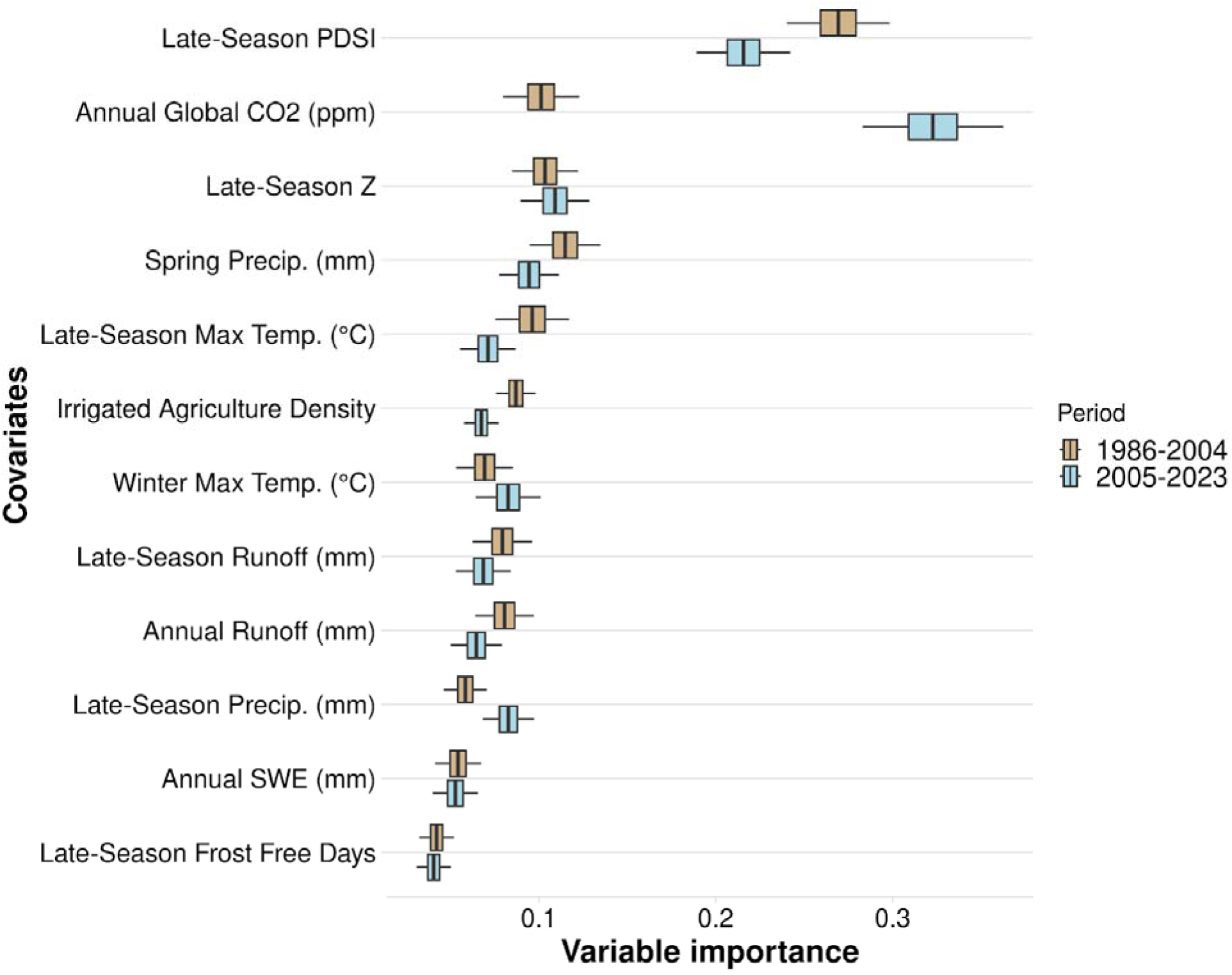
Variable importance (VIMP) of variables predicting mesic resource productivity across the sagebrush biome during 1984-2004 (P1) and 2005-2024 (P2) using the temporal random forest analysis. Variable importance was assessed using the Breiman-Cutler permutation method, with higher values indicating greater predictive power. Center black lines represent median importance values from 100 subsampled VIMP scores; boxes cover the 25th to 75th percentiles; whiskers extend to 95% confidence intervals. Variable importance is standardized by dividing by the variance of Y. Variables are ranked by their mean importance across both periods. Variables shown as densities are measured in hectares per 500-meter radius.

**Figure 5.**
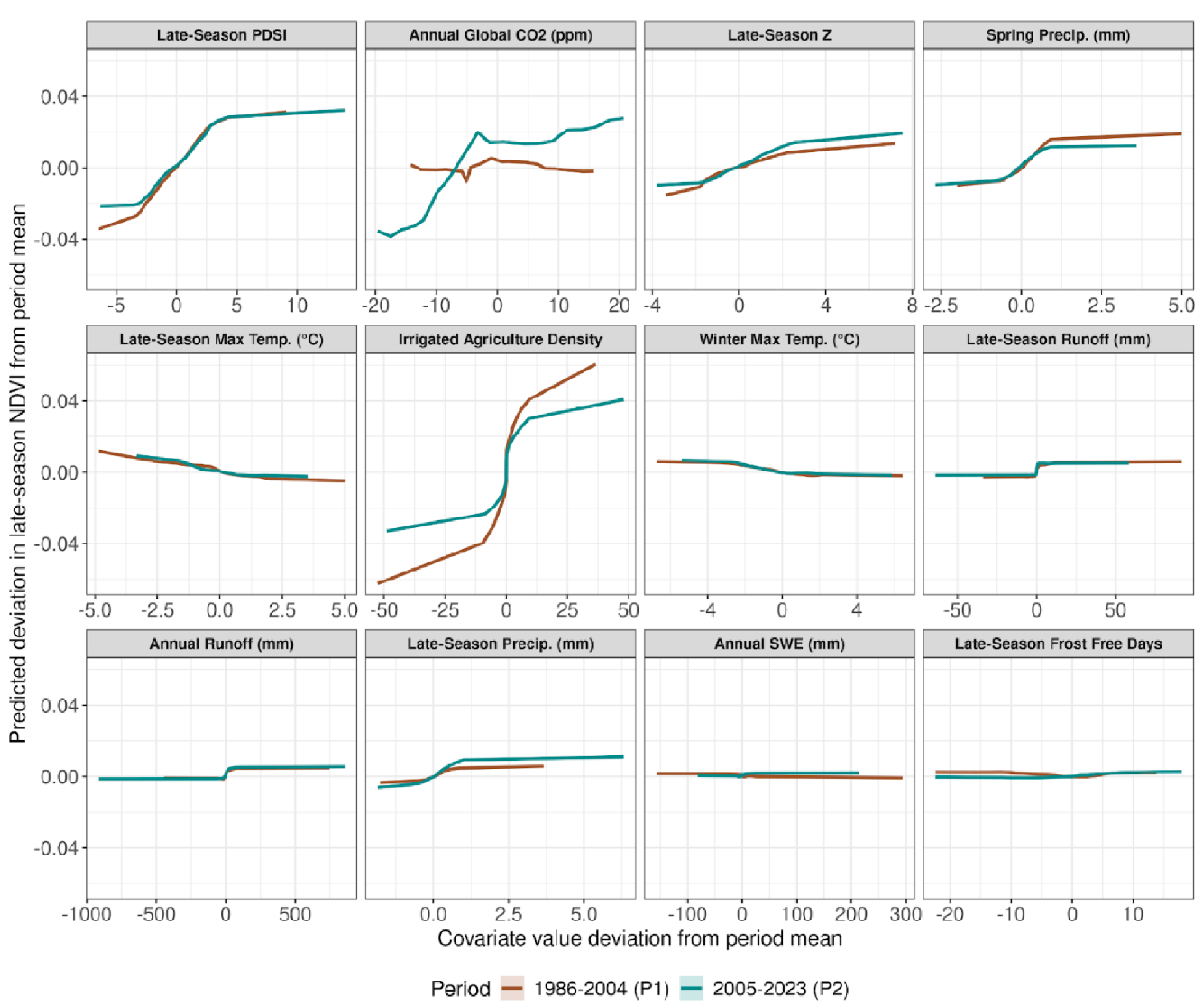
Partial dependency plots for each variable used in the temporal model on predicting mesic resource productivity in the sagebrush biome. Y-axis values are the predicted deviation of late-season NDVI from the period mean, and the X-axis shows variable deviation from its period mean. Values below 0 indicate below-average values for the time period, and values above 0 are above-average values for both the Y and X axes. Different bar colors for P1 (brown; 1986-2004) and P2 (blue; 2005-2023) indicate differences in the effects of variables on mesic productivity over time. Variables shown as densities are measured in hectares per 500-meter radius.

Partial dependence plots indicated that the temporal effects of predictors were generally consistent in direction across periods, except for global CO_2_, which showed a positive relationship only in P2. The magnitude of change between periods differed most for PDSI, irrigated agriculture density, and CO_2_. More severe droughts in P1 were associated with lower productivity, a trend that persisted in P2, albeit with higher productivity at the same drought severity level. Similarly, the effect of irrigation density on predicted NDVI was weakened in P2. While differences in the partial effect of the predictors between P1 and P2 were consistent across ecoregions, the order and magnitude of importance varied. VIMP scores and PDP figures for each ecoregion’s temporal model are shown in Appendix B, Figures B.1-B.16.

Spatial RF variable importances showed that irrigated agriculture density was the primary predictor of mesic productivity at the biome level (Fig. 6). Productivity increased sharply with irrigation density (Fig. 7). Other important predictors included late-season maximum temperature (negatively associated with productivity), temporary wetland density (positive but weaker effect), precipitation, elevation, and snow water equivalent (Fig. 6). Ecoregion-level models revealed differences in key spatial predictors: irrigation and wetland density were important in the Great Basin, whereas climate-related variables were more influential in the Northern Great Plains (Fig. 8). VIMP score plots and PDP figures for all ecoregion-level spatial models are shown in Appendix B, Figures B.17-B.32.

**Figure 6.**
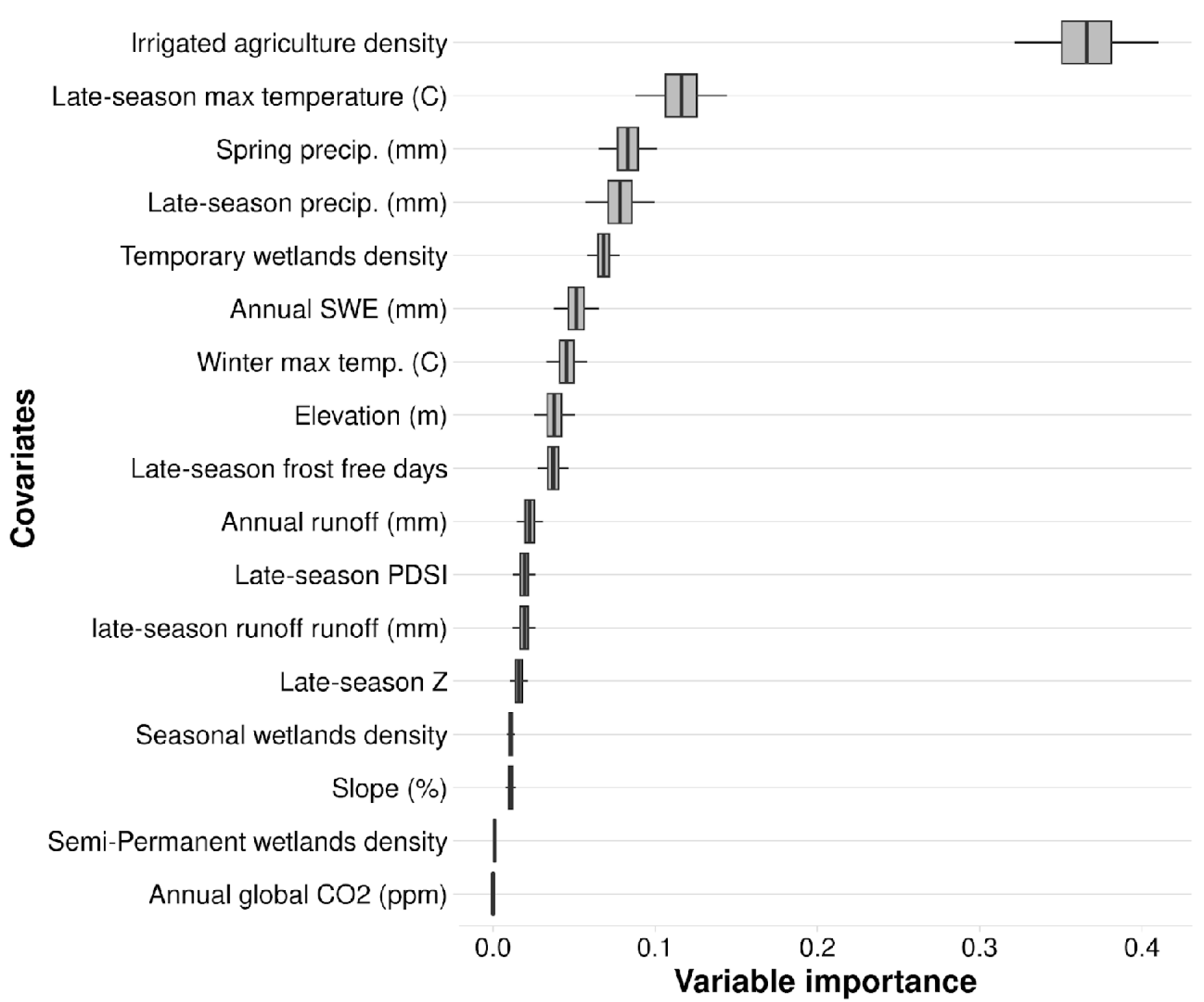
Variable importance (VIMP) of variables predicting mesic resource productivity across the sagebrush biome from 1986 to 2023 using the spatial random forest analysis. Variable importance was assessed using the Breiman-Cutler permutation method, with higher values indicating greater predictive power. Center black lines represent median importance values from 100 subsampled VIMP scores; boxes cover the 25th to 75th percentiles; whiskers extend to 95% confidence intervals. Variable importance is standardized by dividing by the variance of Y. Variables shown as densities are measured in hectares per 500-meter radius.

**Figure 7.**
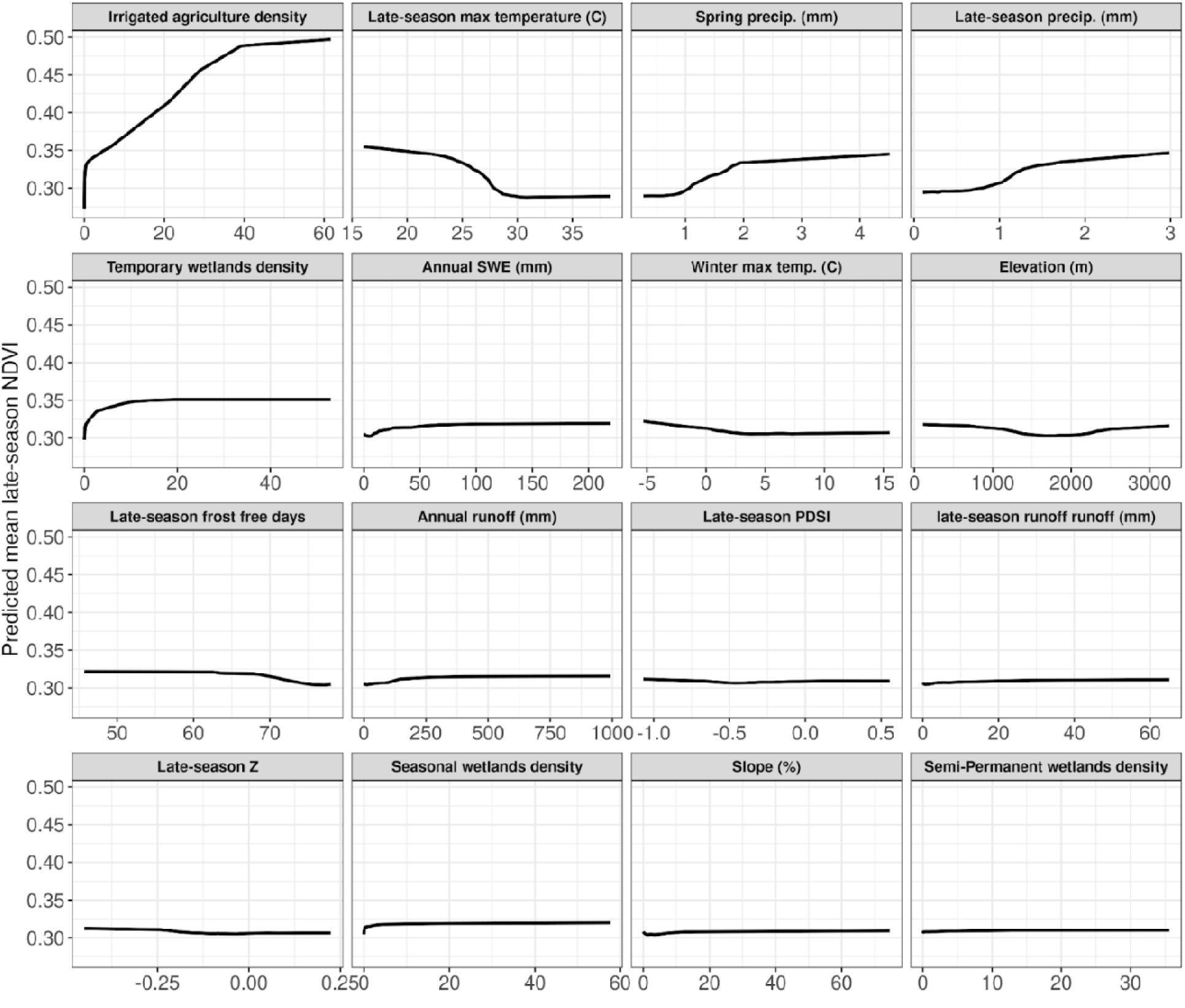
Partial dependency plots for each variable used in the spatial model on predicting mesic resource productivity in the sagebrush biome (1986-2023). Y-axis values are the predicted mean late-season NDVI. The X-axis shows untransformed variable values. Variables shown as densities are measured in hectares per 500-meter radius.

**Figure 8.**
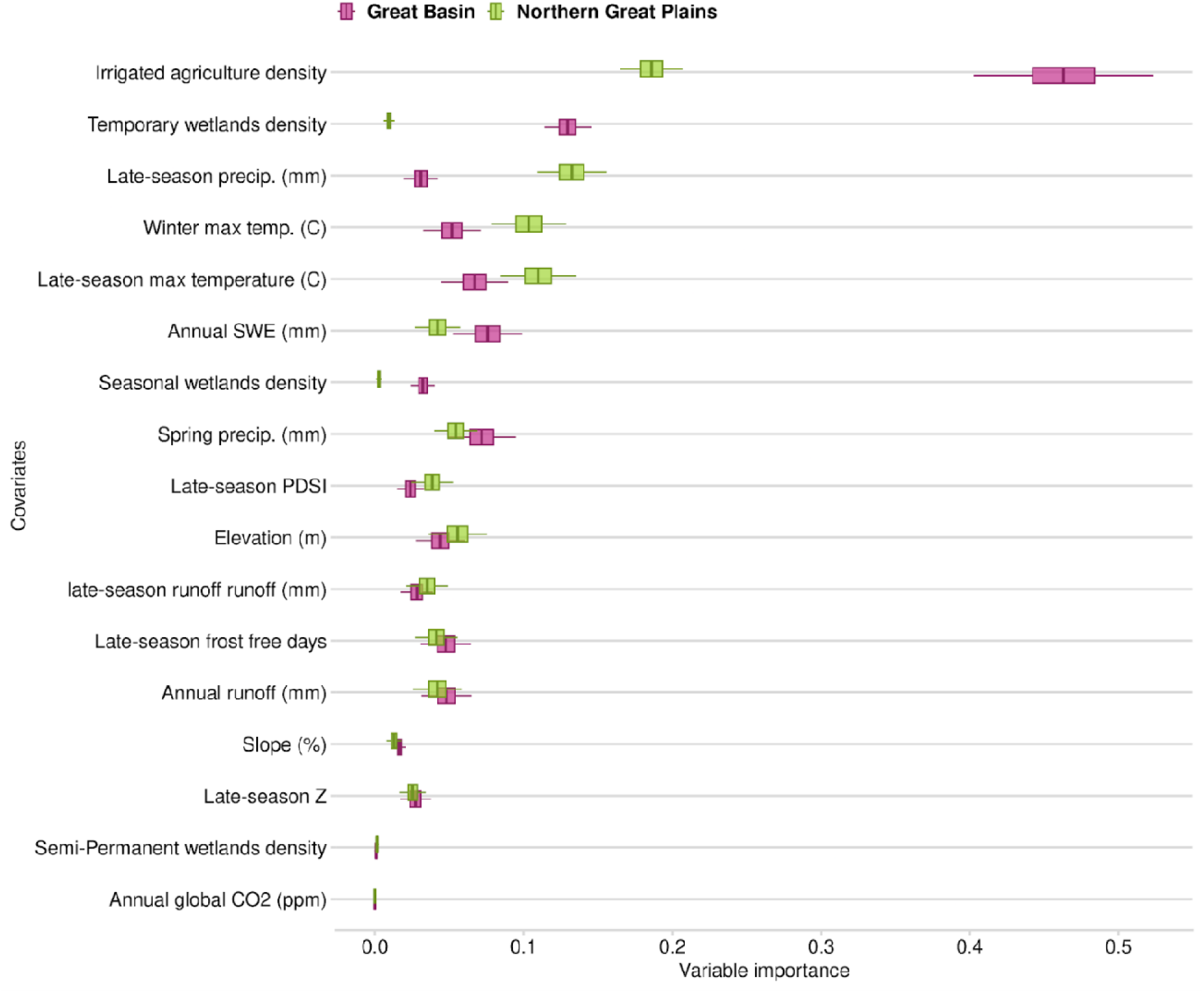
Comparison of variable importance (VIMP) scores of variables predicting mesic resource productivity for the Great Basin and Northern Great Plains from 1986 to 2023 using the spatial random forest analysis. Variables shown as densities are measured in hectares per 500-meter radius.

## 4 Discussion

Our analysis reveals a fundamental shift in how productivity in mesic resources across western U.S. rangelands responds to environmental limiting factors, such as drought. Although drought frequency remained relatively stable from 2005 to 2024, mesic productivity increased and became entirely decoupled from PDSI. This historical drought metric, which explained 92% of the productivity variation between 1984 and 2004, accounted for only 28% of the variation over the last two decades (Fig. 3). This divergence suggests that factors other than meteorological drought now dictate mesic productivity in these systems. By distinguishing between temporal and spatial predictors, our models identify the primary environmental and anthropogenic factors associated with this apparent resilience in mesic productivity. Because our biome-wide analysis captures large spatial extents, these dynamics predominantly reflect those of large floodplain valley systems, which are heavily mediated by flood irrigation (Donnelly, Jensco, et al., 2024); consequently, these large-scale trends may not fully capture the dynamics of smaller, unregulated headwater streams.

During the historical baseline of our study (P1:1986–2004), PDSI was the strongest predictor of mesic productivity (Fig. 4). This aligns closely with established ecohydrological principles, showing meteorological drought severely restricts plant productivity by depleting soil moisture and shallow groundwater (Getabalew & Alemneh, 2019; Selmants et al., 2023). The Palmer-Z index, a short-term drought metric highly sensitive to soil moisture (Karl, 1986), also strongly predicted productivity during this era. However, in the recent period (P2: 2005 - 2023), the predictive strength of PDSI decreased (Fig. 4). The fact that mesic systems maintained productivity during the megadroughts of the 21st century suggests that maintaining floodplain connectivity with groundwater can buffer these habitats against acute climate stress (Stromberg et al., 2013; A. P. Williams et al., 2020).

As the predictive strength of meteorological drought waned in P2, global atmospheric CO_2_ concentrations emerged as the dominant temporal predictor of mesic productivity (Fig. 4). This shift aligns with growing global evidence of a CO_2_ fertilization effect, which has increasingly driven the ‘greening but drying’ phenomenon observed in drylands since the early 2000s (Gonsamo et al., 2021; Mankin et al., 2017; Wang et al., 2022; Zhang et al., 2022). Elevated CO_2_ likely enhances plant water use efficiency (WUE), enabling riparian herbaceous vegetation to sustain photosynthesis and growth even when meteorological drought persists (Ukkola et al., 2015). However, interpreting this ‘greening’ as purely beneficial requires ecological caution. Our NDVI-based analysis measures total vegetation vigor and cannot perfectly isolate herbaceous from woody productivity. Recent experimental evidence indicates that woody plants respond to elevated CO_2_ with nearly double the productivity increase (24%) of grasses (13%) under similar conditions (Pan et al., 2022). Therefore, while rising CO_2_ may currently boost overall mesic greenness when water is available in floodplains, it could inadvertently accelerate woody encroachment into riparian zones, fundamentally altering the competitive dynamics and functional composition of these vital herbaceous habitats (Keen et al., 2023; Körner, 2006; Saintilan & Rogers, 2015).

While CO_2_ largely explains the temporal shift in productivity, irrigation density emerged as the overwhelming spatial predictor across the biome (Fig. 6). Consistent with recent findings, large agricultural valley floodplains act as anthropogenic subsidies, sustaining productive mesic resources through late summer via flood irrigation and subsequent groundwater recharge (Donnelly, Jensco, et al., 2024; Essaid & Caldwell, 2017; Gordon et al., 2020). Donnelly et al. (2024) found that 93% of flood irrigation for grass-hay production occurred within historical wetland systems, which currently supports 58% of remaining palustrine wetlands in the western U.S. Unsurprisingly, the densities of both irrigation and temporary wetlands were among the top spatial predictors in our models. Partial dependence plots illustrated a strong positive threshold effect: mesic productivity increased sharply with the presence of irrigation before plateauing at higher densities (Fig. 7). While flood irrigation may help sustain floodplain connectivity and supports late-season mesic resources, excessive diversions can deplete instream flows below thresholds needed to sustain aquatic communities (Donnelly, Jensco, et al., 2024; Gordon et al., 2020), highlighting the need for management approaches that balance agricultural and ecological water demands.

Other spatial predictors, including snow water equivalent, temperature, and precipitation, highlighted regional vulnerabilities (Fig. 6). The Northern Rockies and Wyoming Basin exhibited the largest productivity increases during P2 (Appendix A, Fig. A.3), likely reflecting increased precipitation that extends the growing season (Polley et al., 2013). In the Great Basin, snow water equivalent was highly predictive, as winter snowpack drives both groundwater recharge and summer streamflow (Lauenroth et al., 2014). However, warming winters threaten this dynamic by shifting precipitation from snow to rain, potentially reducing snow-derived runoff in the West by one-third by century’s end (Dettinger et al., 2015; Li et al., 2017; Lundblad et al., 2022). Conversely, in the Northern Great Plains, spatial models emphasized summer precipitation and temperature (Fig. 8), reflecting a landscape driven by intense, episodic summer storms rather than snowmelt (Lauenroth et al., 2014; Morgan et al., 2011). Because groundwater recharge, streamflow timing, and growing season length are driven by fundamentally different processes in these regions, effective conservation strategies must be customized to their specific regional hydroclimates.

Several limitations warrant consideration. First, we focused on late-season (July 15–September 30) productivity, the most critical period for wildlife forage, meaning these findings do not capture early-season dynamics, though late-season trends largely mirrored annual patterns (Appendix A, Fig. A.4). Second, our reliance on 30-m resolution imagery captures large floodplain dynamics well but may obscure fine-scale changes in narrow headwater riparian corridors. Finally, the decoupling between drought and productivity since 2005 should not be interpreted as universal drought immunity. Rather, it reflects physiological adaptations and the massive redistribution of human water in select landscapes. In fact, approximately 7% of mesic pixels exhibited declining productivity during P2, largely concentrated in areas experiencing acute groundwater depletion or localized land-use conversion, as is the case in the Harney Basin of Eastern Oregon (C. Albano et al., 2020) (Appendix A, Fig. A.5), underscoring how local stressors can override broad biome-wide patterns.

## 5 Conclusions

Our findings reveal a fundamental shift in how western U.S. mesic resources respond to drought. In the last two decades, late-season productivity has become decoupled from meteorological drought indices, with rising atmospheric CO_2_ and human water management emerging as the strongest predictors of sustained productivity. Yet, this apparent resilience is spatially variable and should not be mistaken for stability. Key spatial indicators of productivity differ across regions, from snowmelt-dependent basins to pulse-precipitation-driven plains. Critically, the resilience these trends reflect is not intrinsic to the ecosystems themselves — it is contingent on continued flood irrigation and stable snowpack, both of which face increasing pressure from climate change and water policy. These results argue for a reorientation of conservation strategy. Passive climate monitoring alone is insufficient; maintaining mesic resources will require active management of water availability at the landscape scale. Where flood irrigation sustains late-season groundwater recharge, strategic partnerships with private landowners are critical to preserving these hydrological benefits as water policy tightens. Where agricultural water use is limited, process-based restoration is needed to retain water on the landscape. As water scarcity grows, resilience will depend on adaptive, localized approaches that integrate beneficial agricultural water use and restoration to keep mesic resources available for productive forage.

## Data Availability

Source code from Google Earth Engine and R scripts required for data reproducibility, generation, tables, and figures are accessible via a GitHub repository (https://github.com/krisMueller/shifting-resilience). Data extracted from Google Earth Engine and used in R script analyses have been permanently archived in a Zenodo repository (https://doi.org/10.5281/zenodo.18672909).

## Acknowledgments

This research used compute and storage resources provided by Google Earth Engine. The US Bureau of Land Management of Montana/Dakotas provided funding for this research (grant code: F21AC00546) through the Intermountain West Joint Venture (IWJV) with help from the USDA-NRCS Working Lands for Wildlife.

## Author CRediT Statement

**K. R. M:** Conceptualization, Methodology, Software, Formal analysis, Writing - original draft preparation. **S. L. M**: Conceptualization, Methodology, Supervision, Writing - reviewing and editing, **J. S. K:** Writing - reviewing and editing, **J. T. S:** Methodology, Writing - reviewing and editing, **J. P. D:** Writing - reviewing and editing, **D. E. N:** Conceptualization, Supervision, Writing - reviewing and editing.

## Declaration of Generative AI-assisted Technologies

While preparing this manuscript, the authors used Grammarly to correct spelling and grammar errors. AI technologies were not used in any part of the methodology or data generation process. The authors reviewed and edited the manuscript and take full responsibility for the content of the published article.

