## Appendix A & B for "Shifting Resilience: Trends and Predictors of Mesic Resource Productivity in Western U.S. Rangelands"

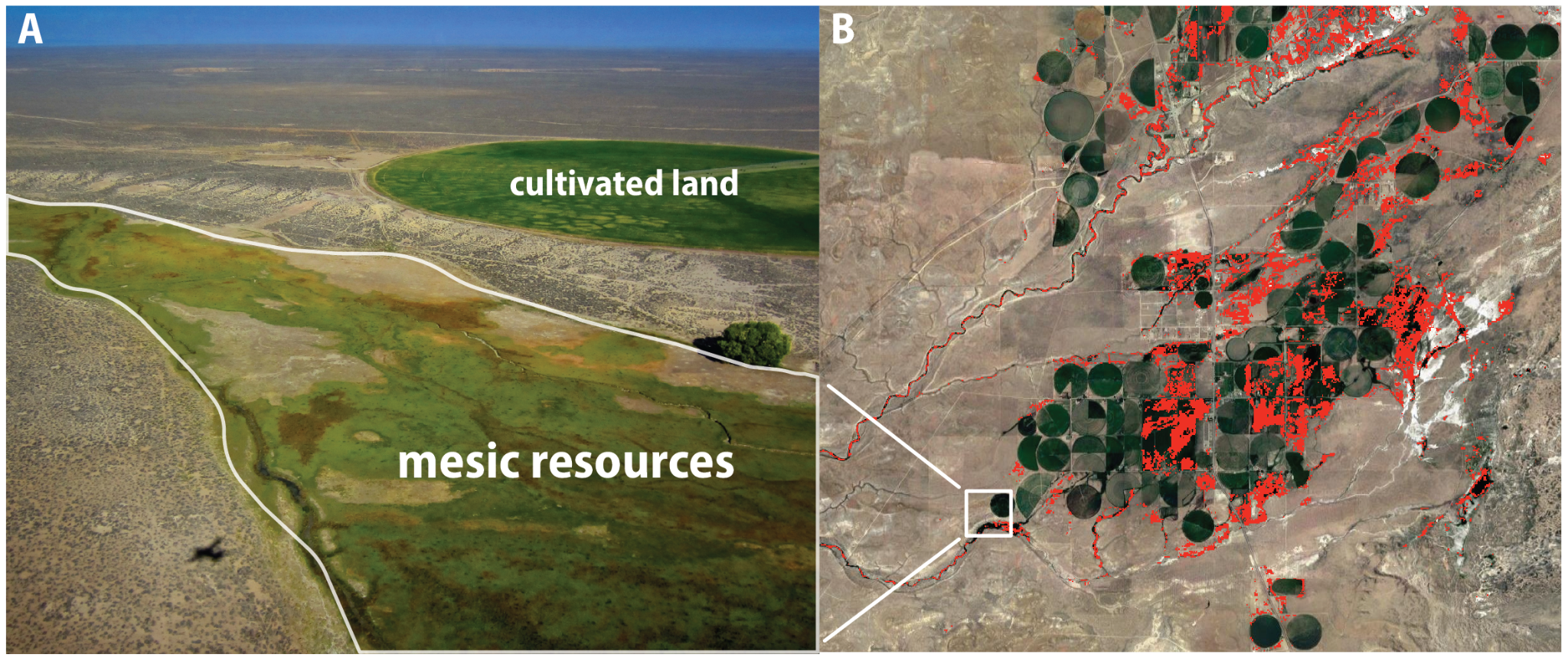

Figure A.1. An example of late-season mesic resources and the final mesic resource mask used for analysis near Farson, Wyoming. Panel A displays an aerial view of mesic resources near cultivated lands; photo credit: Patrick Donnelly (2018). Panel B shows the final classification with mesic resource pixels (red) after filtering out cultivated lands and other non-mesic land cover types.

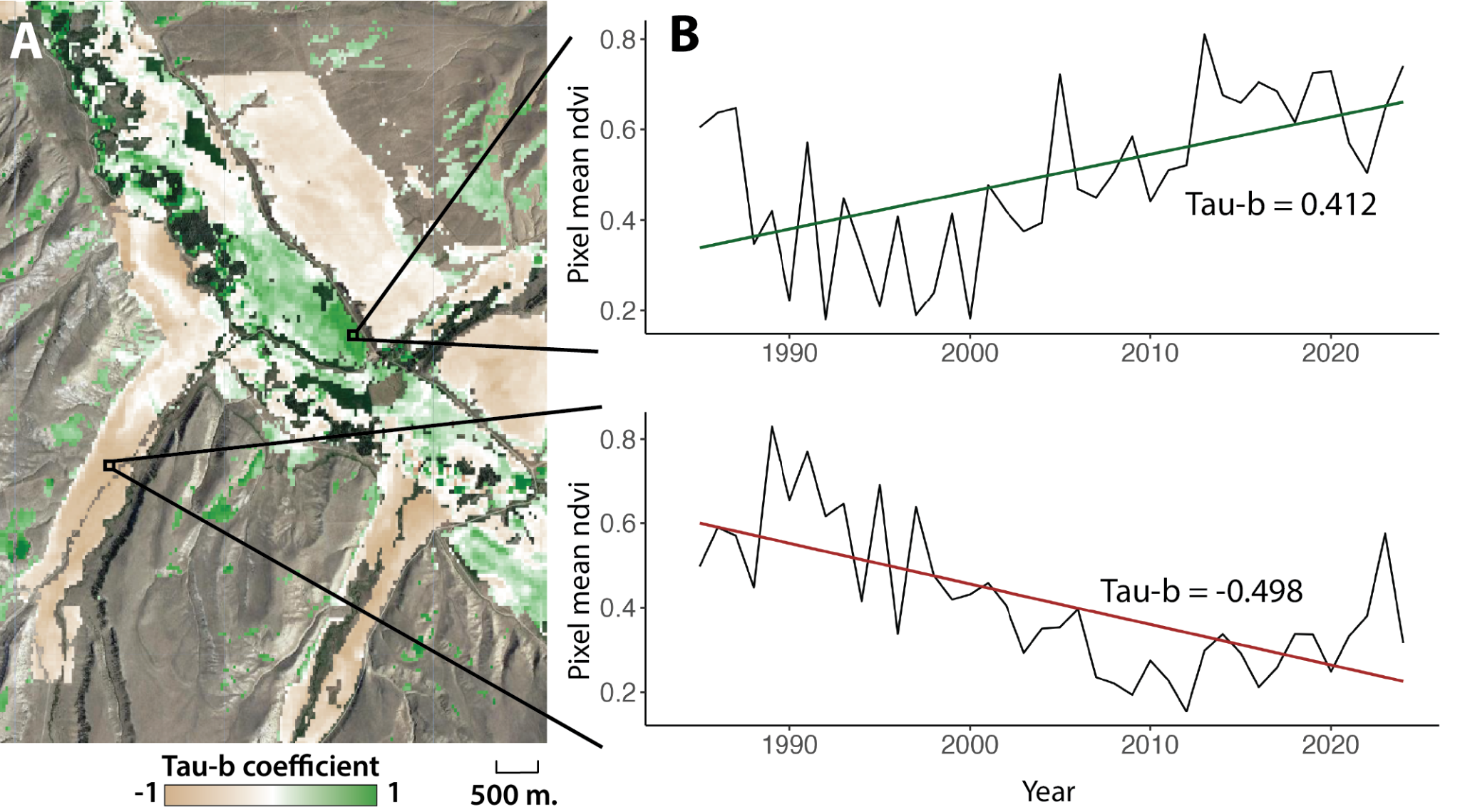

Figure A.2. An example of Mann-Kendall (MK) tau-b trend analyses in a riparian floodplain. A: Spatial distribution of tau-b values, with brown indicating decreasing productivity (tau-b near -1), green indicating increasing productivity (tau-b near 1), and white indicating no trend (tau-b close to 0). B: time series of NDVI values for representative pixels, showing examples of positive and negative trends with their corresponding tau-b values. The green and red lines in panel B are linear trend lines and do not directly represent the MK tau-b trend.

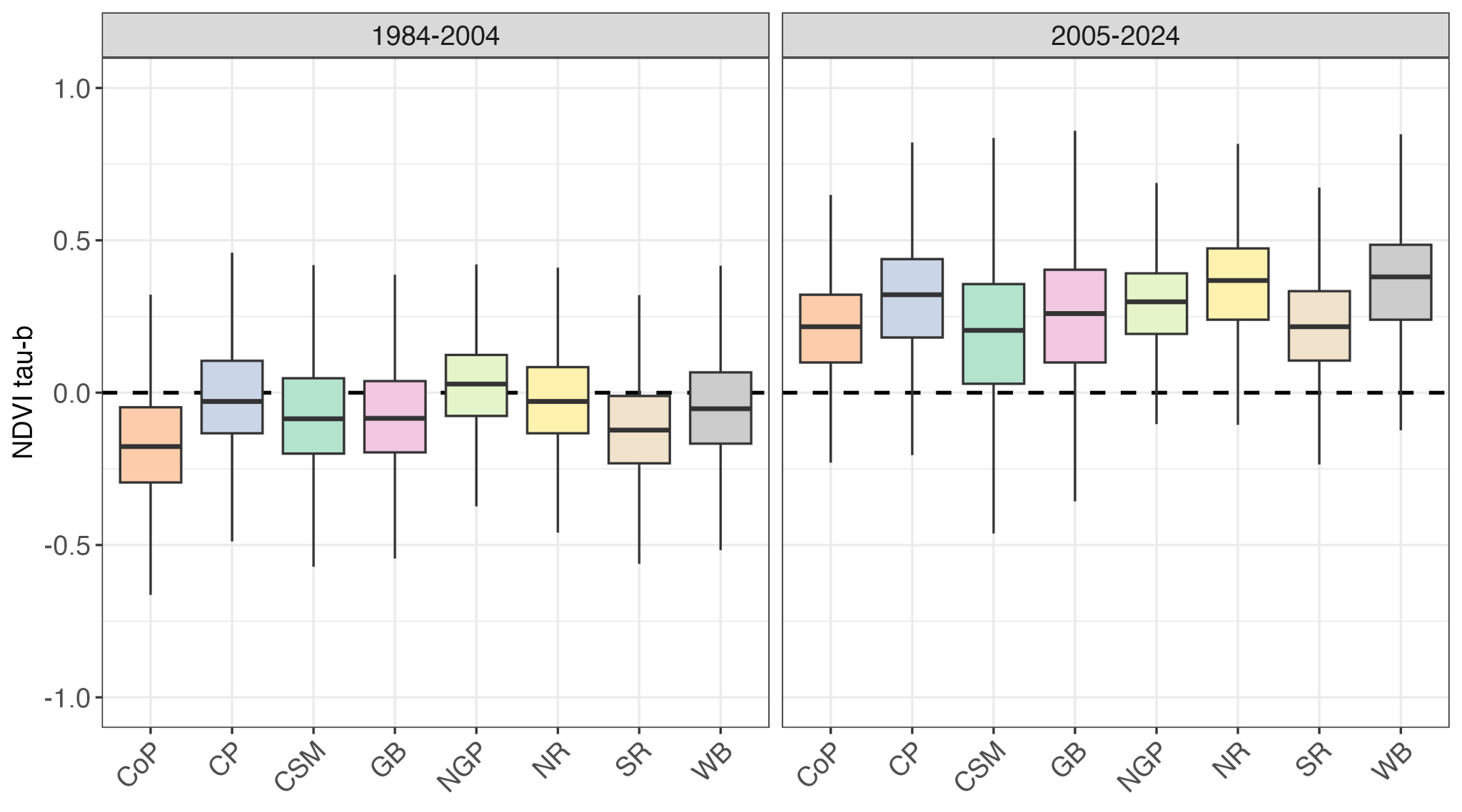

Figure A.3. Regional patterns in mesic resource productivity trends across ecoregions. Boxplots illustrate the distribution of Mann-Kendall Tau-b values for two time periods: 1984-2004 (P1) and 2005-2024 (P2). The boxes show the 25th (lower), 50th (median), and 75th (upper) percentiles of sampled data. Positive Tau-b values (approaching 1) indicate increasing productivity trends, negative values (approaching -1) indicate decreasing trends, and values near 0 suggest no trend. Each boxplot displays 200,000 randomly sampled points from mesic areas within each ecoregion: Colorado Plateau (CoP), Columbia Plateau (CP), Cascade/Sierra Mountains (CSM), Great Basin (GB), Northern Great Plains (NGP), Northern Rockies (NR), Southern Rockies (SR), and Wyoming Basin (WB). Notice the shift toward positive trends in P2, especially in the Northern Rockies and Wyoming Basin.

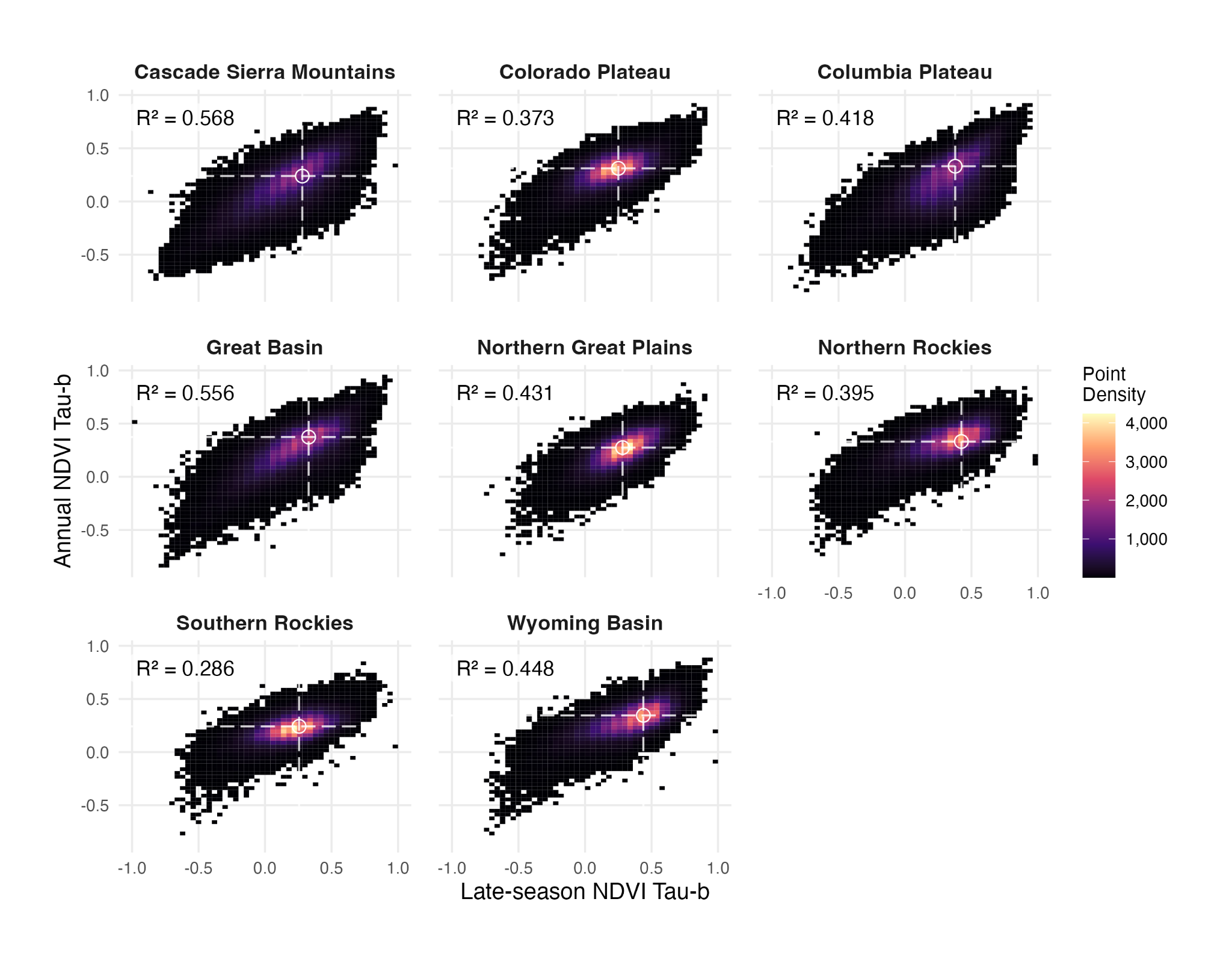

Figure A.4. The relationship between late-season (July 15 - Sep. 30) mean NDVI trends (Mann-Kendall Tau-B) and annual NDVI trends (2005-2024) using 200,000 sampled points per ecoregion.

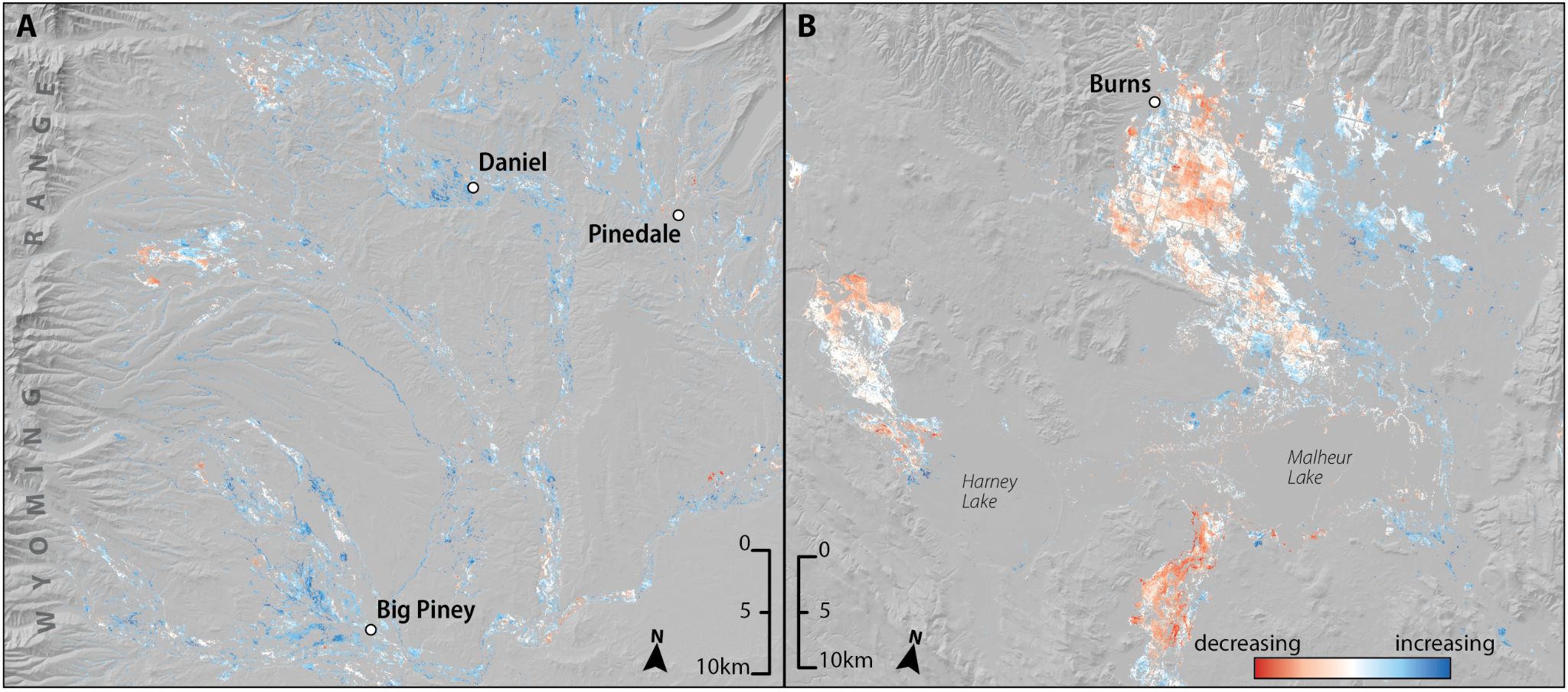

Figure A.5. Visualization of mesic resource trends during P2 (2005 - 2024) of the study period in A) the Upper Green River Basin of western Wyoming, and B) Harney Basin in Eastern Oregon. Mesic pixels shown in red had increasing productivity values during P2, while blue pixels showed increasing productivity values.

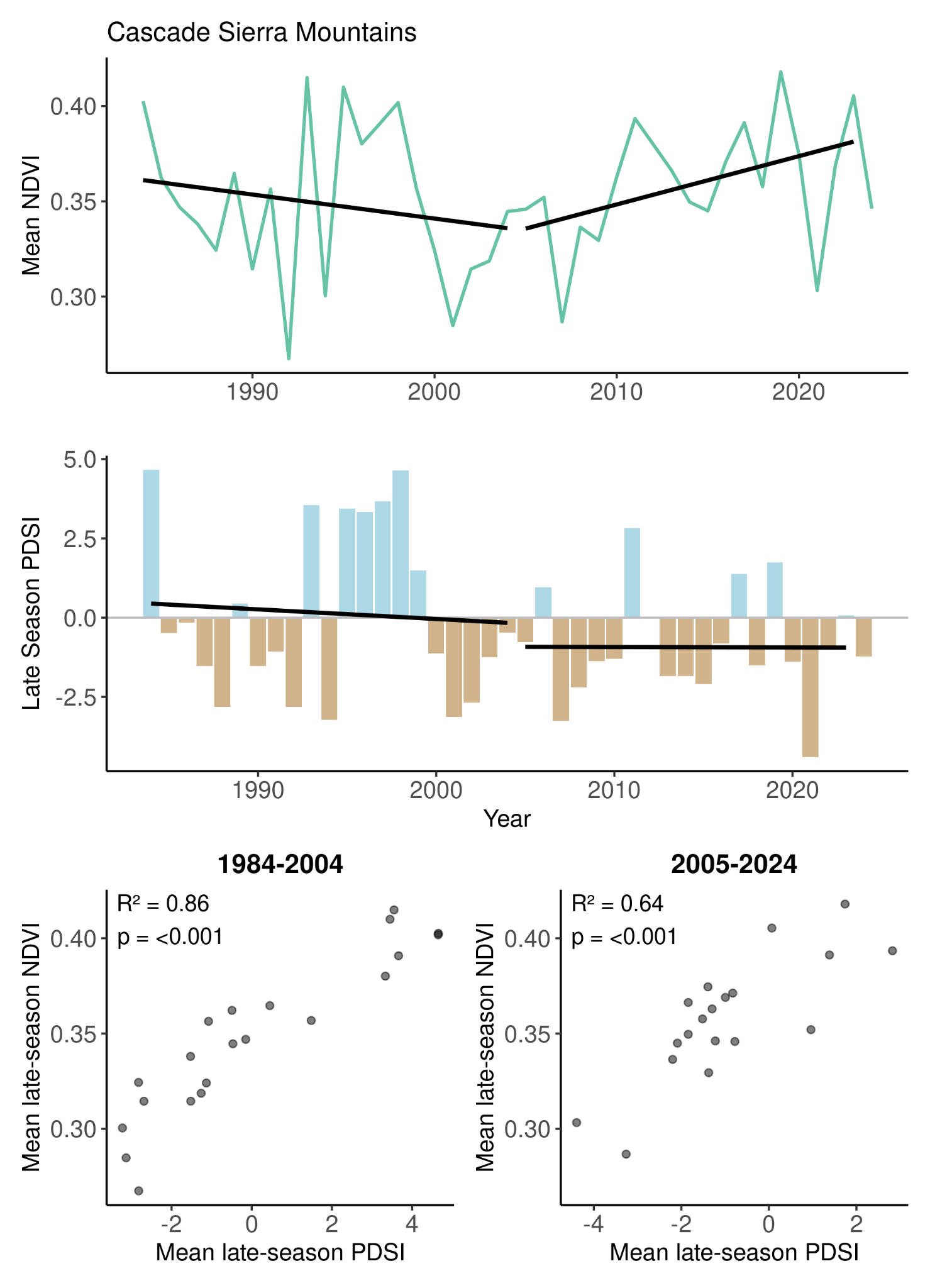

Figure A.7. The relationship between mean late-season (July 15 - September 30) mesic resource productivity and drought conditions across the Cascade Sierra Mountains region (1984-2024). Time series plots show the trends in the Normalized Difference Vegetation Index (NDVI) and the Palmer Drought Severity Index (PDSI) for periods 1 (1984-2004) and 2 (2005-2024), with linear regression statistics for both periods. Black lines indicate linear regression trends for each period.

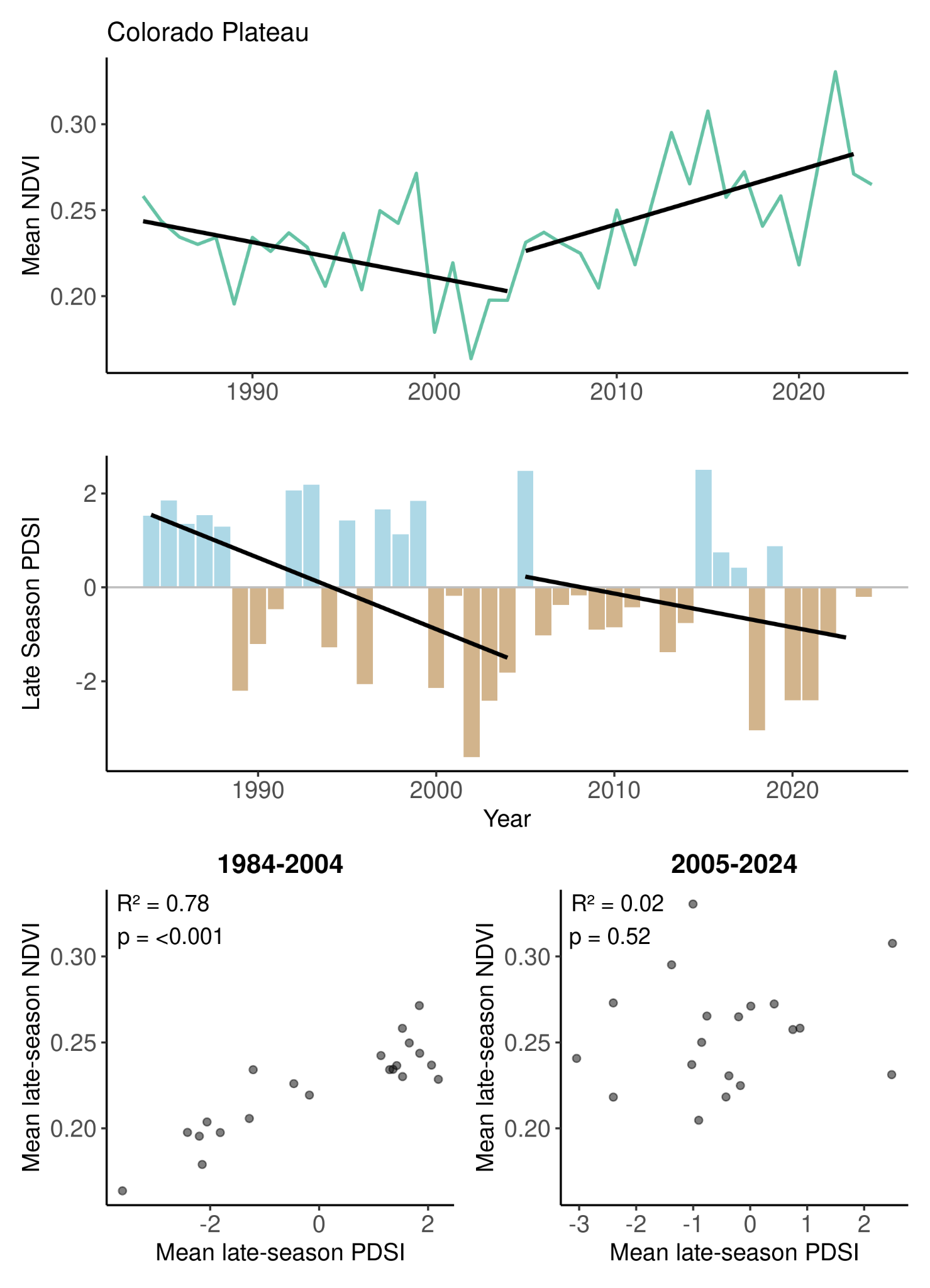

Figure A.8. The relationship between mean late-season (July 15 - September 30) mesic resource productivity and drought conditions across the Colorado Plateau region (1984-2024). Time series plots show the trends in the Normalized Difference Vegetation Index (NDVI) and the Palmer Drought Severity Index (PDSI) for periods 1 (1984-2004) and 2 (2005-2024), with linear regression statistics for both periods. Black lines indicate linear regression trends for each period.

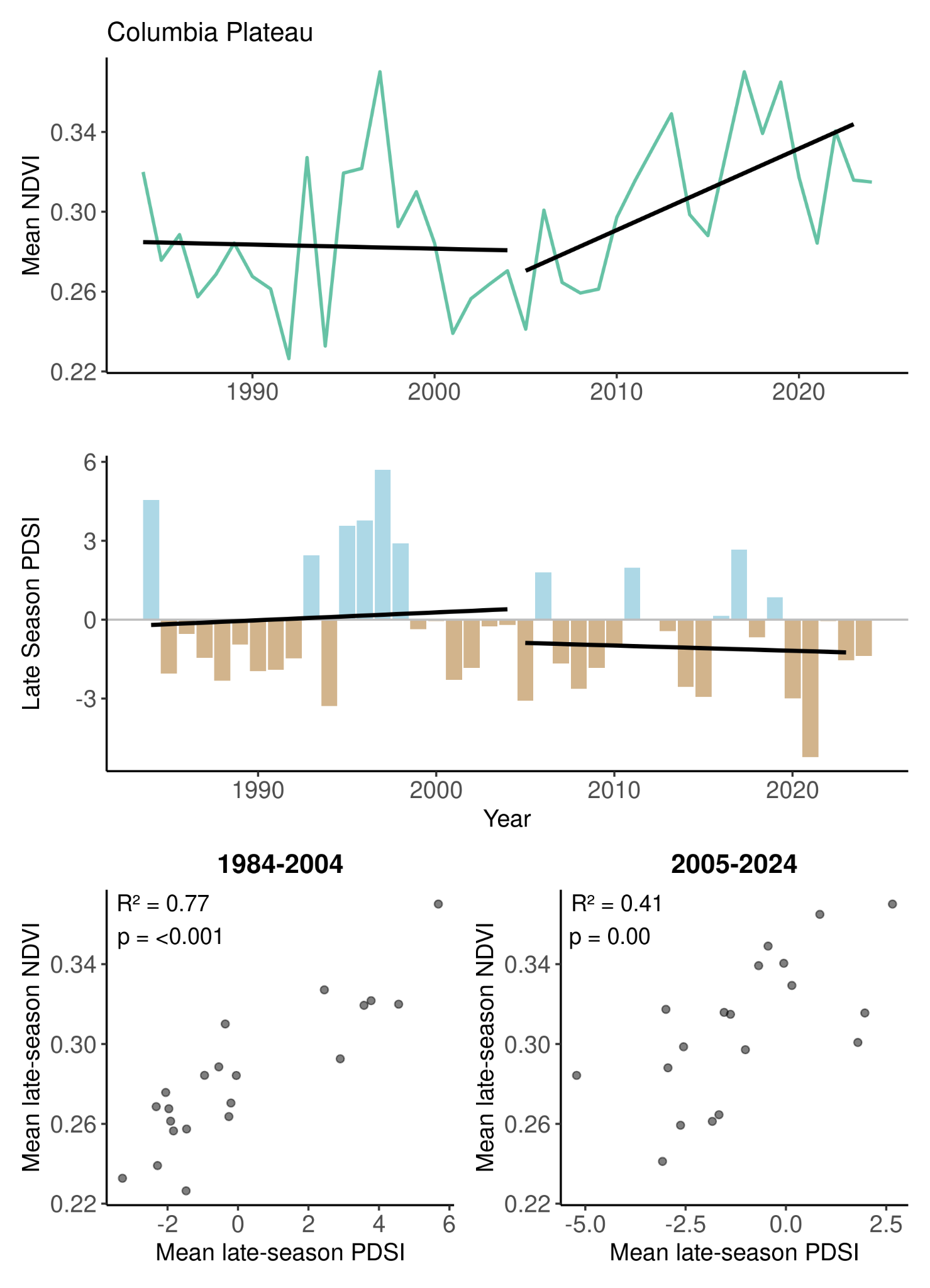

Figure A.9. The relationship between mean late-season (July 15 - September 30) mesic resource productivity and drought conditions across the Columbia Pleteau region (1984-2024). Time series plots show the trends in the Normalized Difference Vegetation Index (NDVI) and the Palmer Drought Severity Index (PDSI) for periods 1 (1984-2004) and 2 (2005-2024), with linear regression statistics for both periods. Black lines indicate linear regression trends for each period.

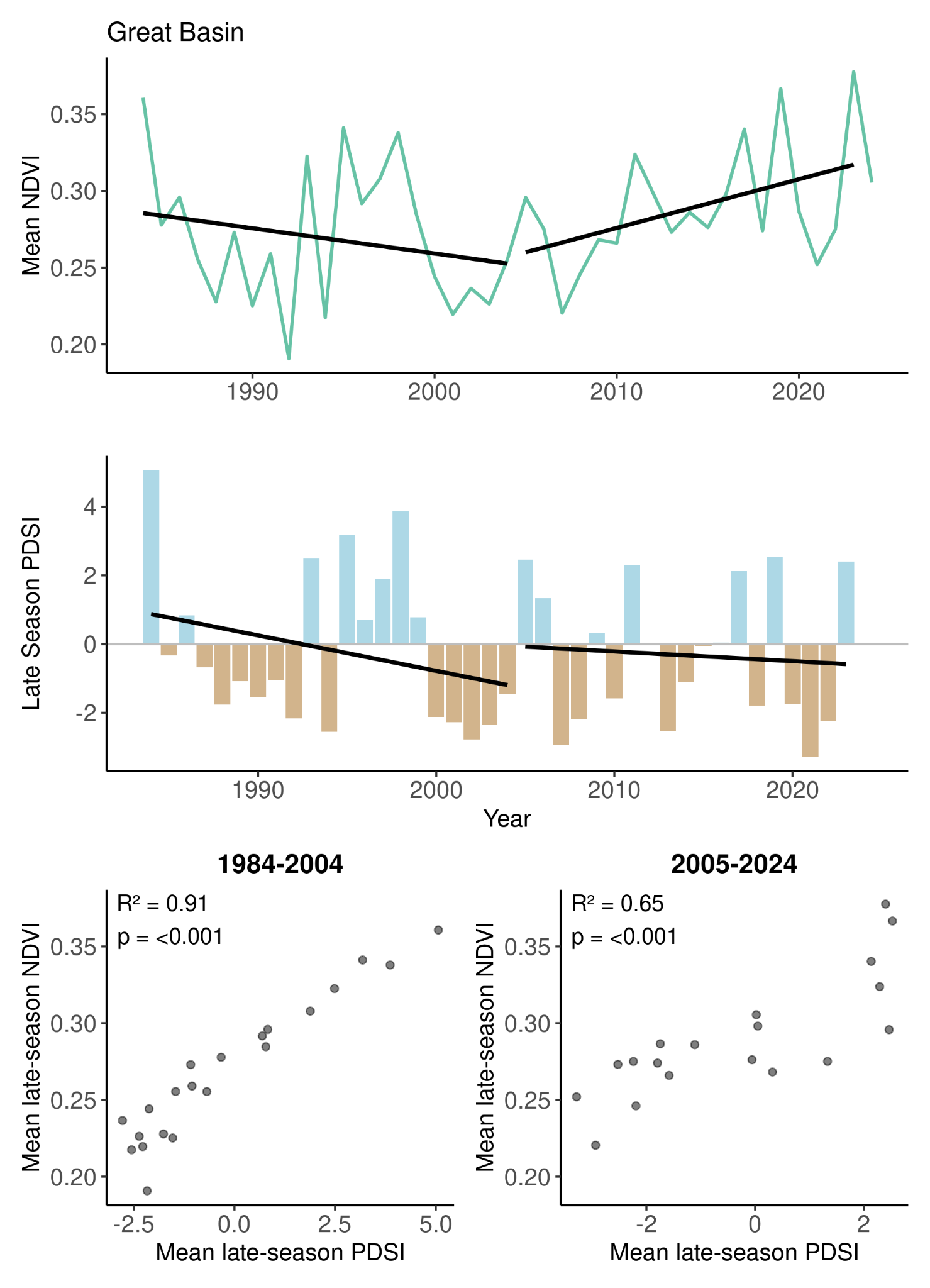

Figure A.10. The relationship between mean late-season (July 15 - September 30) mesic resource productivity and drought conditions across the Great Basin region (1984-2024). Time series plots show the trends in the Normalized Difference Vegetation Index (NDVI) and the Palmer Drought Severity Index (PDSI) for periods 1 (1984-2004) and 2 (2005-2024), with linear regression statistics for both periods. Black lines indicate linear regression trends for each period.

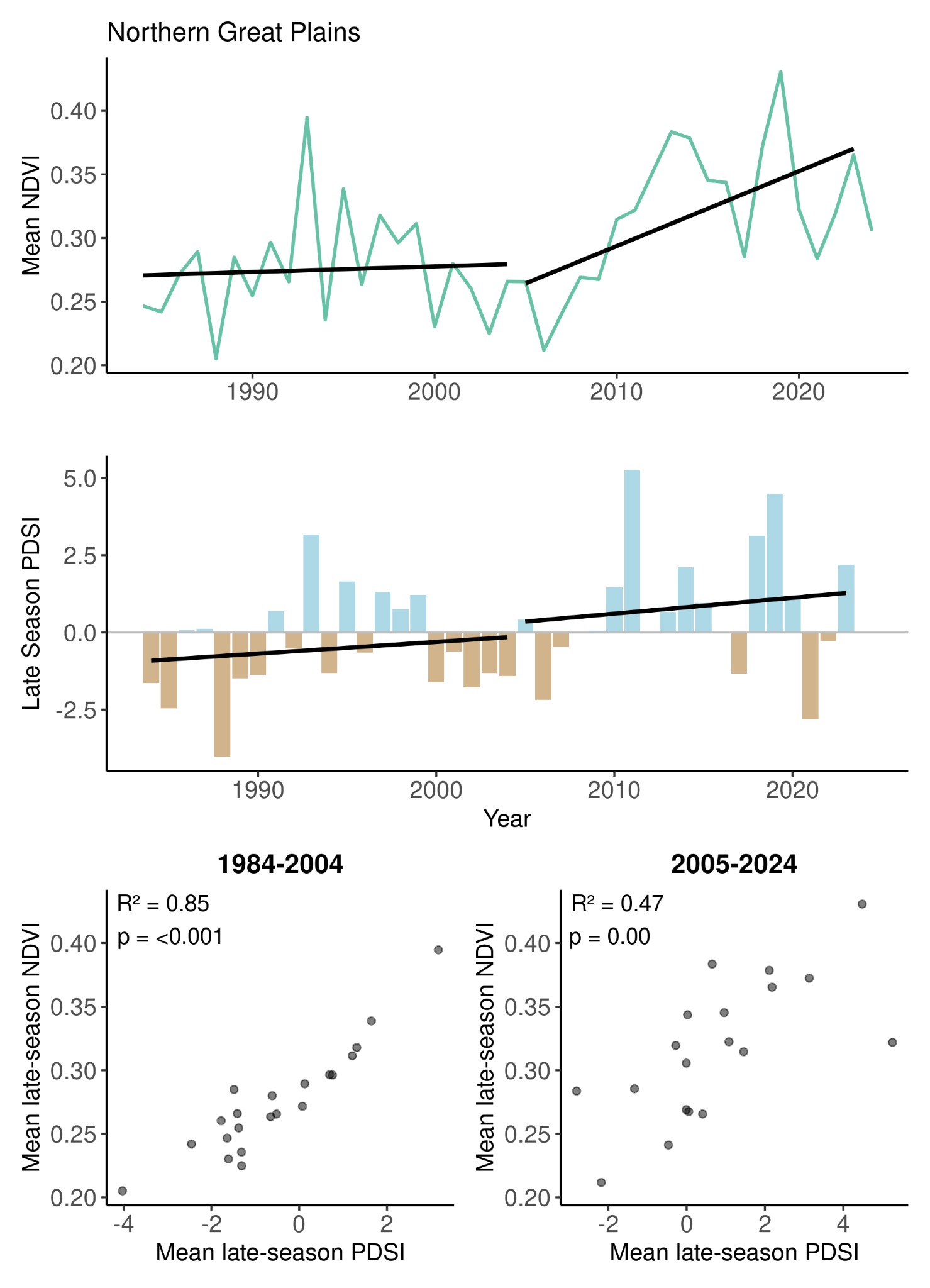

Figure A.11. The relationship between mean late-season (July 15 - September 30) mesic resource productivity and drought conditions across the Northern Great Plains region (1984-2024). Time series plots show the trends in the Normalized Difference Vegetation Index (NDVI) and the Palmer Drought Severity Index (PDSI) for periods 1 (1984-2004) and 2 (2005-2024), with linear regression statistics for both periods. Black lines indicate linear regression trends for each period.

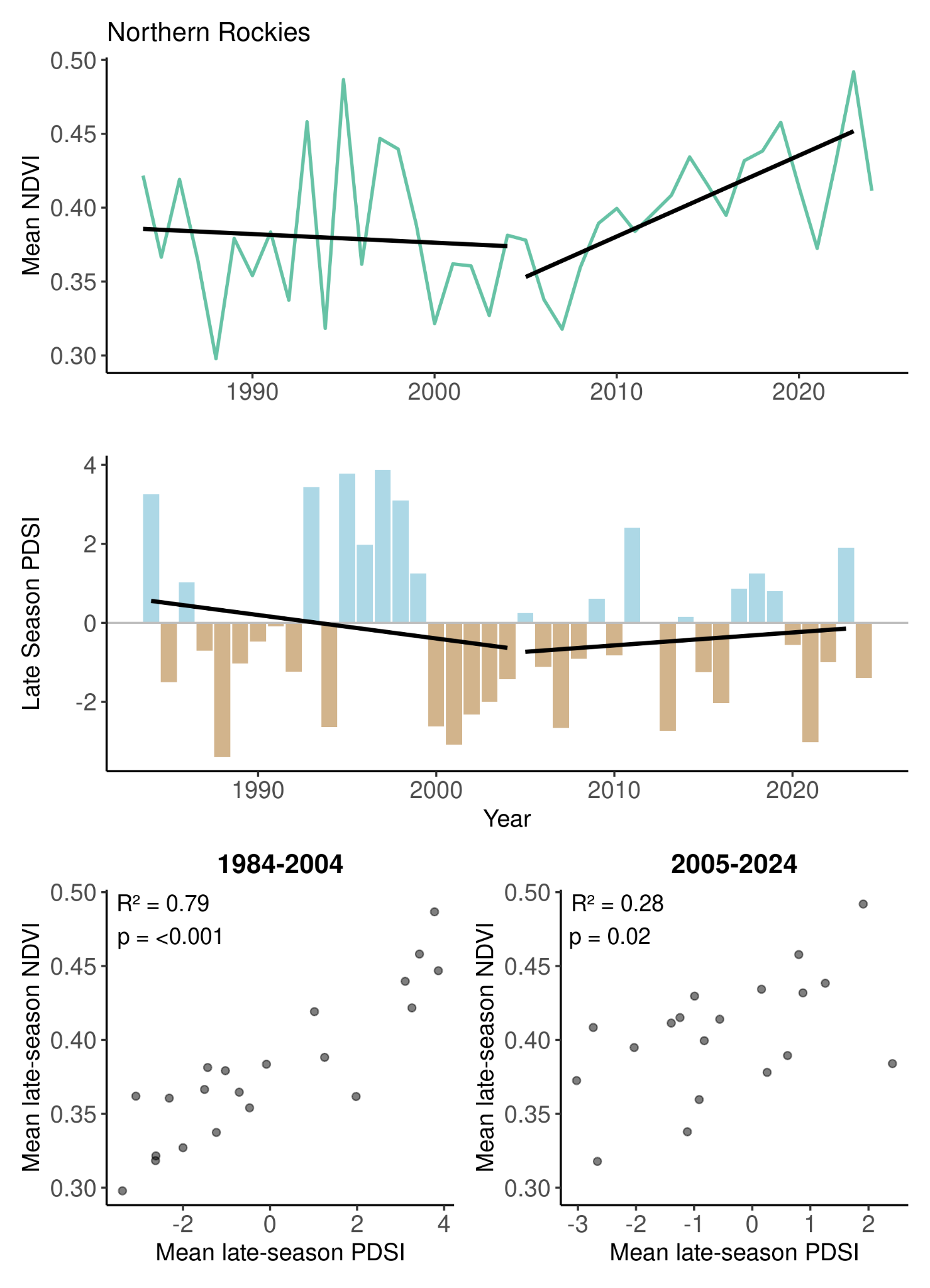

Figure A.12. The relationship between mean late-season (July 15 - September 30) mesic resource productivity and drought conditions across the Northern Rockies region (1984-2024). Time series plots show the trends in the Normalized Difference Vegetation Index (NDVI) and the Palmer Drought Severity Index (PDSI) for periods 1 (1984-2004) and 2 (2005-2024), with linear regression statistics for both periods. Black lines indicate linear regression trends for each period.

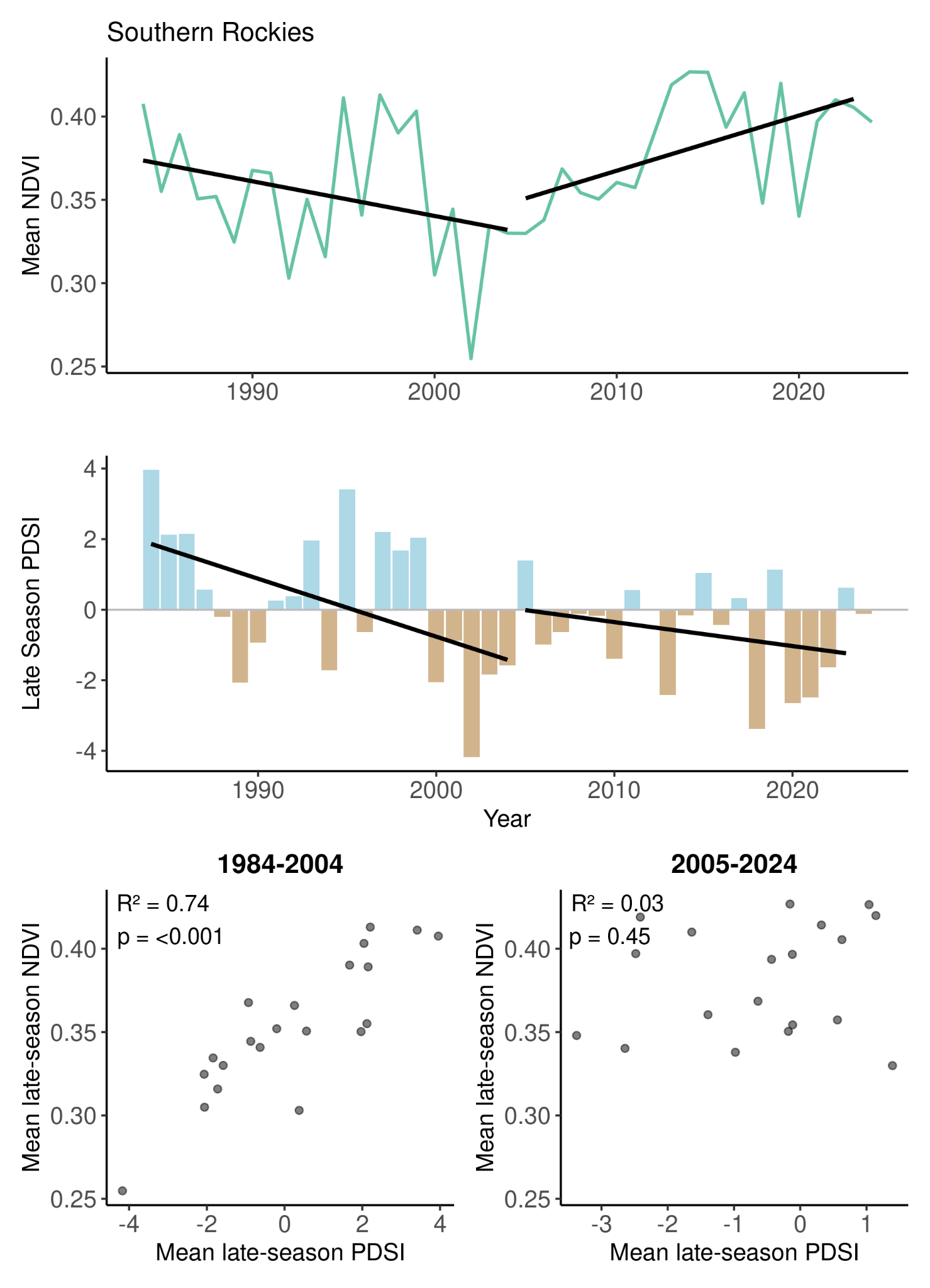

Figure A.13. The relationship between mean late-season (July 15 - September 30) mesic resource productivity and drought conditions across the Southern Rockies region (1984-2024). Time series plots show the trends in the Normalized Difference Vegetation Index (NDVI) and the Palmer Drought Severity Index (PDSI) for periods 1 (1984-2004) and 2 (2005-2024), with linear regression statistics for both periods. Black lines indicate linear regression trends for each period.

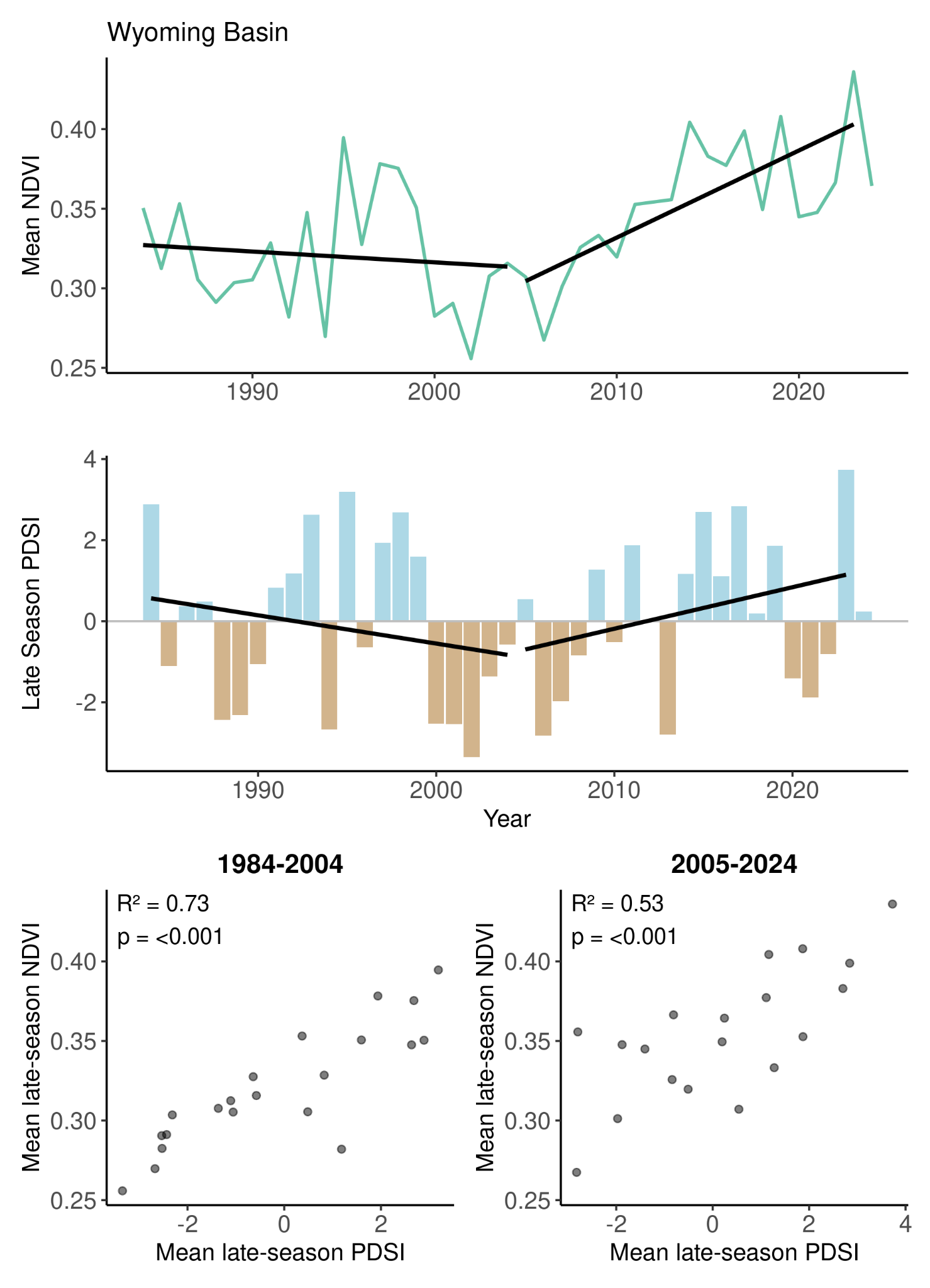

Figure A.14. The relationship between mean late-season (July 15 - September 30) mesic resource productivity and drought conditions across the Wyoming Basin region (1984-2024). Time series plots show the trends in the Normalized Difference Vegetation Index (NDVI) and the Palmer Drought Severity Index (PDSI) for periods 1 (1984-2004) and 2 (2005-2024), with linear regression statistics for both periods. Black lines indicate linear regression trends for each period.

Table A.1. Datasets used for mesic mask creation within Google Earth Engine.

| **Image mask** | **Dataset** |
| --- | --- |
| Mesic identification | USGS Landsat Collection ^1^ |
| Urban development | USDA NASS CDL ^2^ |
| Cultivated agriculture | USDA NASS CDL ^2^ |
| Water (supplemental) | USDA NASS CDL ^2^ |
| Forests/woodlands and trends | Rangeland Analysis Platform ^3^ |
| High elevation | USGS 3DEP 1m ^4^ |
| Riparian and wetlands | USDA NASS CDL ^2^, LANDFIRE BPS ^5^, VBET^6^,  Riverscapes Data Exchange^7^ |

^1^ Landsat (5, 7, 8, 9) Level 2, Collection 2, Tier 2 Surface Reflectance. Courtesy of the U.S. Geological Survey.

^2^ USDA National Agricultural Statistics Service Cropland Data Layer. {1997-2023}. Published crop-specific data layer [Online – Google Earth Engine]. (accessed 2024-11-22; verified 2024-11-22). USDA-NASS, Washington, DC.

^3^ Rangeland Analysis Platform Cover (Allred et al., 2021).

^4^ U.S. Geological Survey, 3D Elevation Program 1-Meter Resolution Digital Elevation Model.

^5^ LANDFIRE, 2016, Existing Vegetation Type Layer, LANDFIRE 2.0.0, U.S. Department of the Interior, Geological Survey, and U.S. Department of Agriculture.

^6^ (Gilbert et al., 2016)

^7^ *Riverscapes Data Exchange*. (2026). Riverscapes.net. https://data.riverscapes.net/

Table A.2. A detailed list of datasets used to identify herbaceous mesic resources across the sagebrush biome.

| **Masking Dataset** | **Details** |
| --- | --- |
| Mesic identification | Pixels with a mean late-season NDVI value ≥ 0.3 across at least one year in the time series were considered possible mesic resources and included in our mesic mask. |
| Urban development  Cultivated agriculture  Water (supplemental) | The Cropland Development layer has annual 30-meter pixel values for urban development and water from 1997 to 2022 within Google Earth Engine. Cultivated pixels began with the dataset in 2011.  Any pixels classified as these data were removed from our mesic mask. |
| Forest/Woodlands  Shrub/Tree cover trends | Pixels that were ever classified as ≥ 10% tree cover from RAP (1986 - 2023) were excluded from the mesic mask.  Any pixel that had a significant increase or decrease in tree or shrub cover (using RAP) from 1984 to 2024 using *ee.Reducer.kendallsCorrelation* |
| High elevation | Pixels were retained in the mesic mask if they were below an elevation percentile within each ecoregion:  Rocky Mountains = 90th percentile  Great Basin or Cascade/Sierra Mountains = 97th percentile  No filter was applied to the remaining ecoregions |
| Riparian areas | Pixels that were ever classified as riparian in the NASS CDL dataset (1997-2022) were retained for the mask.  LANDFIRE BPS data were used to retain pixels classified as ‘riparian’ from the 2022 dataset.  Valley bottoms were extracted from the Valley Bottom Extraction Tool (VBET) with a probability of being a valley bottom threshold of ≥ 60% (0.6) |

Table A.3. Random forest model output accuracies (OOB) and errors for each model run in each of the temporal and spatial analysis models.

| **Ecoregion** | **P1**  **Temporal Samples** | **P1 Temporal R^2^ (OOB)** | **P1**  **Temporal**  **Error** | **P2 Temporal**  **Samples** | **P2 Temporal**  **R^2^ (OOB)** | **P2 Temporal**  **Error** | **Spatial**  **Samples** | **Spatial**  **R^2^ (OOB)** | **Spatial Error** |
| --- | --- | --- | --- | --- | --- | --- | --- | --- | --- |
| Sagebrush Biome | 226,726 | 0.54 | 0.002 | 215,624 | 0.57 | 0.002 | 12,000 | 0.65 | 0.005 |
| Cascade Sierra Mountains | 189,563 | 0.56 | 0.002 | 179,837 | 0.54 | 0.002 | 10,000 | 0.66 | 0.005 |
| Colorado Plateau | 189,375 | 0.56 | 0.001 | 179,247 | 0.62 | 0.002 | 10,000 | 0.67 | 0.002 |
| Columbia Plateau | 189,558 | 0.52 | 0.002 | 179,964 | 0.57 | 0.002 | 10,000 | 0.48 | 0.005 |
| Great Basin | 189,238 | 0.59 | 0.002 | 179,778 | 0.60 | 0.002 | 10,000 | 0.67 | 0.003 |
| Northern Great Plains | 189,565 | 0.69 | 0.001 | 179,894 | 0.78 | 0.001 | 10,000 | 0.52 | 0.003 |
| Northern Rockies | 189,674 | 0.66 | 0.002 | 179,452 | 0.63 | 0.002 | 10,000 | 0.67 | 0.006 |
| Southern Rockies | 187,211 | 0.65 | 0.002 | 179,399 | 0.64 | 0.001 | 10,000 | 0.66 | 0.005 |
| Wyoming Basin | 188,034 | 0.57 | 0.002 | 179,913 | 0.60 | 0.002 | 10,000 | 0.69 | 0.005 |

### Appendix B

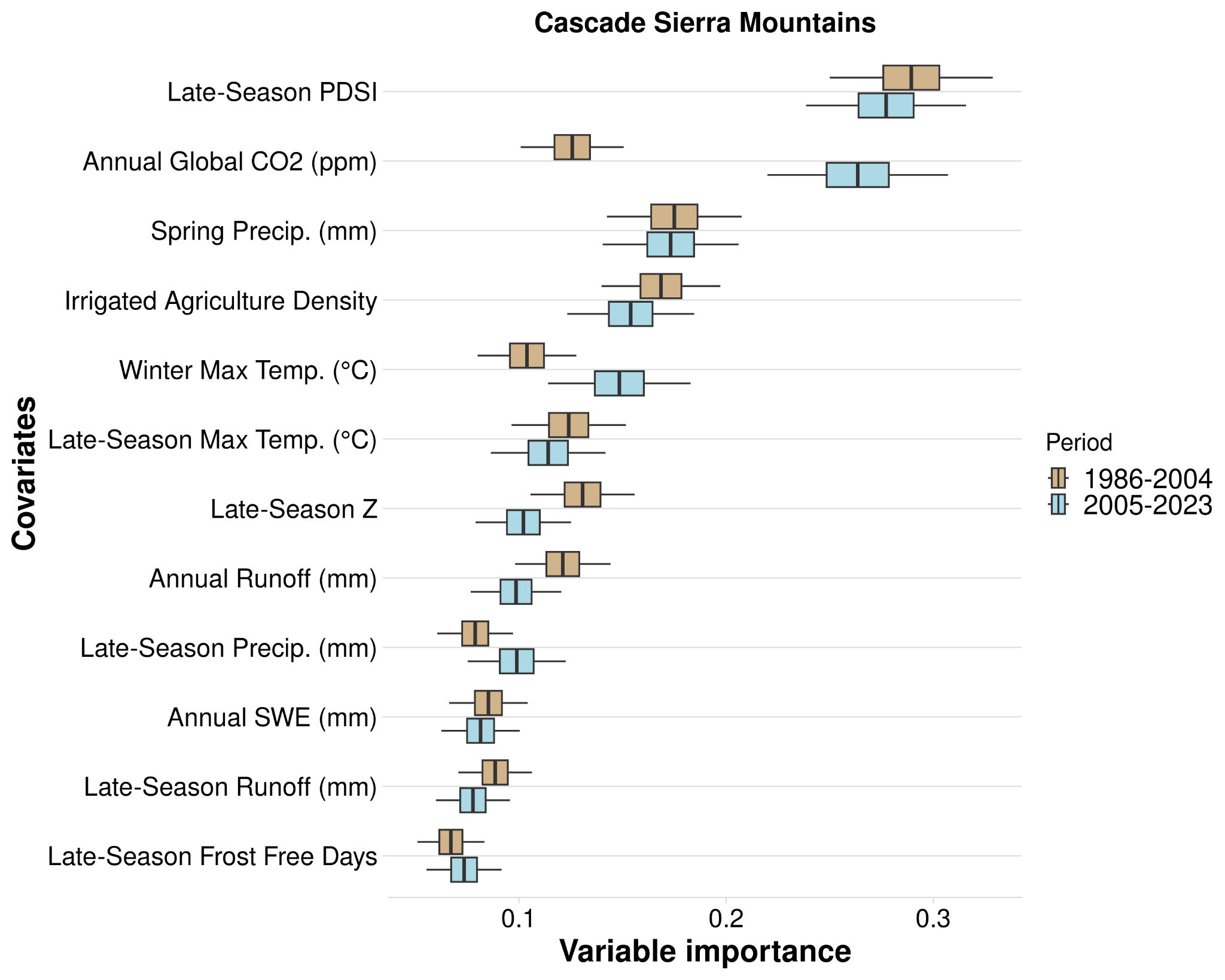

Figure B.1. Variable importance (VIMP) of variables predicting mesic resource productivity across the Cascade Sierra Mountains region during 1984-2004 (P1) and 2005-2024 (P2) using the temporal random forest analysis. Variable importance was assessed using the Breiman-Cutler permutation method, with higher values indicating greater predictive power. Center black lines represent median importance values from 100 subsampled VIMP scores; boxes cover the 25th to 75th percentiles; whiskers extend to 95% confidence intervals. Variable importance is standardized by dividing by the variance of Y. Variables are ranked by their mean importance across both periods. Variables shown as densities are measured in hectares per 500-meter radius.

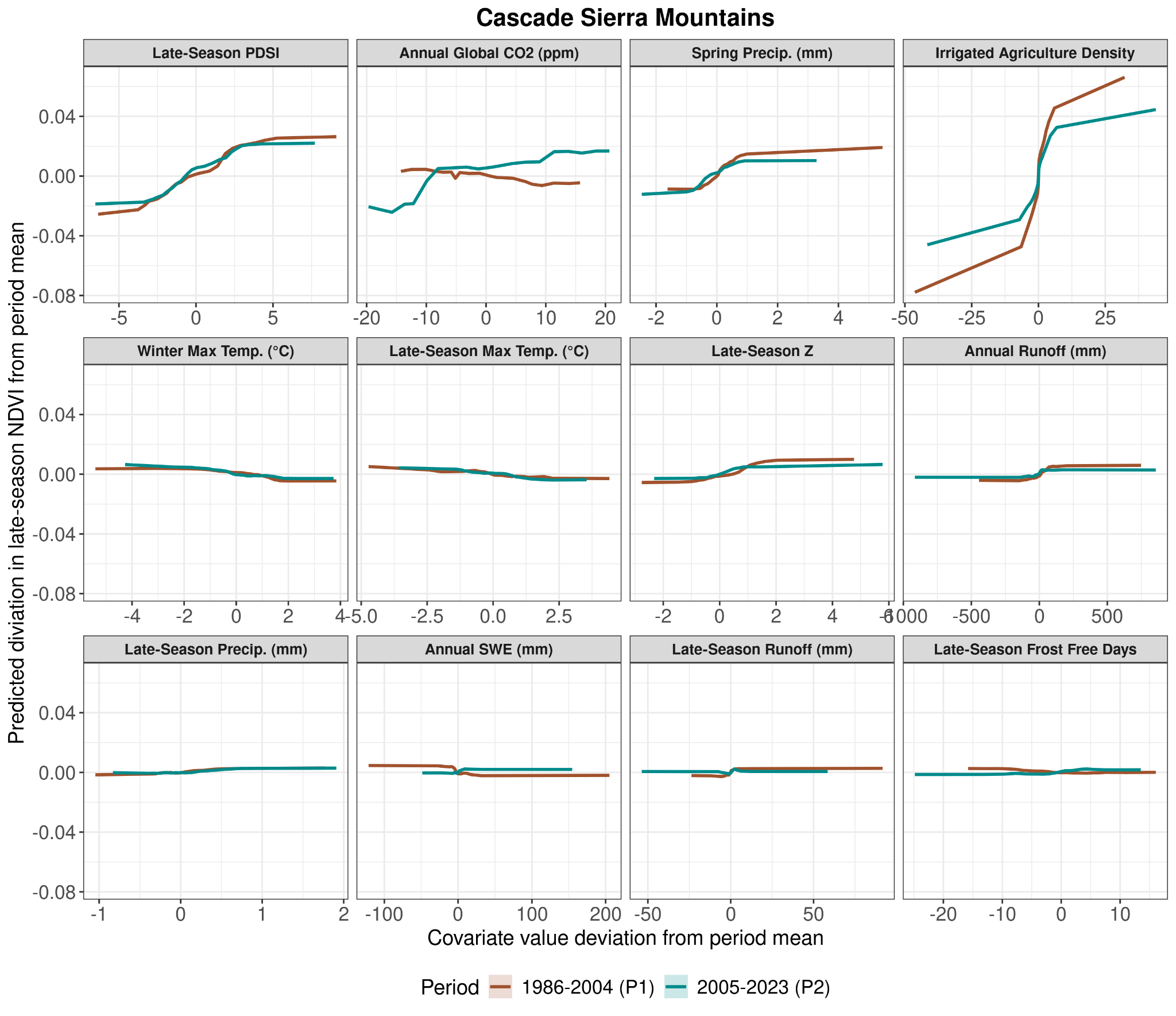

Figure B.2. Partial dependency plots for each variable used in the temporal model on predicting mesic resource productivity in the Cascade Sierra Mountains region. Y-axis values are the predicted deviation of late-season NDVI from the period mean, and the X-axis shows variable deviation from its period mean. Values below 0 indicate below-average values for the time period, and values above 0 are above-average values for both the Y and X axes. Different bar colors for P1 (brown; 1986-2004) and P2 (blue; 2005-2023) indicate differences in the effects of variables on mesic productivity over time. Variables shown as densities are measured in hectares per 500-meter radius.

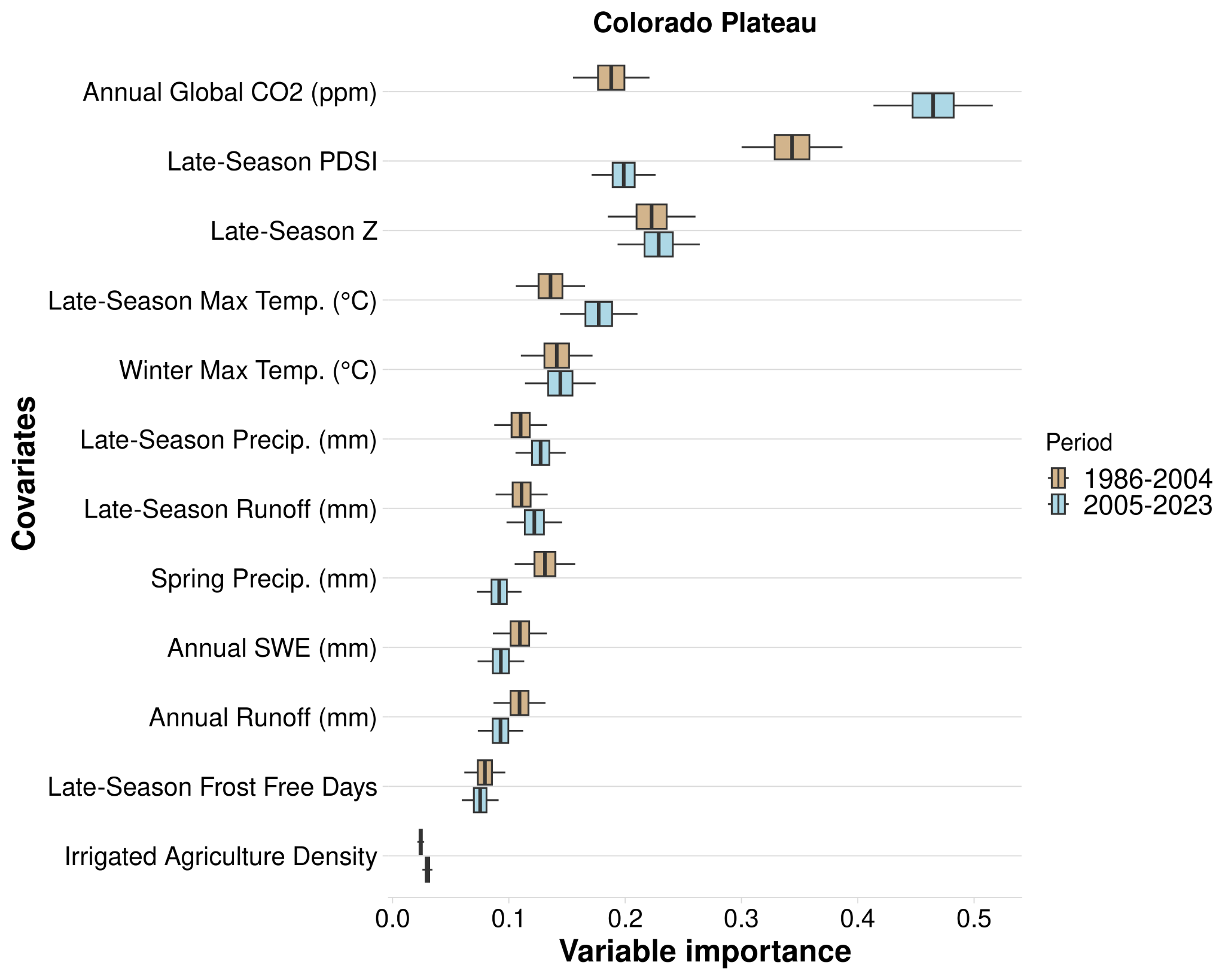

Figure B.3. Variable importance (VIMP) of variables predicting mesic resource productivity across the Colorado Plateau region during 1984-2004 (P1) and 2005-2024 (P2) using the temporal random forest analysis. Variable importance was assessed using the Breiman-Cutler permutation method, with higher values indicating greater predictive power. Center black lines represent median importance values from 100 subsampled VIMP scores; boxes cover the 25th to 75th percentiles; whiskers extend to 95% confidence intervals. Variable importance is standardized by dividing by the variance of Y. Variables are ranked by their mean importance across both periods. Variables shown as densities are measured in hectares per 500-meter radius.
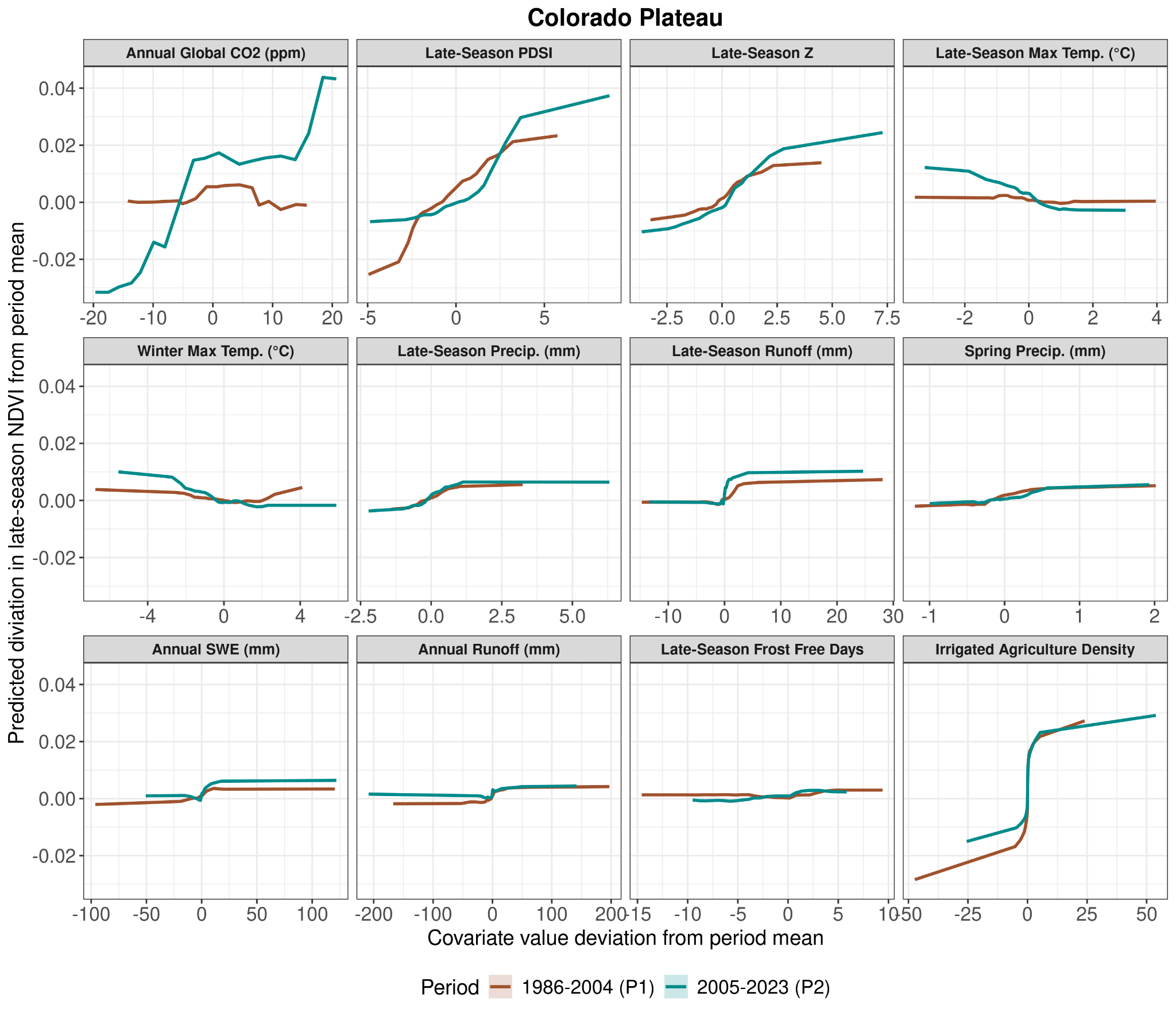
Figure B.4. Partial dependency plots for each variable used in the temporal model on predicting mesic resource productivity in the Colorado Plateau region. Y-axis values are the predicted deviation of late-season NDVI from the period mean, and the X-axis shows variable deviation from its period mean. Values below 0 indicate below-average values for the time period, and values above 0 are above-average values for both the Y and X axes. Different bar colors for P1 (brown; 1986-2004) and P2 (blue; 2005-2023) indicate differences in the effects of variables on mesic productivity over time. Variables shown as densities are measured in hectares per 500-meter radius.

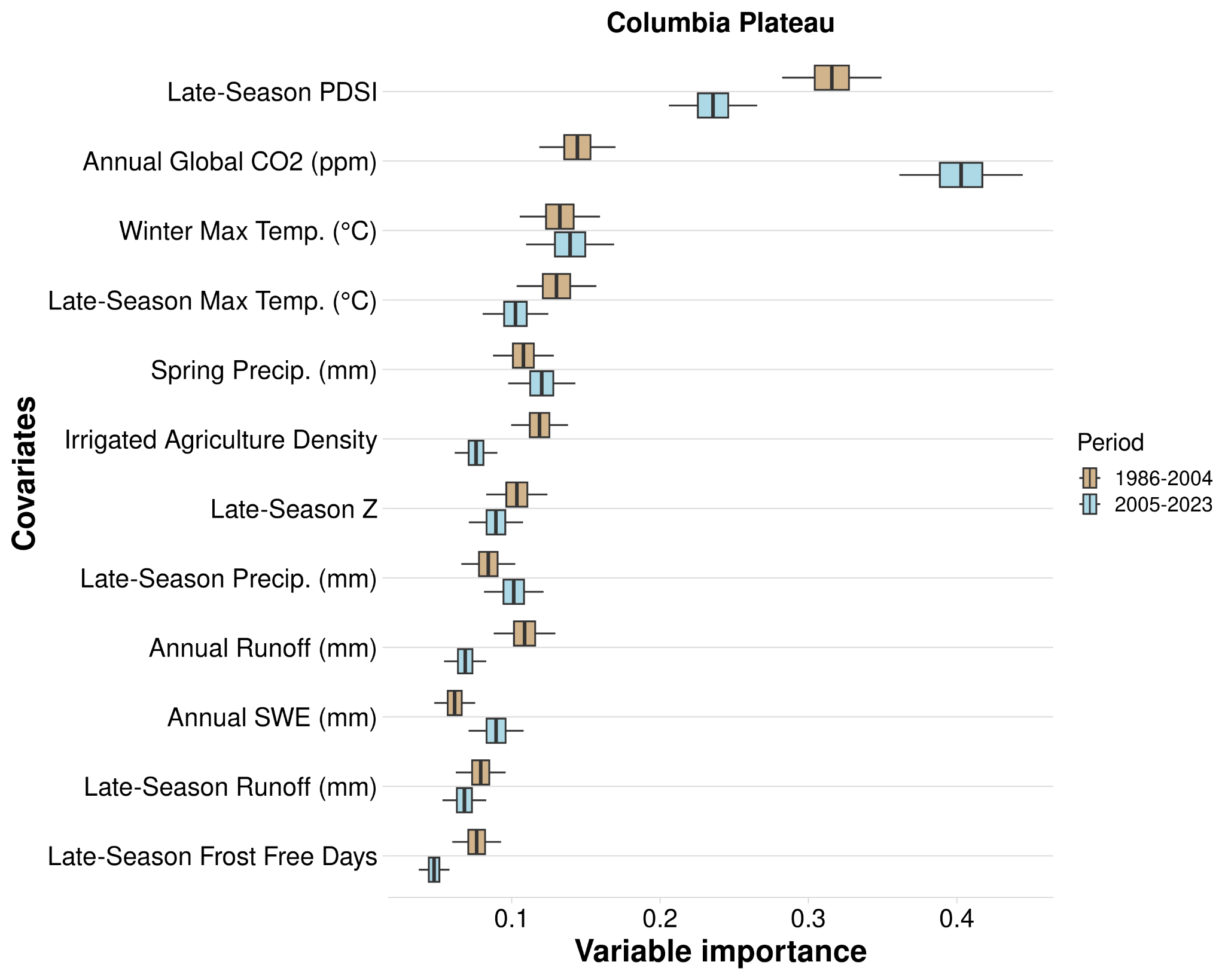

Figure B.5. Variable importance (VIMP) of variables predicting mesic resource productivity across the Columbia Plateau region during 1984-2004 (P1) and 2005-2024 (P2) using the temporal random forest analysis. Variable importance was assessed using the Breiman-Cutler permutation method, with higher values indicating greater predictive power. Center black lines represent median importance values from 100 subsampled VIMP scores; boxes cover the 25th to 75th percentiles; whiskers extend to 95% confidence intervals. Variable importance is standardized by dividing by the variance of Y. Variables are ranked by their mean importance across both periods. Variables shown as densities are measured in hectares per 500-meter radius.

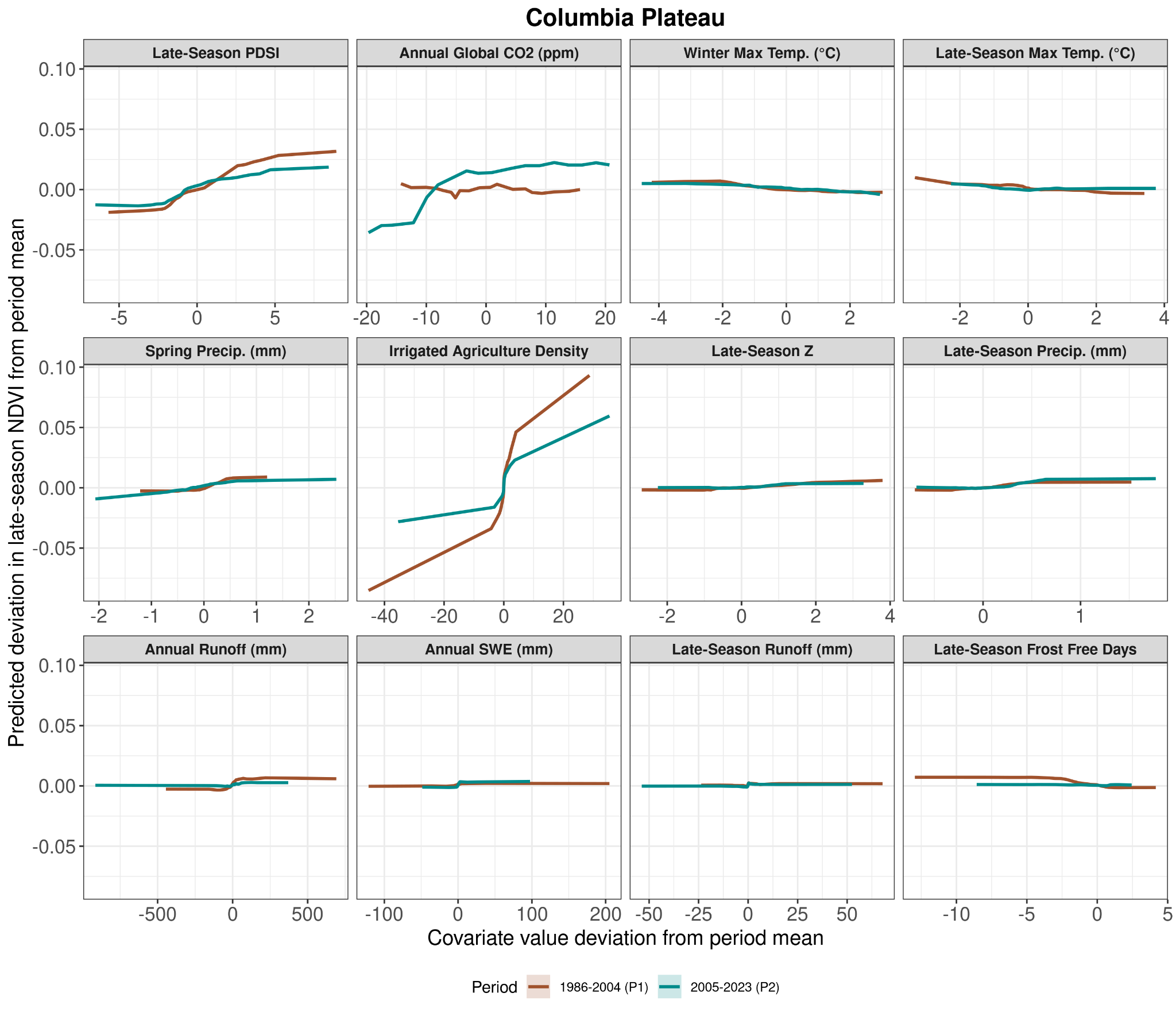

Figure B.6. Partial dependency plots for each variable used in the temporal model on predicting mesic resource productivity in the Columbia Plateau region. Y-axis values are the predicted deviation of late-season NDVI from the period mean, and the X-axis shows variable deviation from its period mean. Values below 0 indicate below-average values for the time period, and values above 0 are above-average values for both the Y and X axes. Different bar colors for P1 (brown; 1986-2004) and P2 (blue; 2005-2023) indicate differences in the effects of variables on mesic productivity over time. Variables shown as densities are measured in hectares per 500-meter radius.

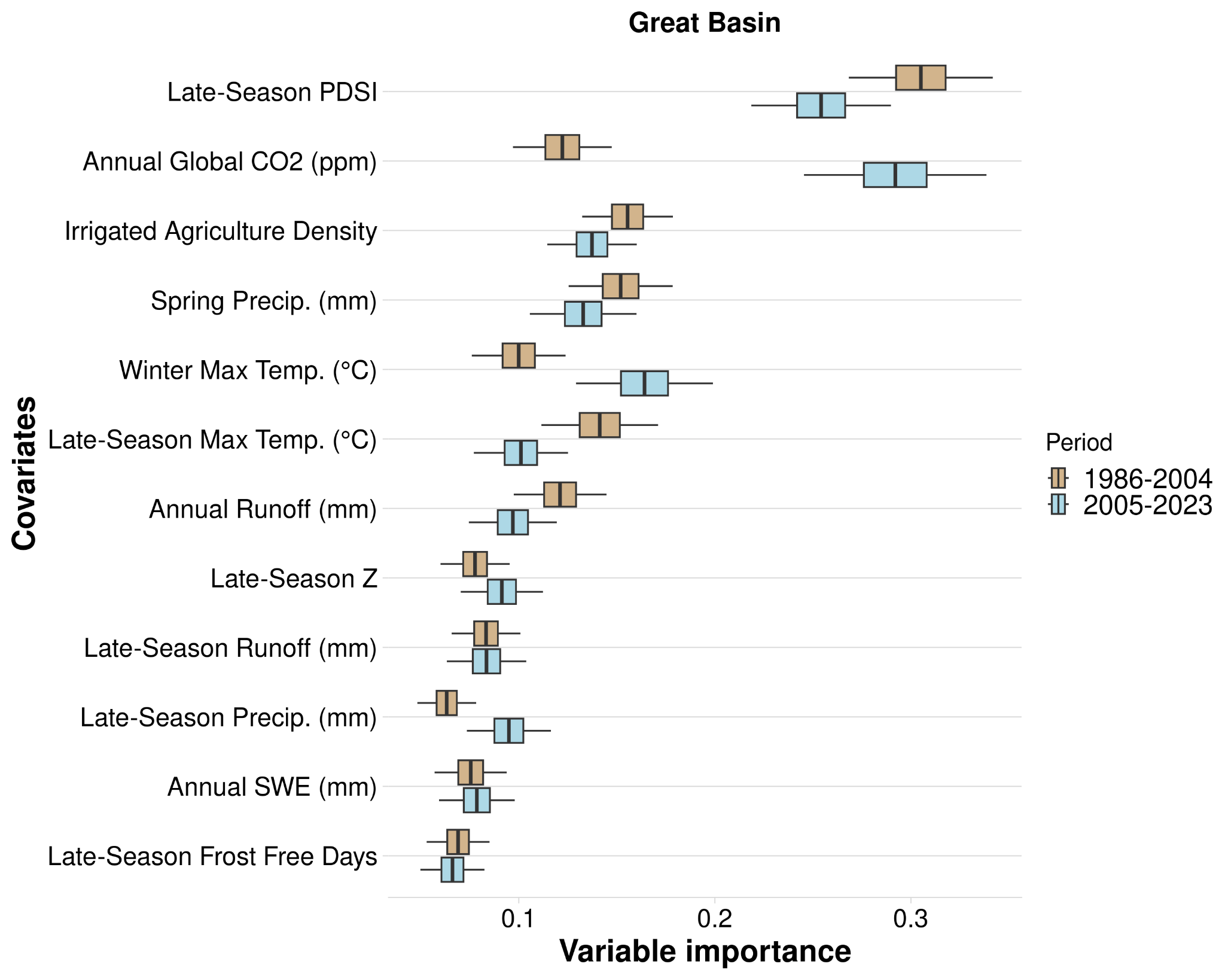

Figure B.7. Variable importance (VIMP) of variables predicting mesic resource productivity across the Great Basin region during 1984-2004 (P1) and 2005-2024 (P2) using the temporal random forest analysis. Variable importance was assessed using the Breiman-Cutler permutation method, with higher values indicating greater predictive power. Center black lines represent median importance values from 100 subsampled VIMP scores; boxes cover the 25th to 75th percentiles; whiskers extend to 95% confidence intervals. Variable importance is standardized by dividing by the variance of Y. Variables are ranked by their mean importance across both periods. Variables shown as densities are measured in hectares per 500-meter radius.

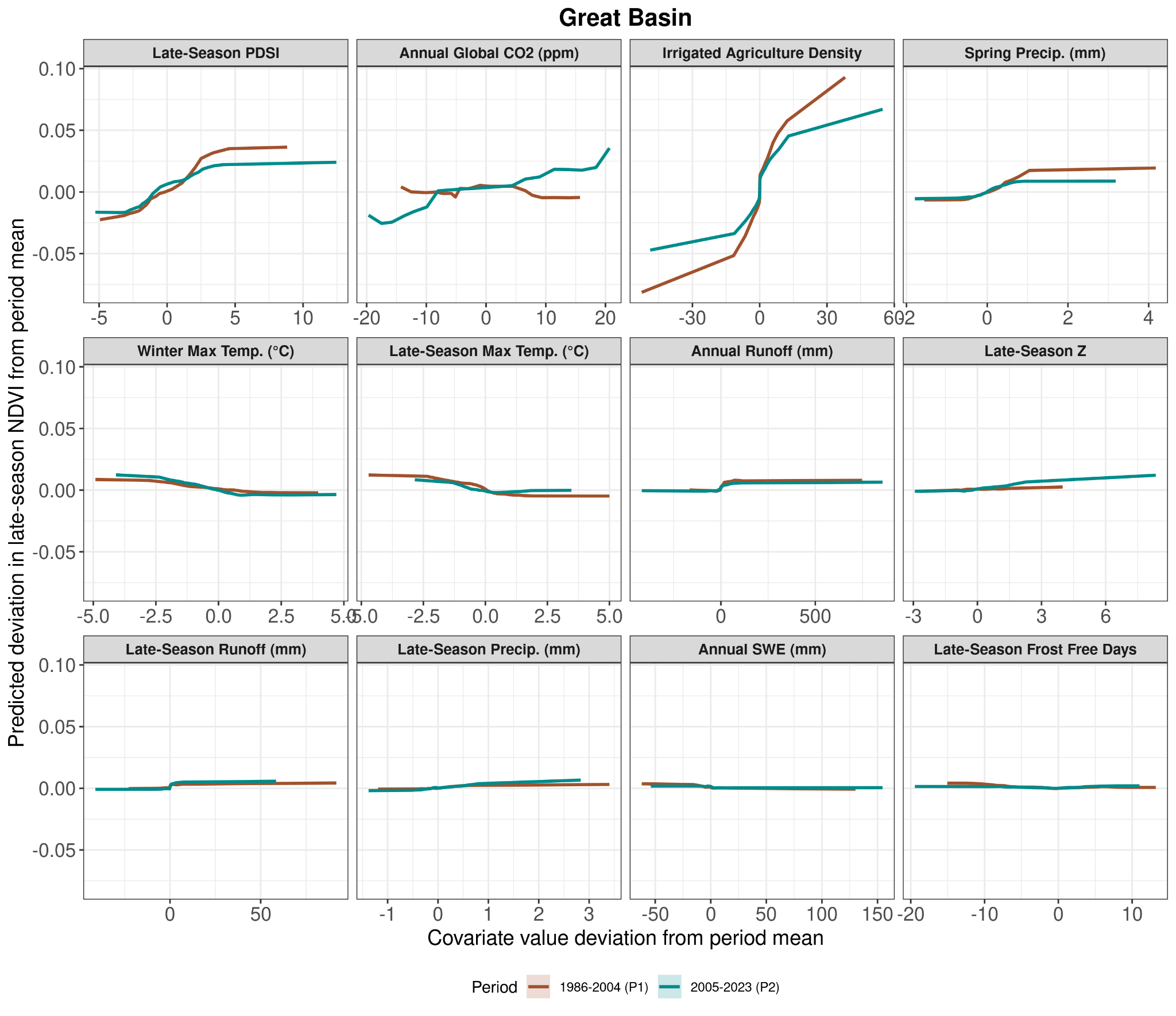

Figure B.8. Partial dependency plots for each variable used in the temporal model on predicting mesic resource productivity in the Great Basin region. Y-axis values are the predicted deviation of late-season NDVI from the period mean, and the X-axis shows variable deviation from its period mean. Values below 0 indicate below-average values for the time period, and values above 0 are above-average values for both the Y and X axes. Different bar colors for P1 (brown; 1986-2004) and P2 (blue; 2005-2023) indicate differences in the effects of variables on mesic productivity over time. Variables shown as densities are measured in hectares per 500-meter radius.

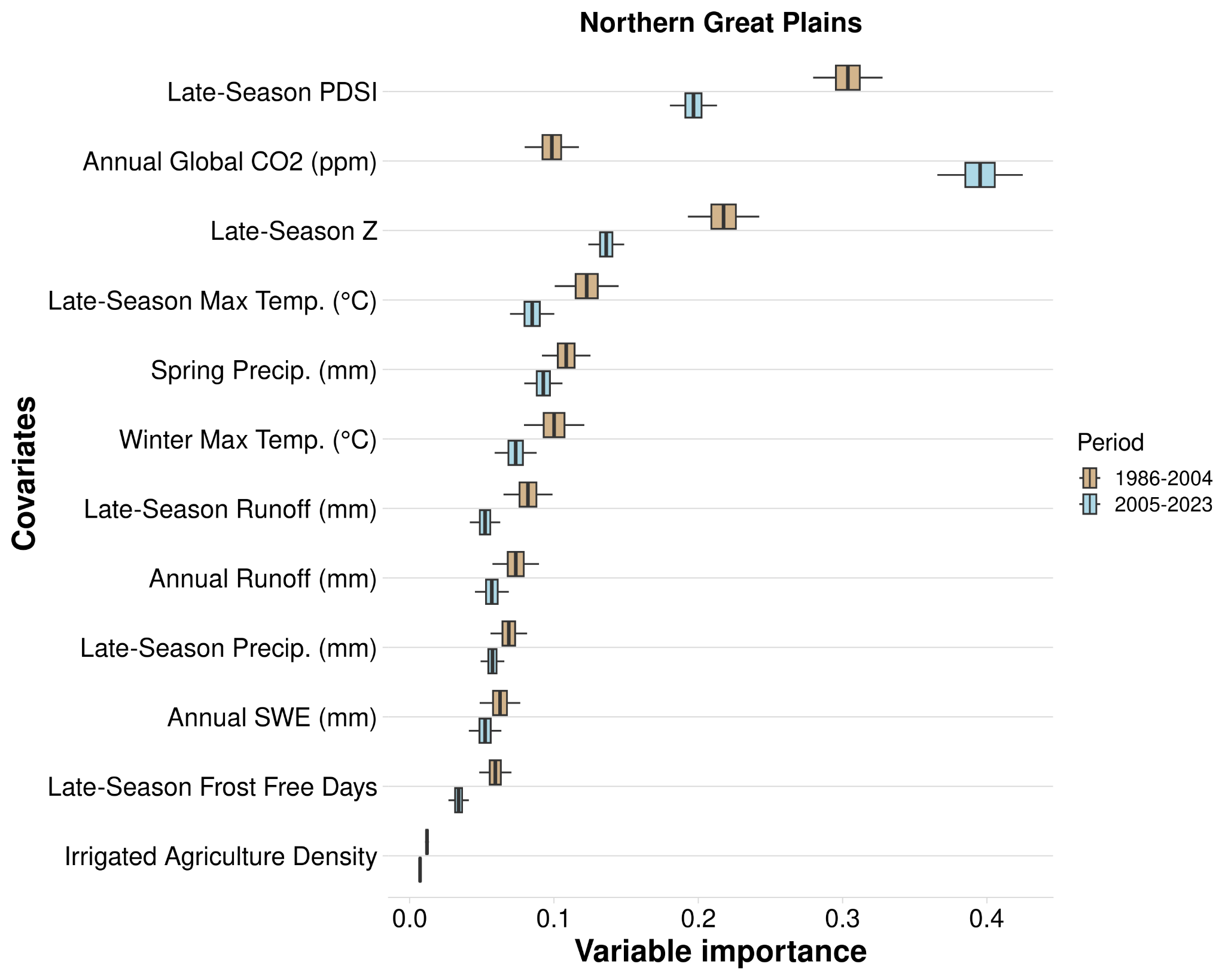

Figure B.9. Variable importance (VIMP) of variables predicting mesic resource productivity across the Northern Great Plains region during 1984-2004 (P1) and 2005-2024 (P2) using the temporal random forest analysis. Variable importance was assessed using the Breiman-Cutler permutation method, with higher values indicating greater predictive power. Center black lines represent median importance values from 100 subsampled VIMP scores; boxes cover the 25th to 75th percentiles; whiskers extend to 95% confidence intervals. Variable importance is standardized by dividing by the variance of Y. Variables are ranked by their mean importance across both periods. Variables shown as densities are measured in hectares per 500-meter radius.

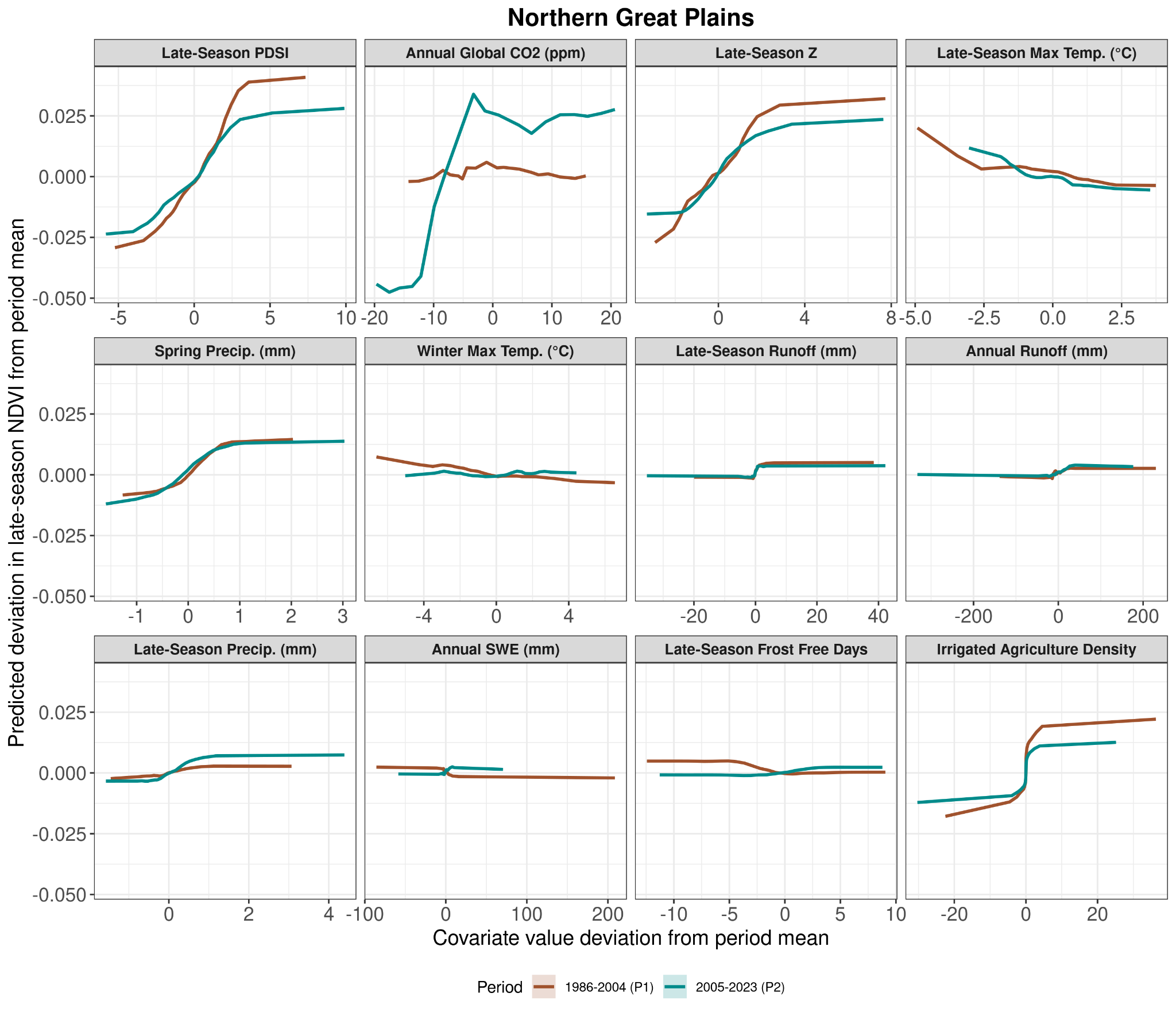

Figure B.10. Partial dependency plots for each variable used in the temporal model on predicting mesic resource productivity in the Northern Great Plains region. Y-axis values are the predicted deviation of late-season NDVI from the period mean, and the X-axis shows variable deviation from its period mean. Values below 0 indicate below-average values for the time period, and values above 0 are above-average values for both the Y and X axes. Different bar colors for P1 (brown; 1986-2004) and P2 (blue; 2005-2023) indicate differences in the effects of variables on mesic productivity over time. Variables shown as densities are measured in hectares per 500-meter radius.

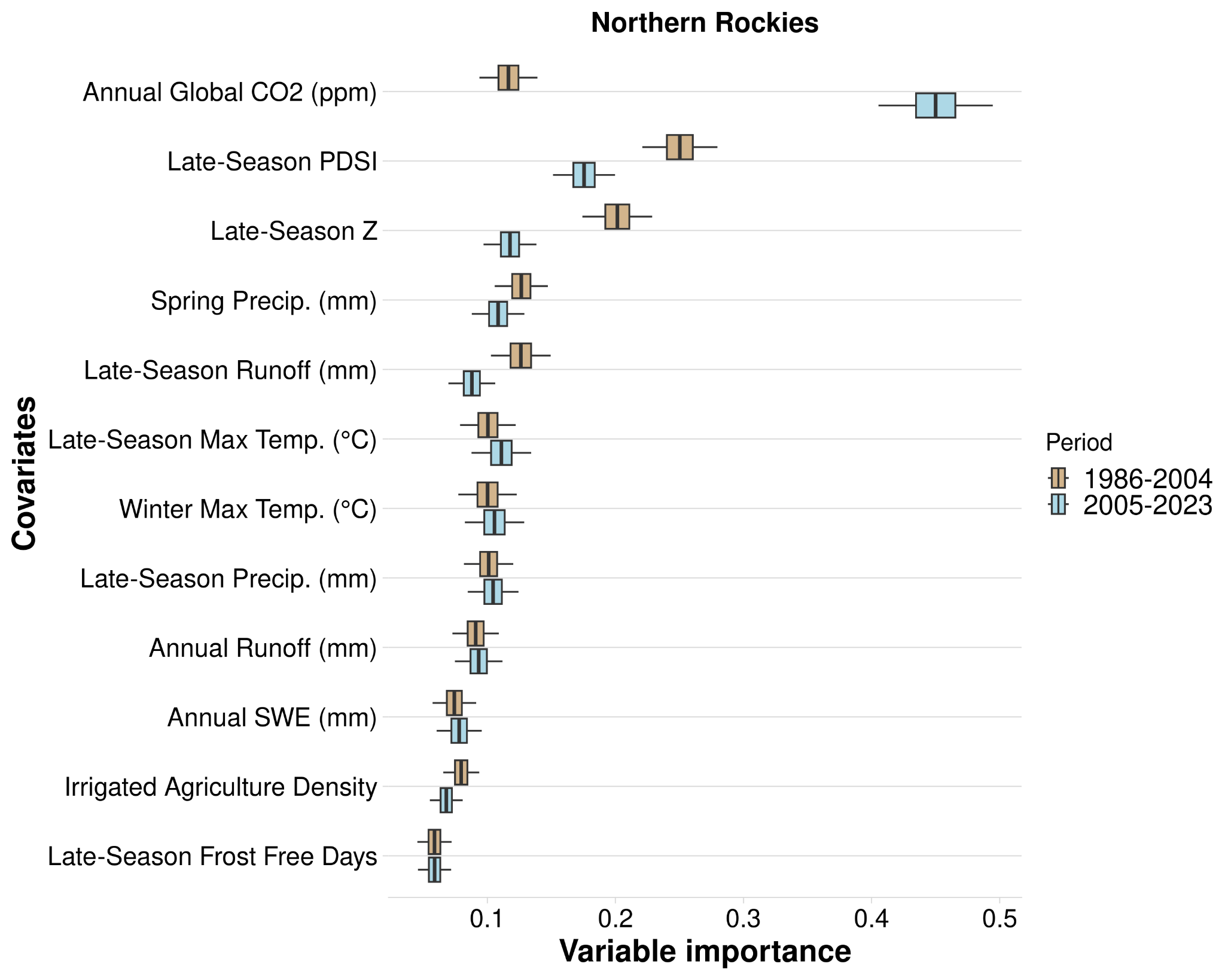

Figure B.11. Variable importance (VIMP) of variables predicting mesic resource productivity across the Northern Rockies region during 1984-2004 (P1) and 2005-2024 (P2) using the temporal random forest analysis. Variable importance was assessed using the Breiman-Cutler permutation method, with higher values indicating greater predictive power. Center black lines represent median importance values from 100 subsampled VIMP scores; boxes cover the 25th to 75th percentiles; whiskers extend to 95% confidence intervals. Variable importance is standardized by dividing by the variance of Y. Variables are ranked by their mean importance across both periods. Variables shown as densities are measured in hectares per 500-meter radius.

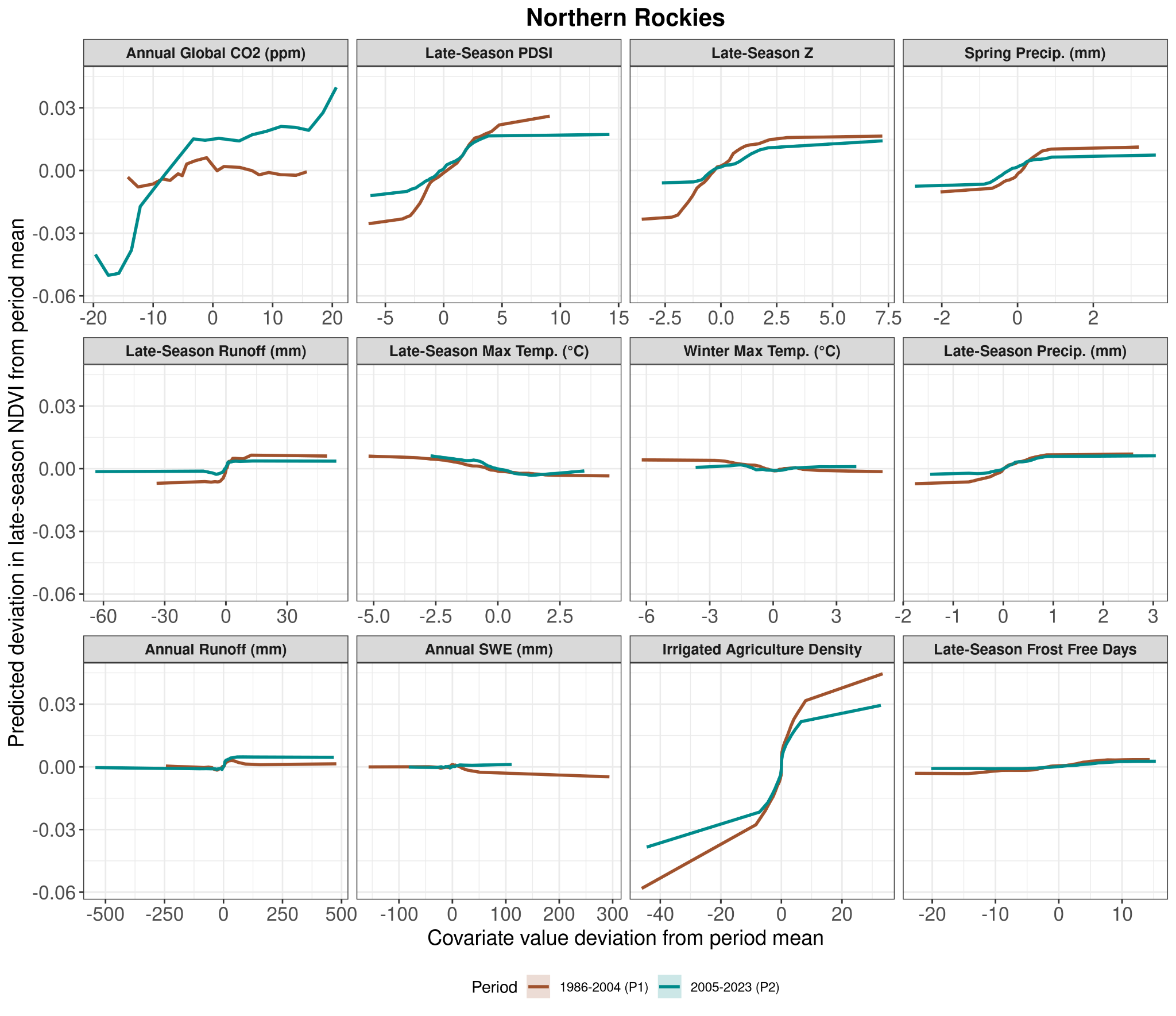

Figure B.12. Partial dependency plots for each variable used in the temporal model on predicting mesic resource productivity in the Northern Rockies region. Y-axis values are the predicted deviation of late-season NDVI from the period mean, and the X-axis shows variable deviation from its period mean. Values below 0 indicate below-average values for the time period, and values above 0 are above-average values for both the Y and X axes. Different bar colors for P1 (brown; 1986-2004) and P2 (blue; 2005-2023) indicate differences in the effects of variables on mesic productivity over time. Variables shown as densities are measured in hectares per 500-meter radius.

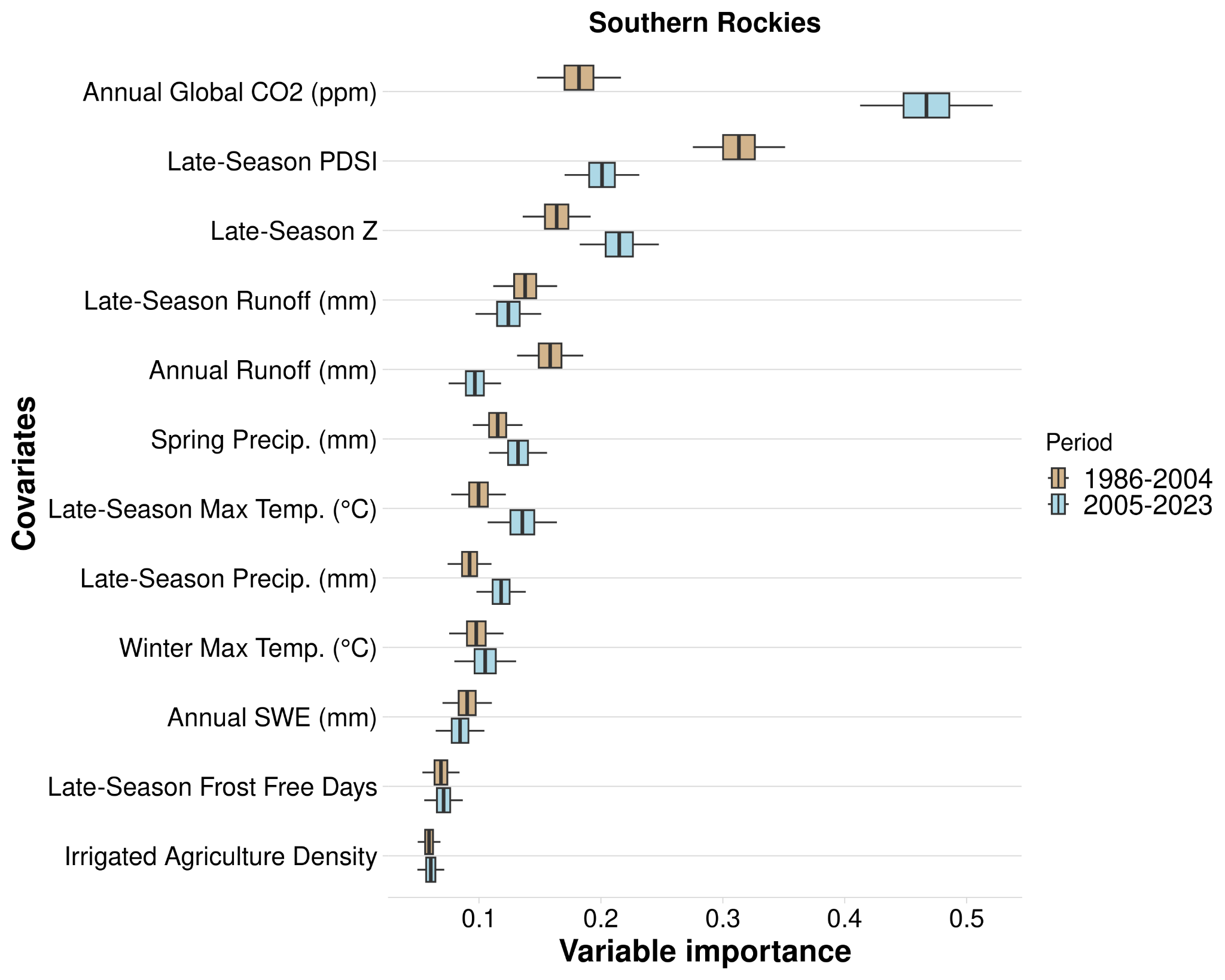

Figure B.13. Variable importance (VIMP) of variables predicting mesic resource productivity across the Southern Rockies region during 1984-2004 (P1) and 2005-2024 (P2) using the temporal random forest analysis. Variable importance was assessed using the Breiman-Cutler permutation method, with higher values indicating greater predictive power. Center black lines represent median importance values from 100 subsampled VIMP scores; boxes cover the 25th to 75th percentiles; whiskers extend to 95% confidence intervals. Variable importance is standardized by dividing by the variance of Y. Variables are ranked by their mean importance across both periods. Variables shown as densities are measured in hectares per 500-meter radius.

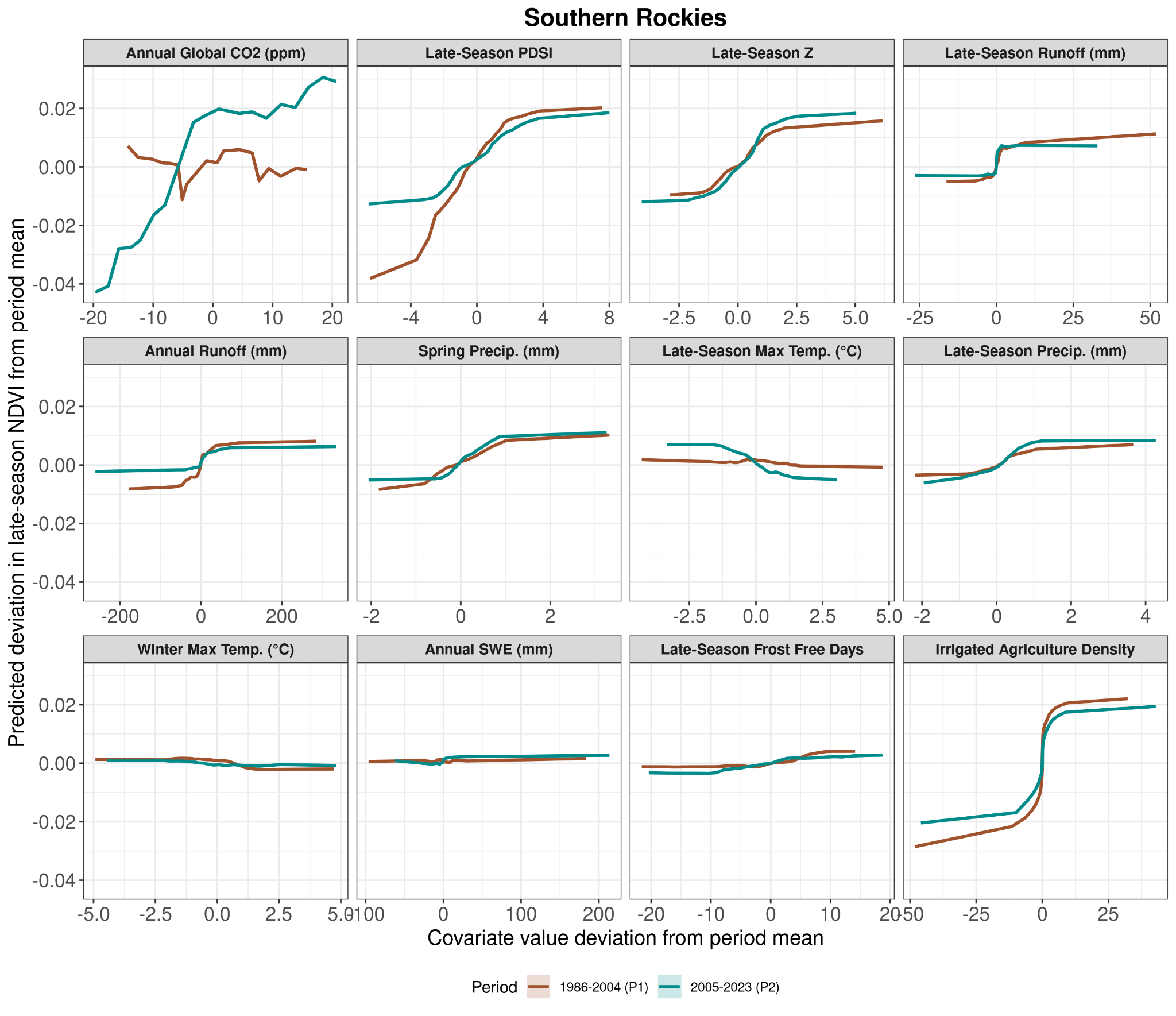

Figure B.14. Partial dependency plots for each variable used in the temporal model on predicting mesic resource productivity in the Southern Rockies region. Y-axis values are the predicted deviation of late-season NDVI from the period mean, and the X-axis shows variable deviation from its period mean. Values below 0 indicate below-average values for the time period, and values above 0 are above-average values for both the Y and X axes. Different bar colors for P1 (brown; 1986-2004) and P2 (blue; 2005-2023) indicate differences in the effects of variables on mesic productivity over time. Variables shown as densities are measured in hectares per 500-meter radius.

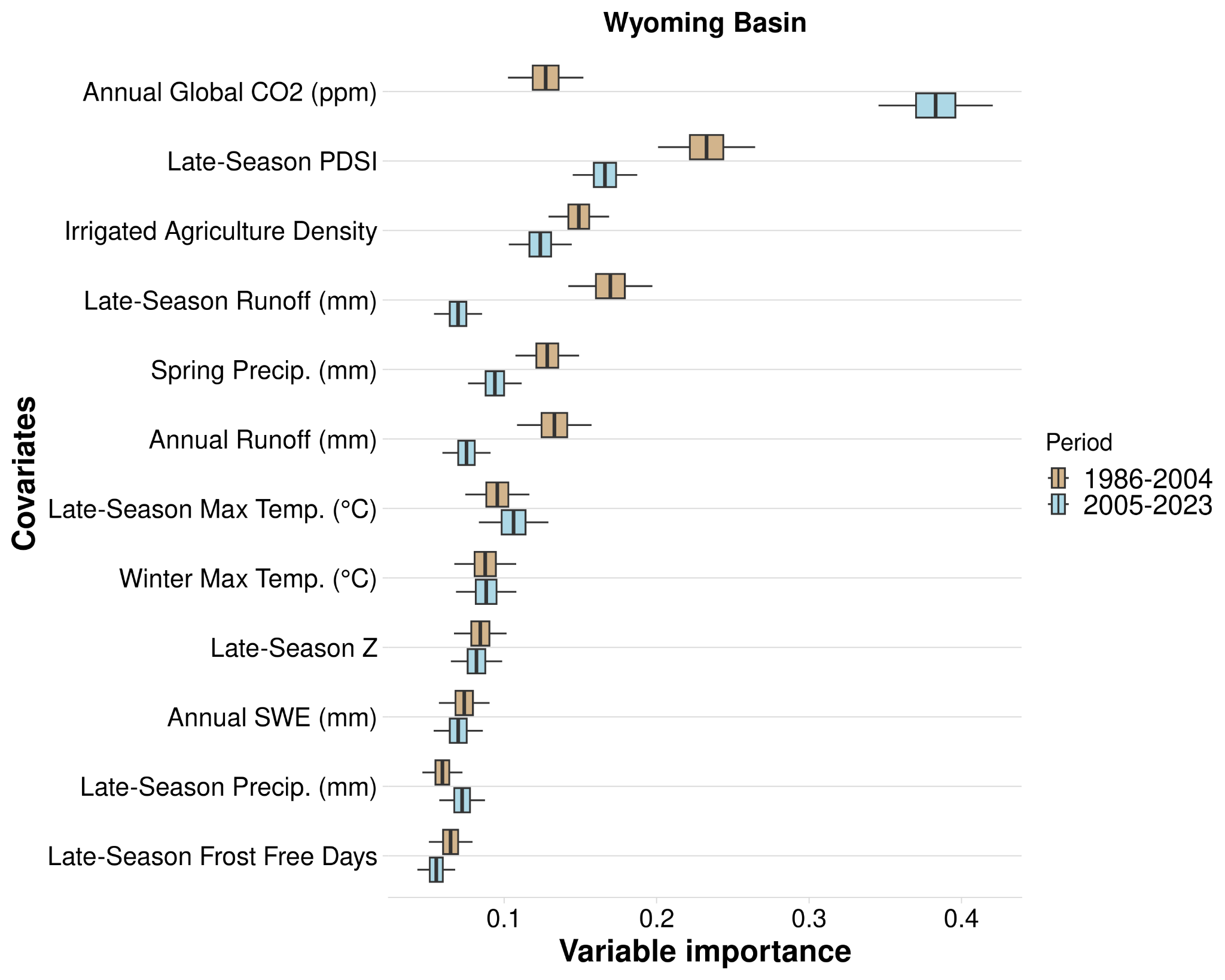

Figure B.15. Variable importance (VIMP) of variables predicting mesic resource productivity across the Wyoming Basin region during 1984-2004 (P1) and 2005-2024 (P2) using the temporal random forest analysis. Variable importance was assessed using the Breiman-Cutler permutation method, with higher values indicating greater predictive power. Center black lines represent median importance values from 100 subsampled VIMP scores; boxes cover the 25th to 75th percentiles; whiskers extend to 95% confidence intervals. Variable importance is standardized by dividing by the variance of Y. Variables are ranked by their mean importance across both periods. Variables shown as densities are measured in hectares per 500-meter radius.

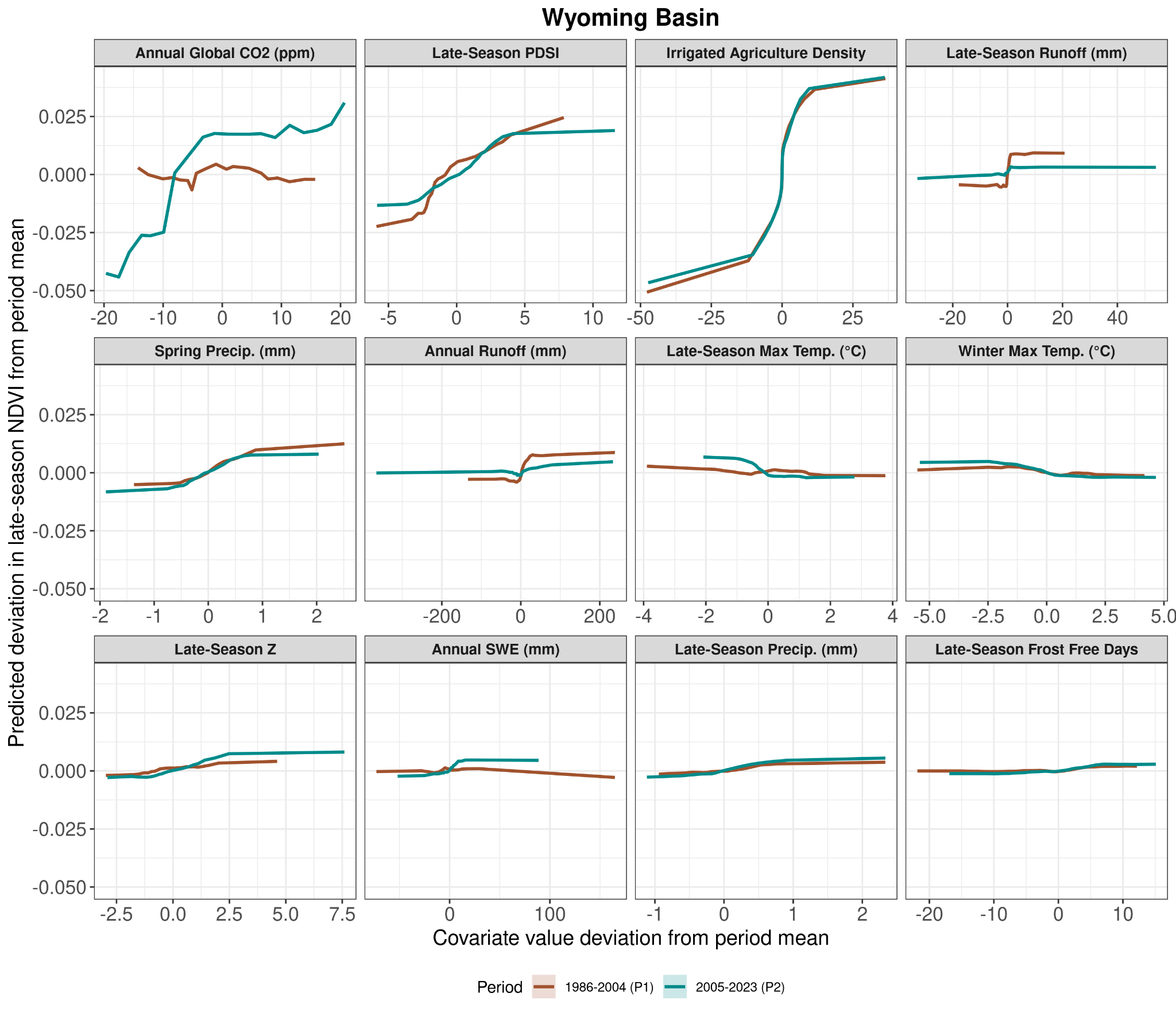

Figure B.16. Partial dependency plots for each variable used in the temporal model on predicting mesic resource productivity in the Wyoming Basin region. Y-axis values are the predicted deviation of late-season NDVI from the period mean, and the X-axis shows variable deviation from its period mean. Values below 0 indicate below-average values for the time period, and values above 0 are above-average values for both the Y and X axes. Different bar colors for P1 (brown; 1986-2004) and P2 (blue; 2005-2023) indicate differences in the effects of variables on mesic productivity over time. Variables shown as densities are measured in hectares per 500-meter radius.

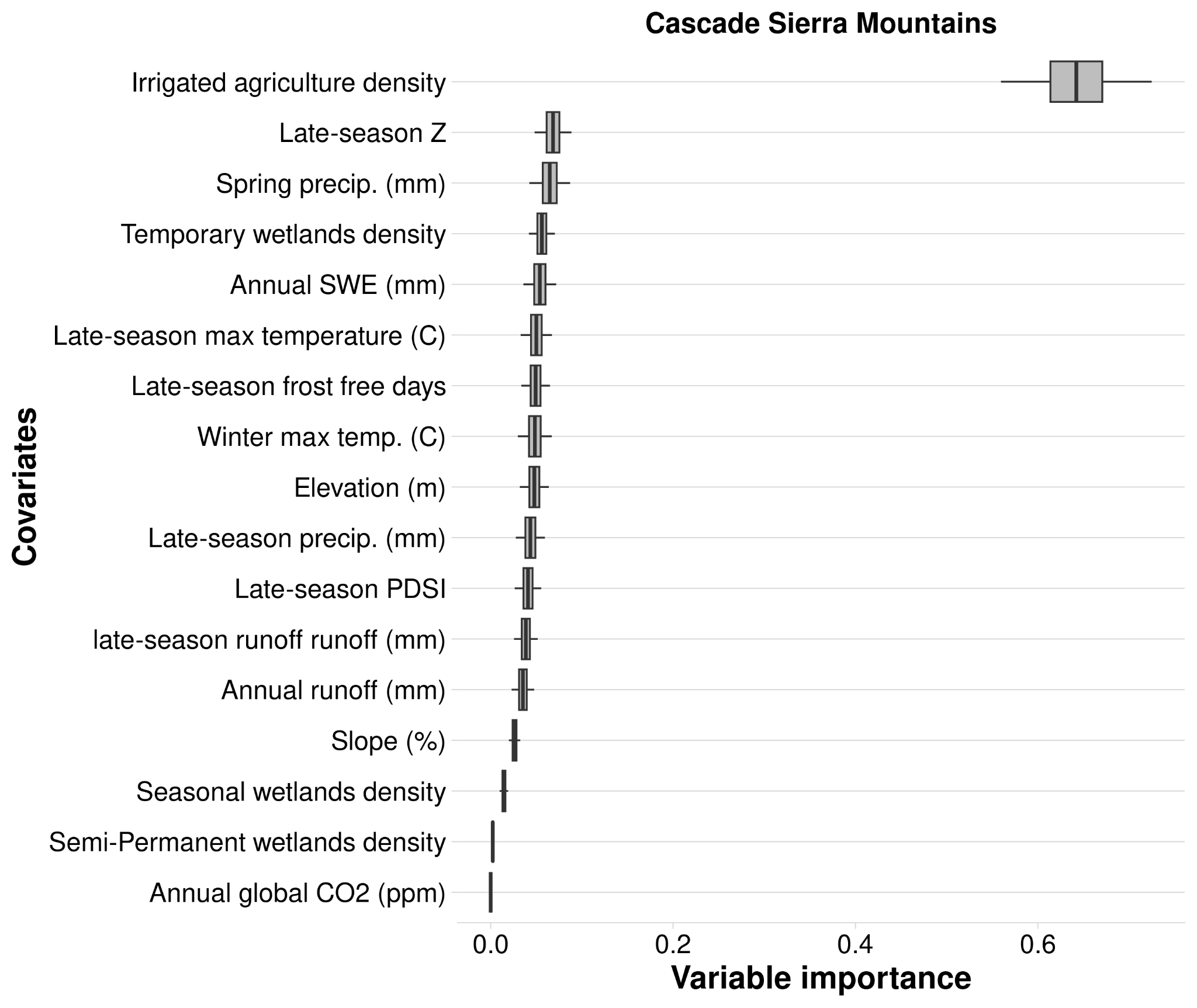

Figure B.17. Variable importance (VIMP) of variables predicting mesic resource productivity across the Cascade Sierra Mountains region from 1986 to 2023 using the spatial random forest analysis. Variable importance was assessed using the Breiman-Cutler permutation method, with higher values indicating greater predictive power. Center black lines represent median importance values from 100 subsampled VIMP scores; boxes cover the 25th to 75th percentiles; whiskers extend to 95% confidence intervals. Variable importance is standardized by dividing by the variance of Y. Variables shown as densities are measured in hectares per 500-meter radius.

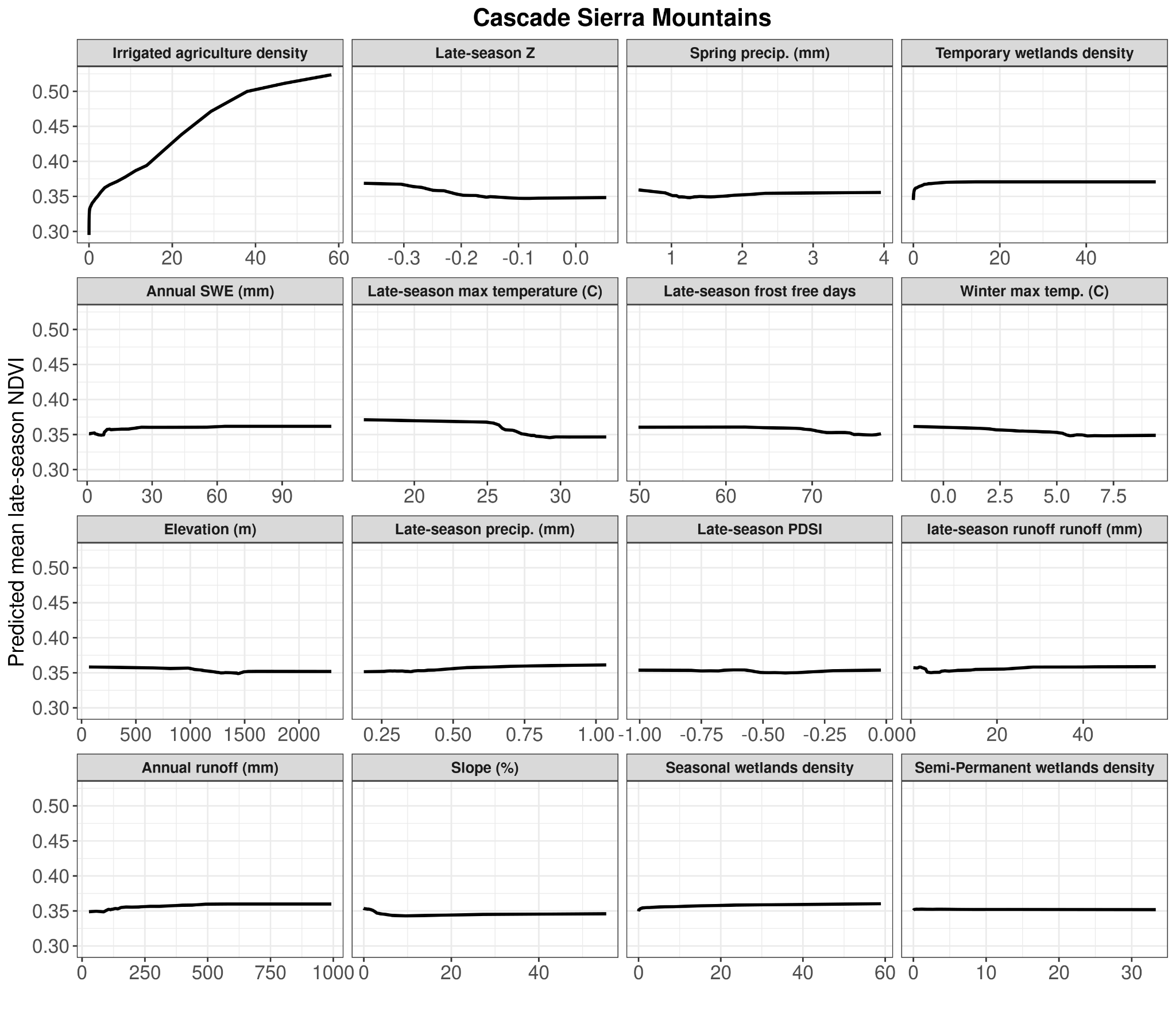

Figure B.18. Partial dependency plots for each variable used in the spatial model on predicting mesic resource productivity in the Cascade Sierra Mountains region (1986-2023). Y-axis values are the predicted mean late-season NDVI. The X-axis shows untransformed variable values. Variables shown as densities are measured in hectares per 500-meter radius.

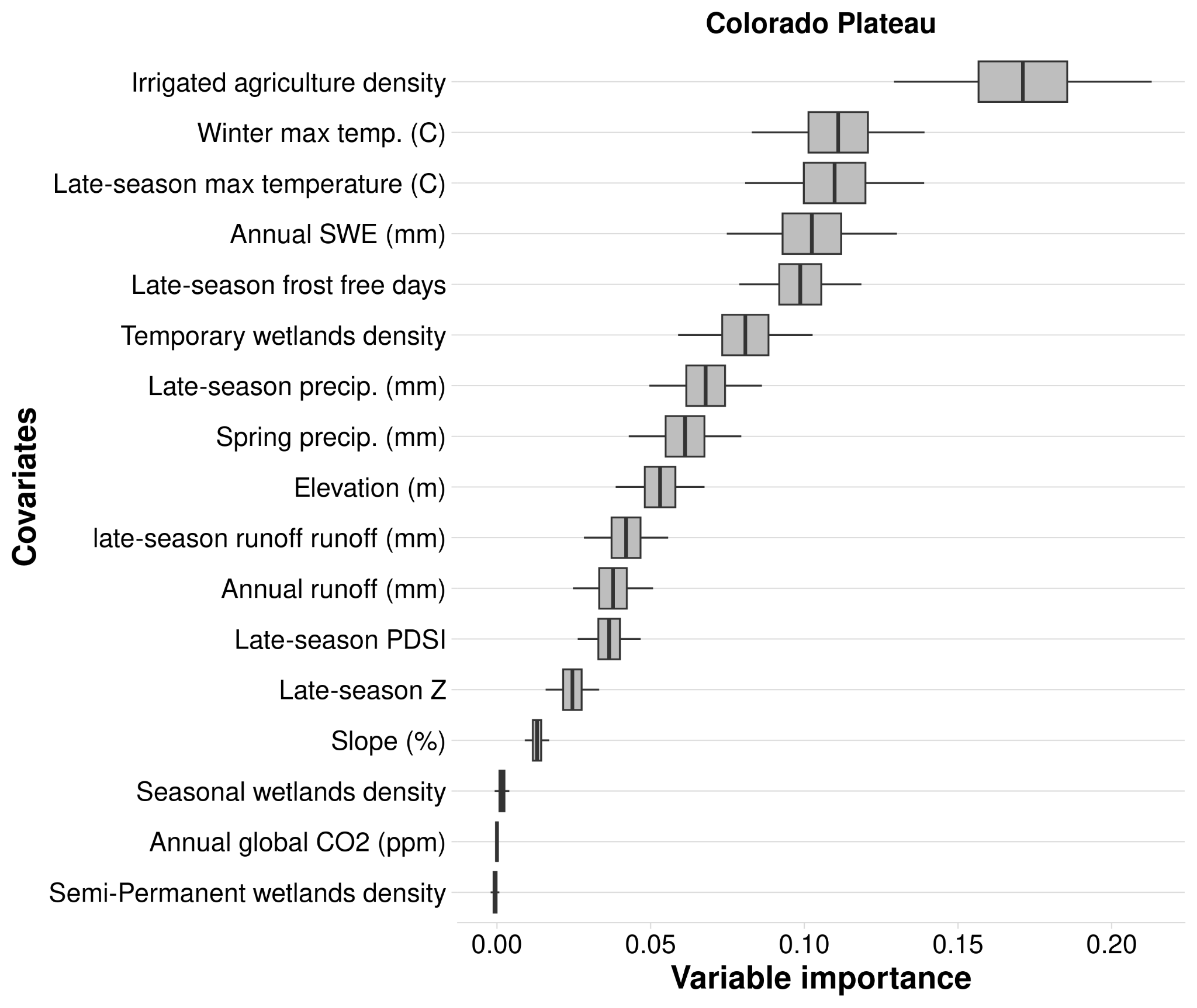

Figure B.19. Variable importance (VIMP) of variables predicting mesic resource productivity across the Colorado Plateau region from 1986 to 2023 using the spatial random forest analysis. Variable importance was assessed using the Breiman-Cutler permutation method, with higher values indicating greater predictive power. Center black lines represent median importance values from 100 subsampled VIMP scores; boxes cover the 25th to 75th percentiles; whiskers extend to 95% confidence intervals. Variable importance is standardized by dividing by the variance of Y. Variables shown as densities are measured in hectares per 500-meter radius.

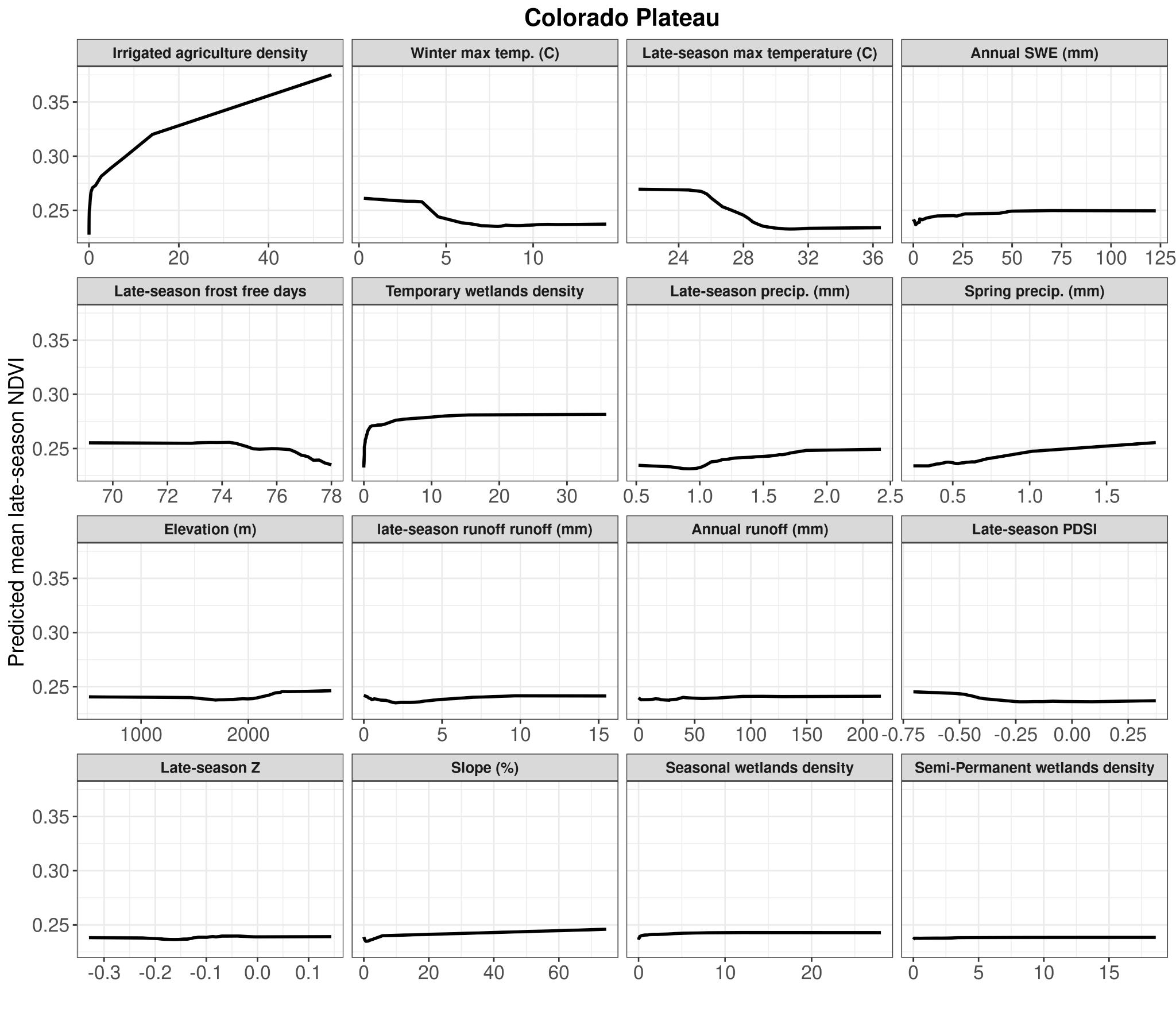

Figure B.20. Partial dependency plots for each variable used in the spatial model on predicting mesic resource productivity in the Colorado Plateau region (1986-2023). Y-axis values are the predicted mean late-season NDVI. The X-axis shows untransformed variable values. Variables shown as densities are measured in hectares per 500-meter radius.

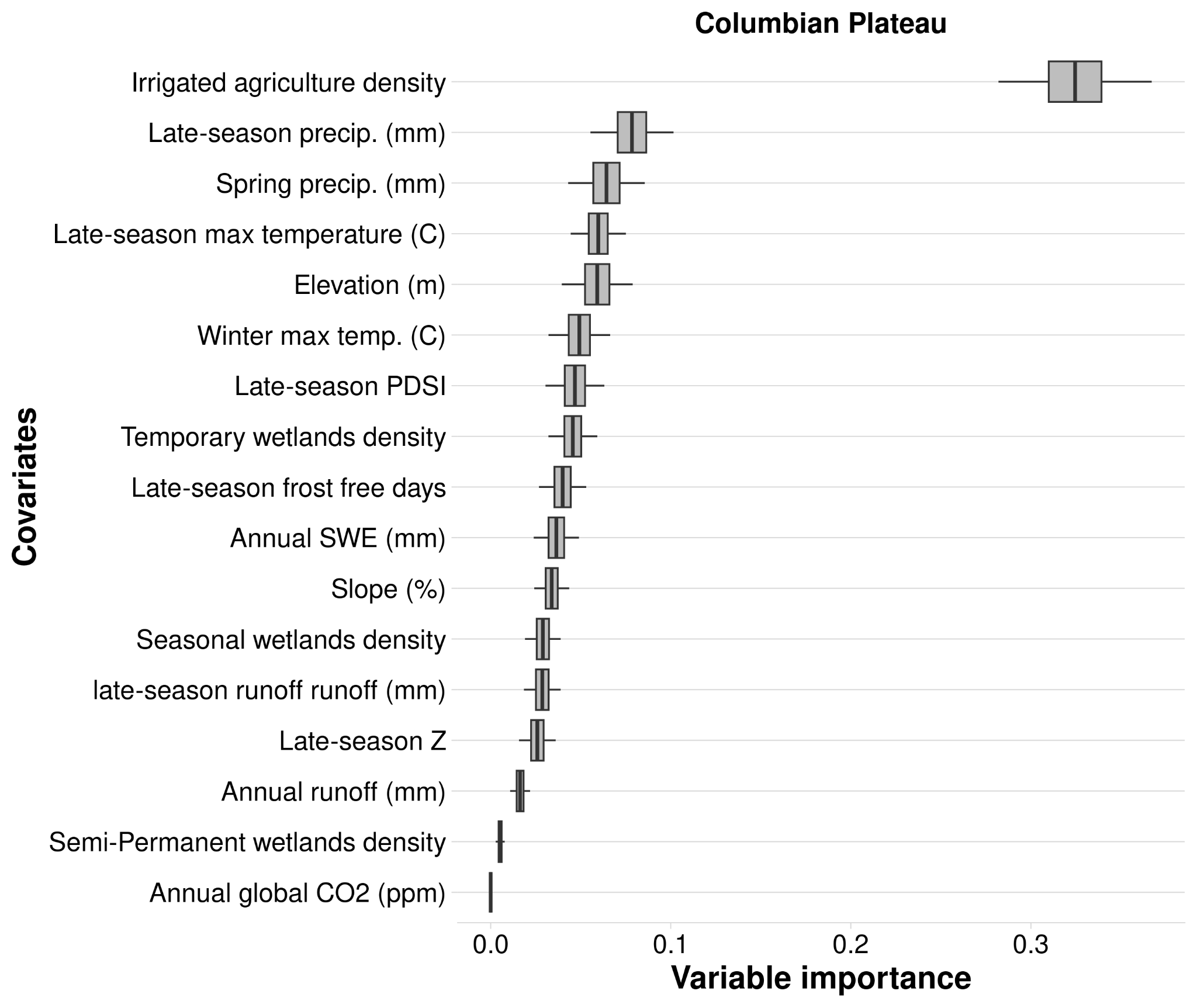

Figure B.21. Variable importance (VIMP) of variables predicting mesic resource productivity across the Columbia Plateau region from 1986 to 2023 using the spatial random forest analysis. Variable importance was assessed using the Breiman-Cutler permutation method, with higher values indicating greater predictive power. Center black lines represent median importance values from 100 subsampled VIMP scores; boxes cover the 25th to 75th percentiles; whiskers extend to 95% confidence intervals. Variable importance is standardized by dividing by the variance of Y. Variables shown as densities are measured in hectares per 500-meter radius.

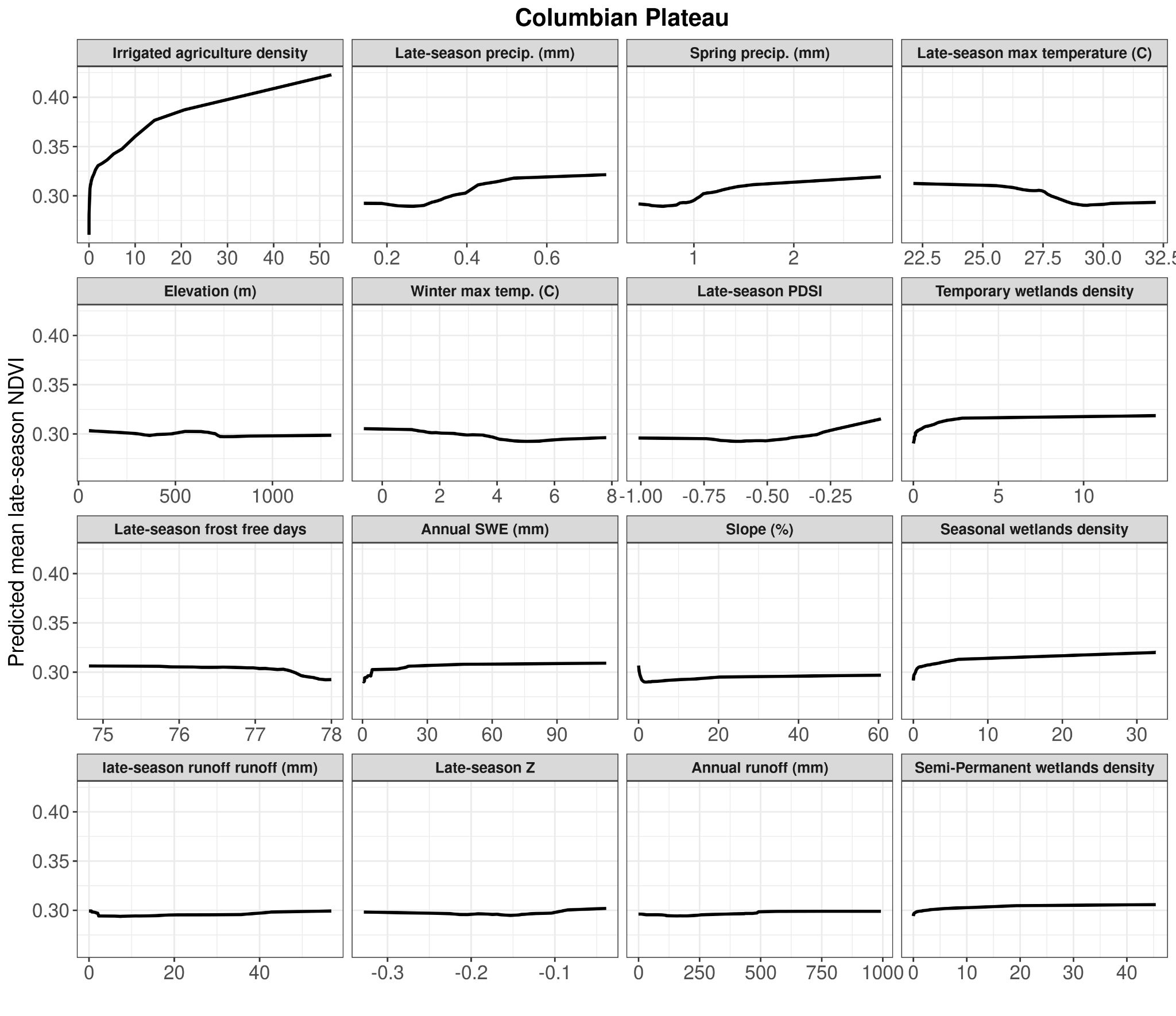

Figure B.22. Partial dependency plots for each variable used in the spatial model on predicting mesic resource productivity in the Columbia Plateau region (1986-2023). Y-axis values are the predicted mean late-season NDVI. The X-axis shows untransformed variable values. Variables shown as densities are measured in hectares per 500-meter radius.

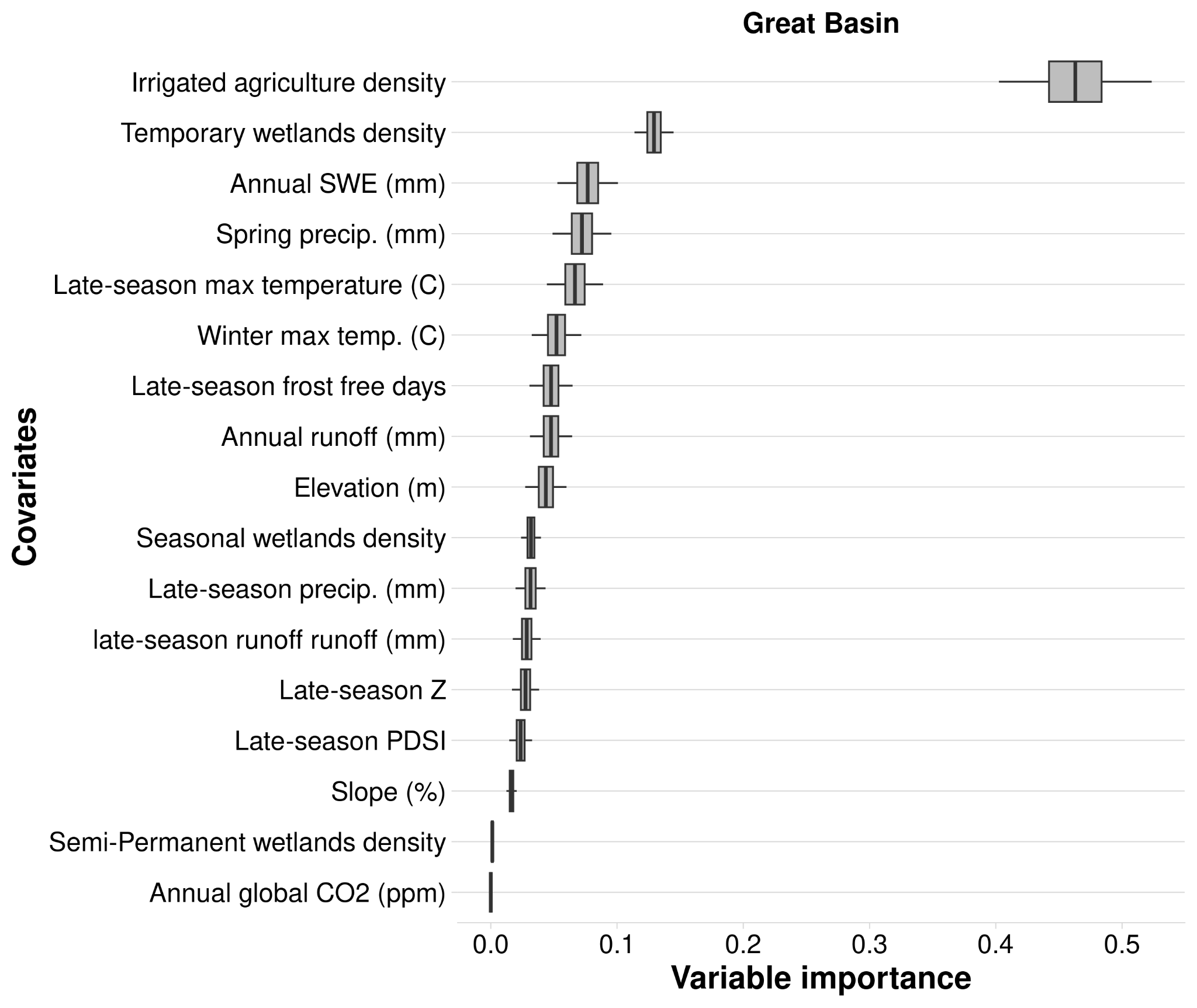

Figure B.23. Variable importance (VIMP) of variables predicting mesic resource productivity across the Great Basin region from 1986 to 2023 using the spatial random forest analysis. Variable importance was assessed using the Breiman-Cutler permutation method, with higher values indicating greater predictive power. Center black lines represent median importance values from 100 subsampled VIMP scores; boxes cover the 25th to 75th percentiles; whiskers extend to 95% confidence intervals. Variable importance is standardized by dividing by the variance of Y. Variables shown as densities are measured in hectares per 500-meter radius.

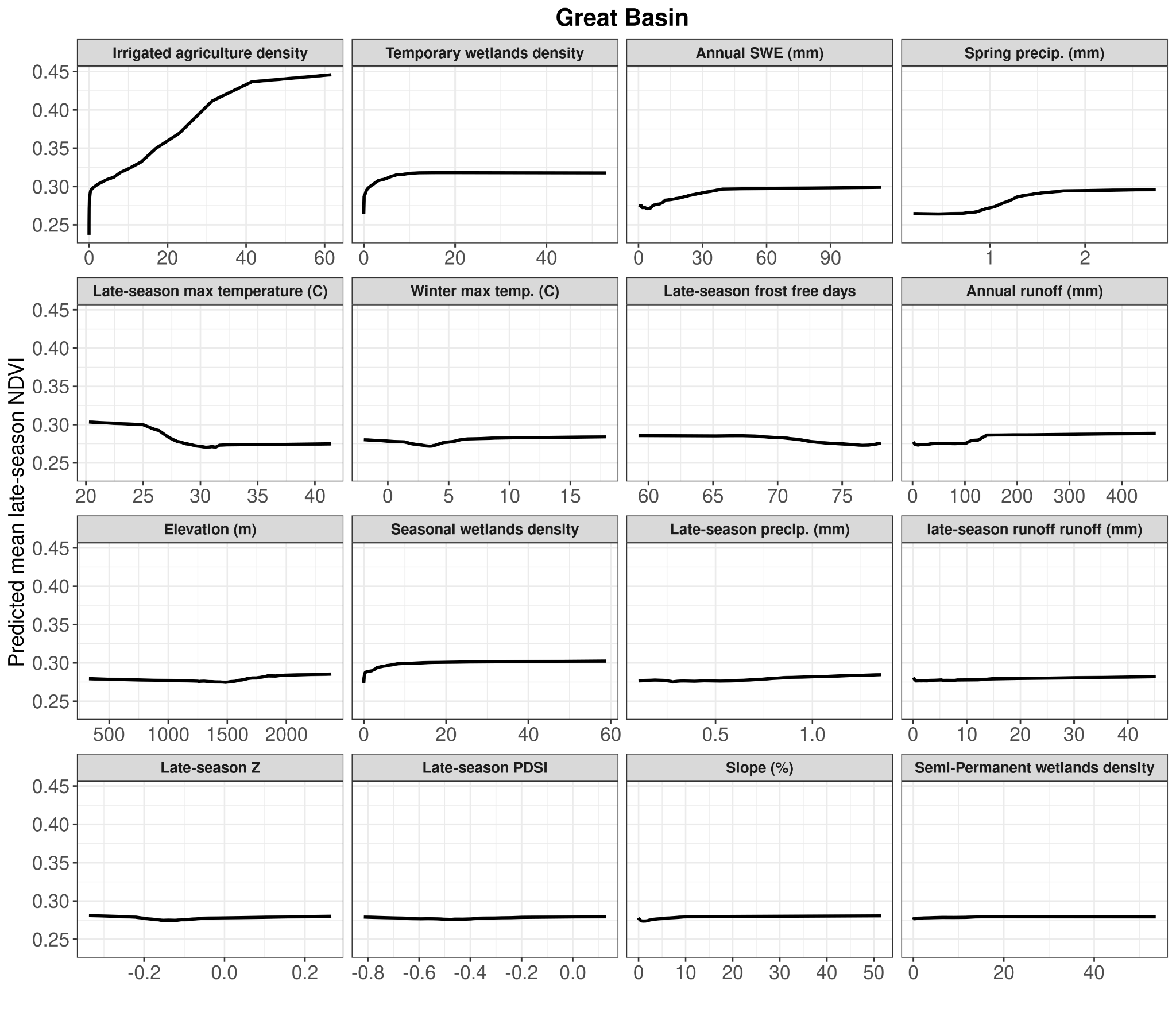

Figure B.24. Partial dependency plots for each variable used in the spatial model on predicting mesic resource productivity in the Great Basin region (1986-2023). Y-axis values are the predicted mean late-season NDVI. The X-axis shows untransformed variable values. Variables shown as densities are measured in hectares per 500-meter radius.

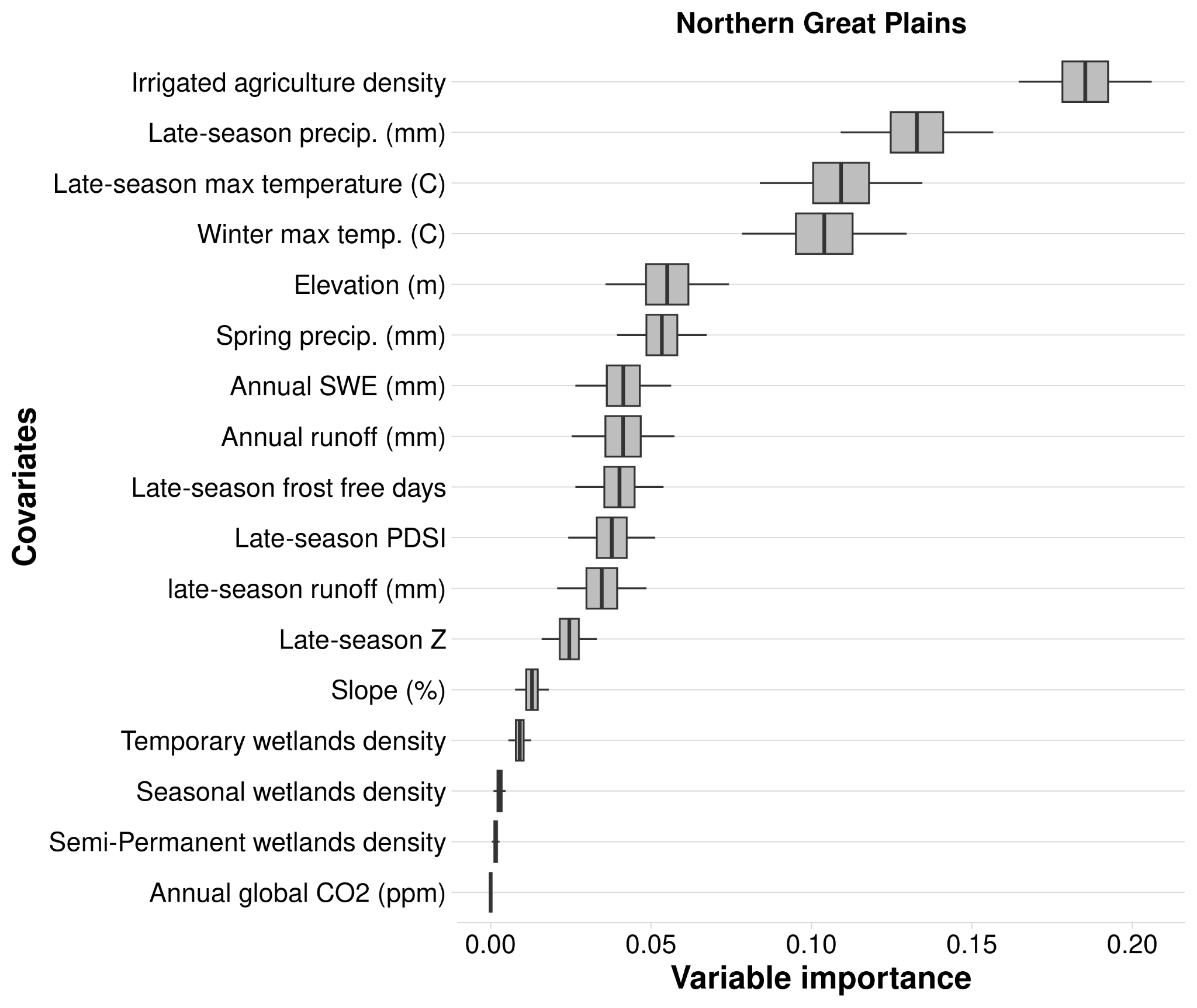

Figure B.25. Variable importance (VIMP) of variables predicting mesic resource productivity across the Northern Great Plains region from 1986 to 2023 using the spatial random forest analysis. Variable importance was assessed using the Breiman-Cutler permutation method, with higher values indicating greater predictive power. Center black lines represent median importance values from 100 subsampled VIMP scores; boxes cover the 25th to 75th percentiles; whiskers extend to 95% confidence intervals. Variable importance is standardized by dividing by the variance of Y. Variables shown as densities are measured in hectares per 500-meter radius.

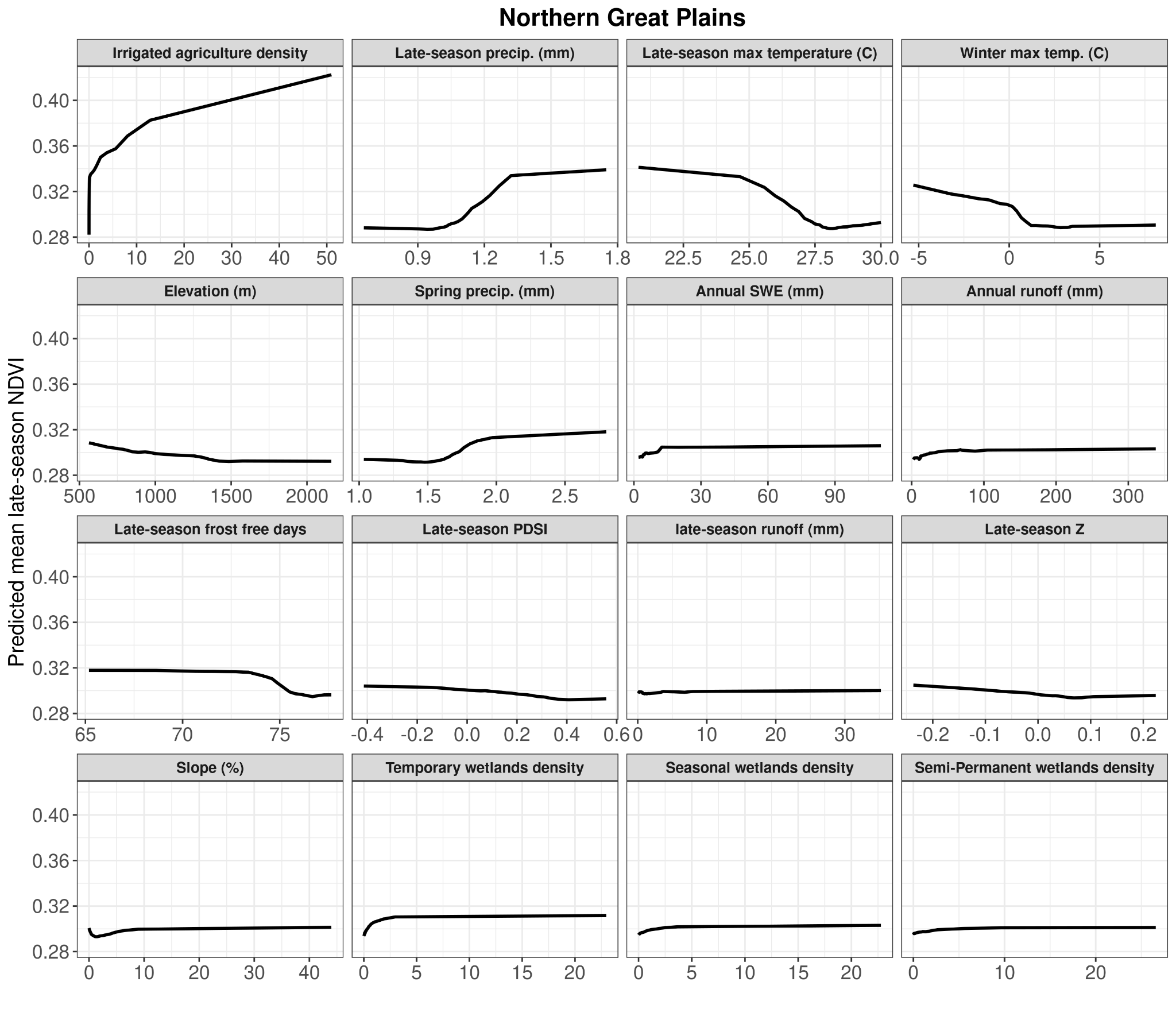

Figure B.26. Partial dependency plots for each variable used in the spatial model on predicting mesic resource productivity in the Northern Great Plains region (1986-2023). Y-axis values are the predicted mean late-season NDVI. The X-axis shows untransformed variable values. Variables shown as densities are measured in hectares per 500-meter radius.

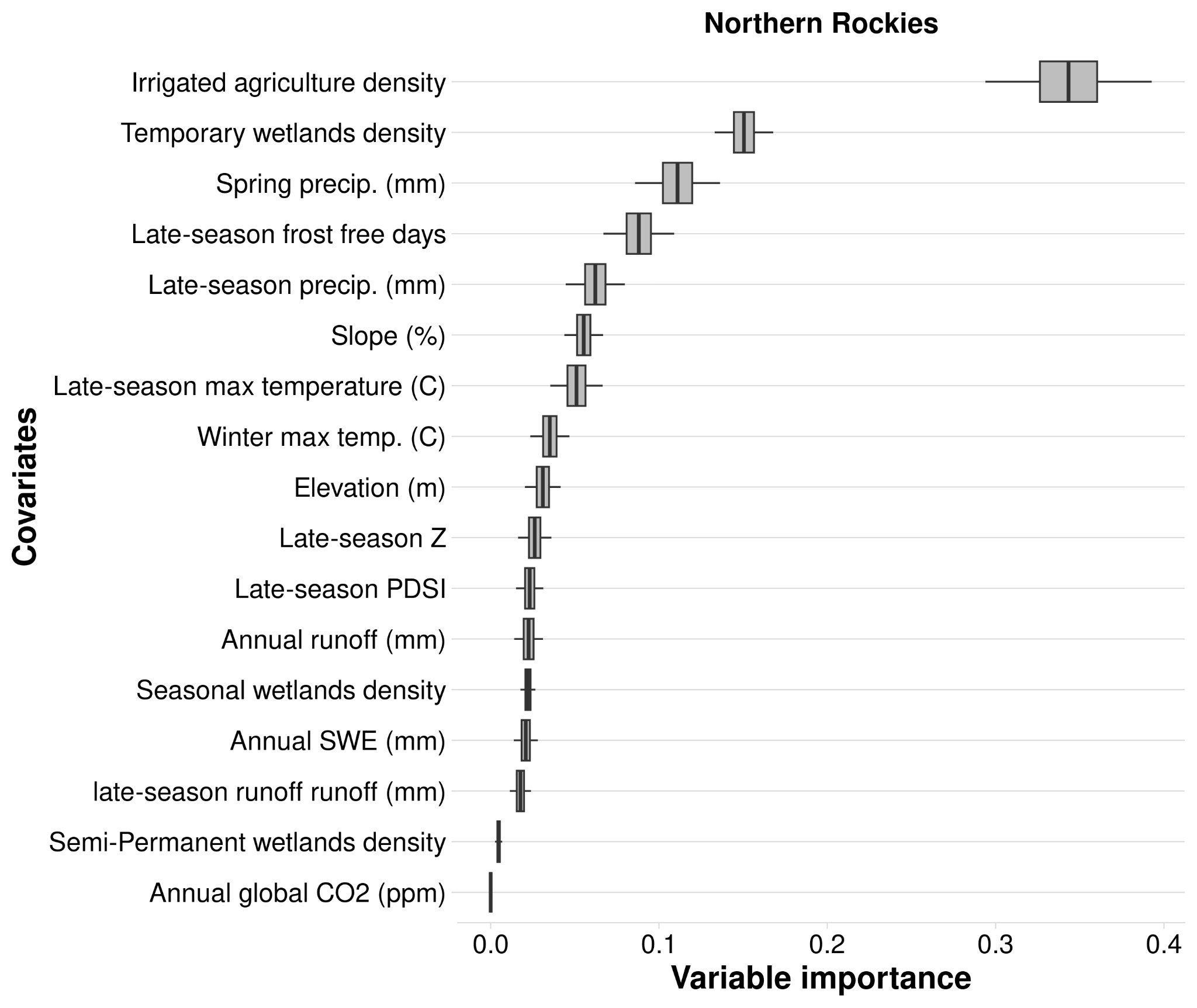

Figure B.27. Variable importance (VIMP) of variables predicting mesic resource productivity across the Northern Rockies region from 1986 to 2023 using the spatial random forest analysis. Variable importance was assessed using the Breiman-Cutler permutation method, with higher values indicating greater predictive power. Center black lines represent median importance values from 100 subsampled VIMP scores; boxes cover the 25th to 75th percentiles; whiskers extend to 95% confidence intervals. Variable importance is standardized by dividing by the variance of Y. Variables shown as densities are measured in hectares per 500-meter radius.

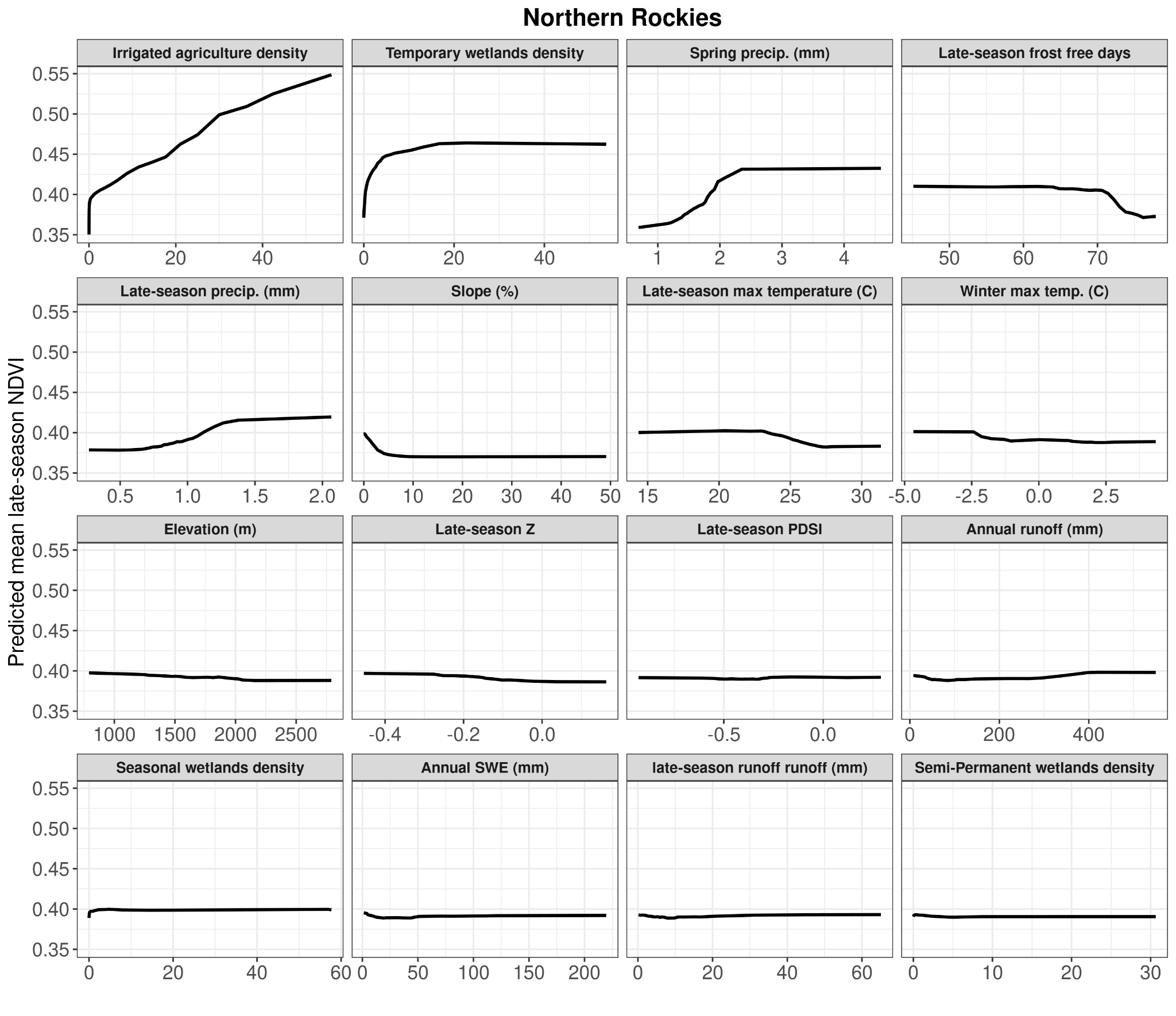

Figure B.28. Partial dependency plots for each variable used in the spatial model on predicting mesic resource productivity in the Northern Rockies region (1986-2023). Y-axis values are the predicted mean late-season NDVI. The X-axis shows untransformed variable values. Variables shown as densities are measured in hectares per 500-meter radius.

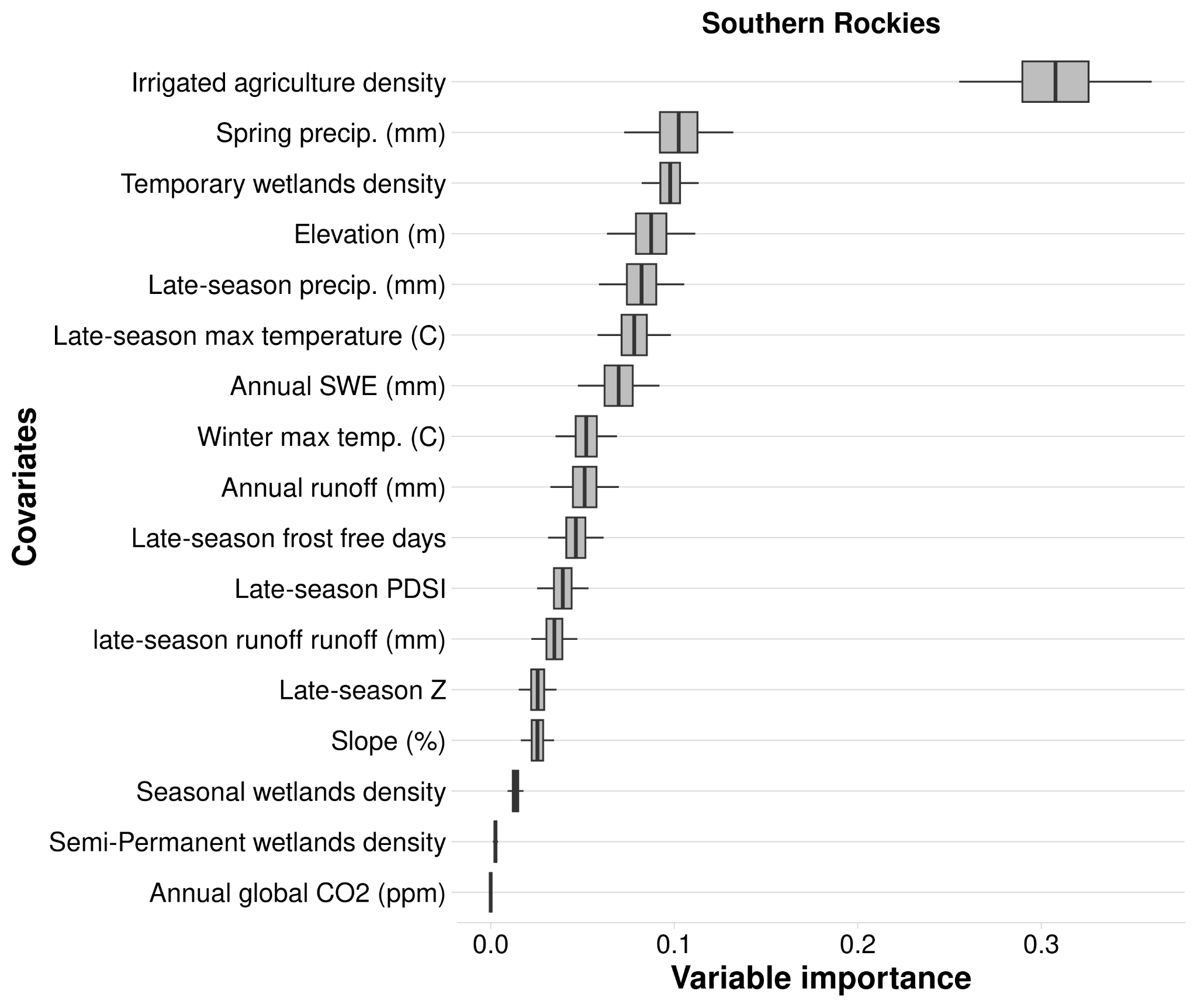

Figure B.29. Variable importance (VIMP) of variables predicting mesic resource productivity across the

Southern Rockies region from 1986 to 2023 using the spatial random forest analysis. Variable importance was assessed using the Breiman-Cutler permutation method, with higher values indicating greater predictive power. Center black lines represent median importance values from 100 subsampled VIMP scores; boxes cover the 25th to 75th percentiles; whiskers extend to 95% confidence intervals. Variable importance is standardized by dividing by the variance of Y. Variables shown as densities are measured in hectares per 500-meter radius.

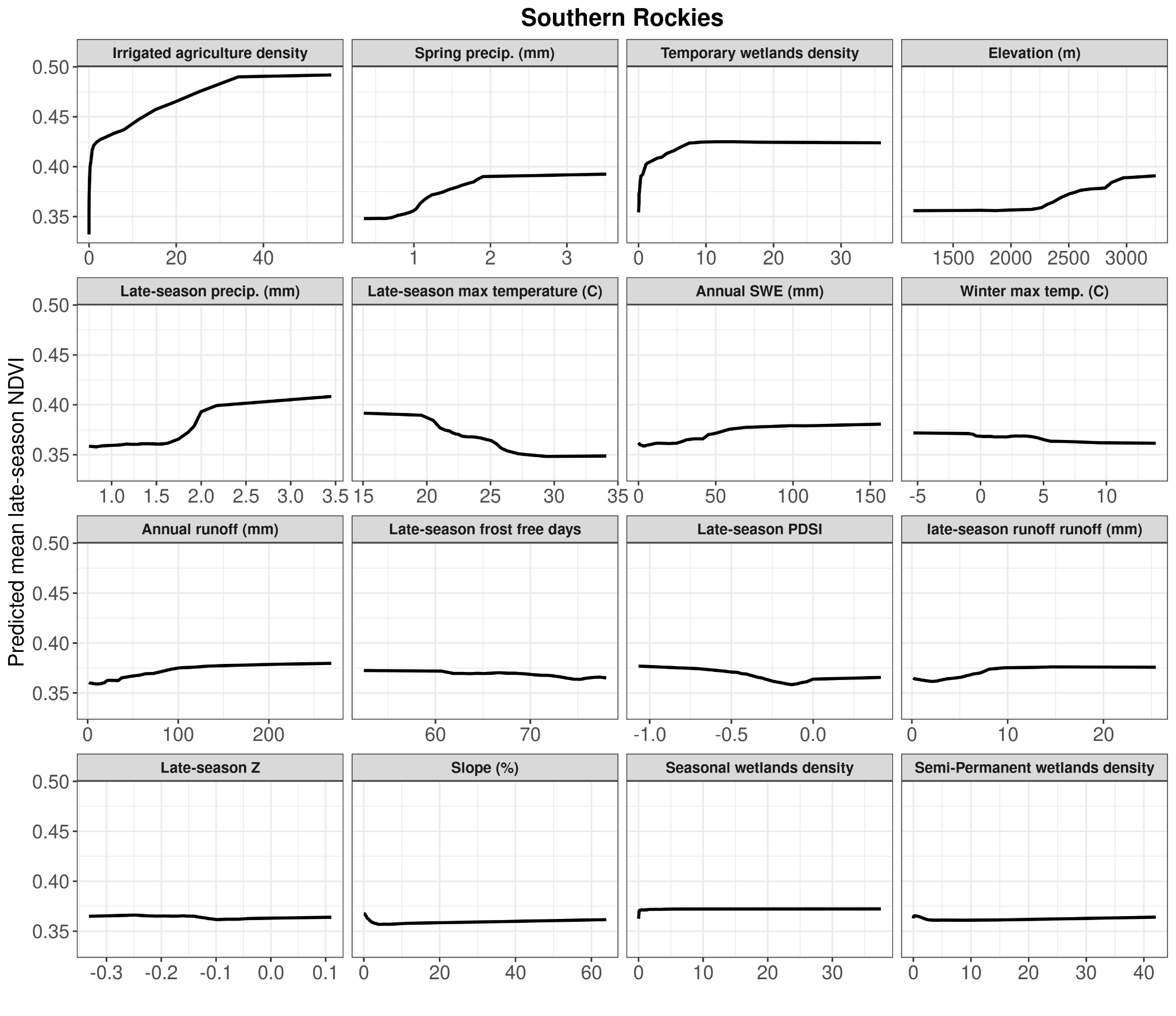

Figure B.30. Partial dependency plots for each variable used in the spatial model on predicting mesic resource productivity in the Southern Rockies region (1986-2023). Y-axis values are the predicted mean late-season NDVI. The X-axis shows untransformed variable values. Variables shown as densities are measured in hectares per 500-meter radius.

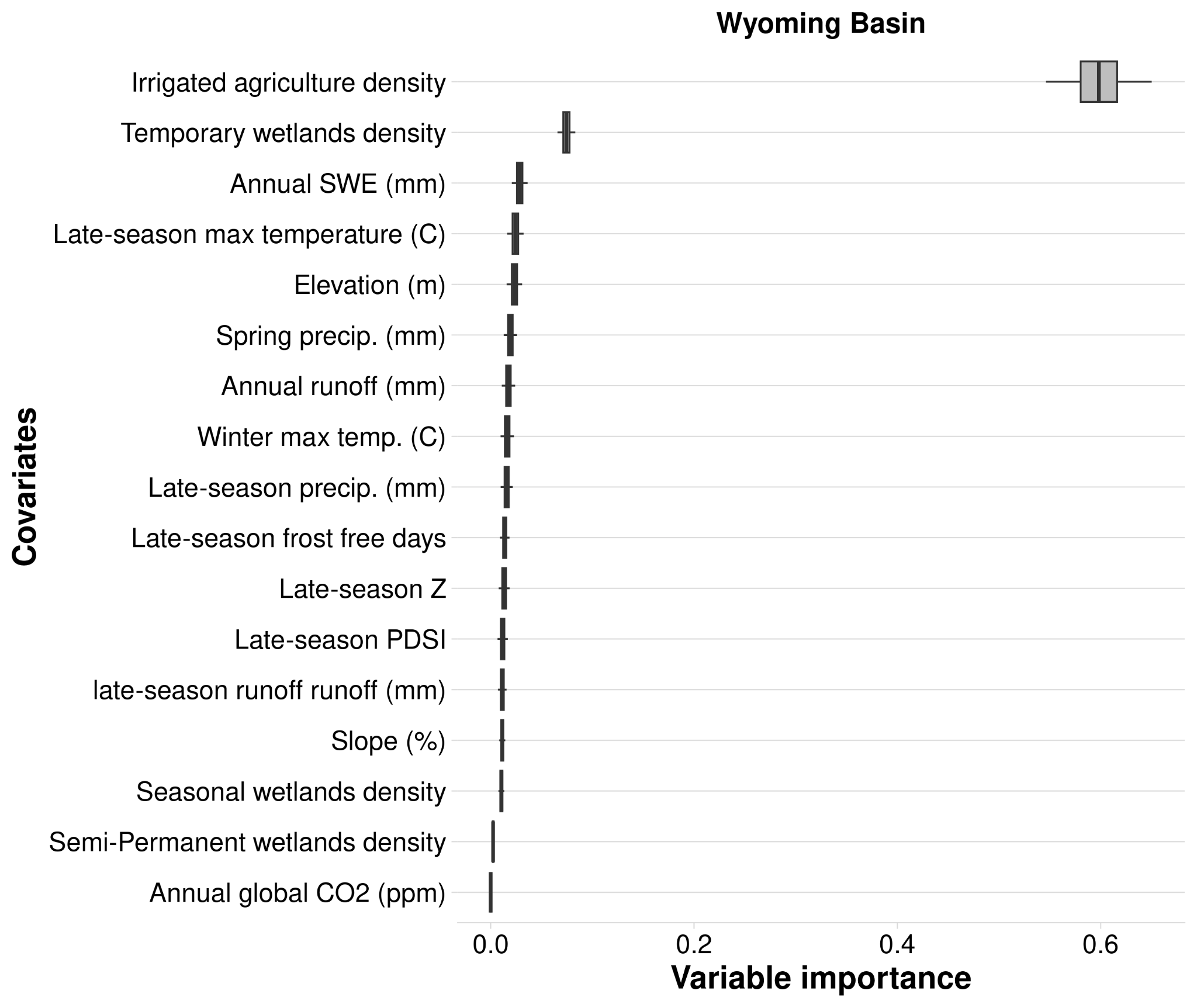

Figure B.31. Variable importance (VIMP) of variables predicting mesic resource productivity across the

Wyoming Basin region from 1986 to 2023 using the spatial random forest analysis. Variable importance was assessed using the Breiman-Cutler permutation method, with higher values indicating greater predictive power. Center black lines represent median importance values from 100 subsampled VIMP scores; boxes cover the 25th to 75th percentiles; whiskers extend to 95% confidence intervals. Variable importance is standardized by dividing by the variance of Y. Variables shown as densities are measured in hectares per 500-meter radius.

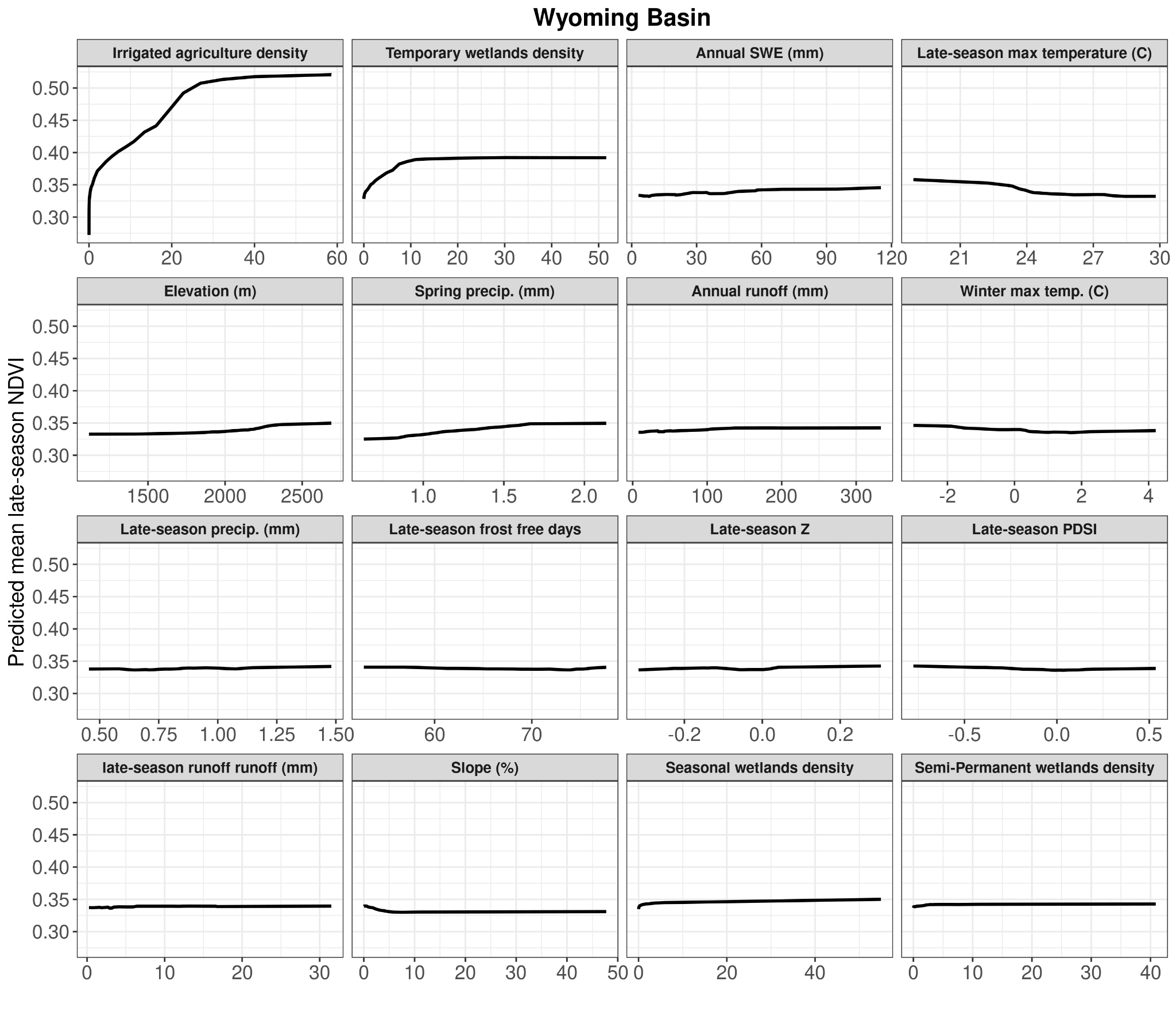

Figure B.32. Partial dependency plots for each variable used in the spatial model on predicting mesic resource productivity in the Wyoming Basin region (1986-2023). Y-axis values are the predicted mean late-season NDVI. The X-axis shows untransformed variable values. Variables shown as densities are measured in hectares per 500-meter radius.
